# Biosynthesis of peptidic thiooxazole metallophores installed by multinuclear nonheme iron enzymes

**DOI:** 10.1101/2025.11.18.689057

**Authors:** Mayuresh G. Gadgil, Shravan R. Dommaraju, Xiaopeng Liu, Alexander J. Battiste, Miriam H. Bregman, Douglas A. Mitchell

## Abstract

Significant effort has been directed toward characterization of nonheme iron enzymes owing to their breadth of unique reactivity. Through genome mining, we identified a conserved biosynthetic gene cluster within Pseudomonadota encoding one such family, the multinuclear nonheme iron-dependent oxidative enzymes (MNIO, formerly DUF692). Using a representative gene cluster from *Fontimonas thermophila*, we heterologously produced the post-translationally modified peptide fontiphorin, and detailed spectral analysis revealed MNIO-catalyzed installation of seven 5-thiooxazole (5TO) moieties. During our work, additional MNIO products were reported with conflicting structural assignments, so we investigated the related biosynthetic gene clusters from *Haemophilus influenzae* and *Neisseria gonorrhoeae*. Using alkylation-assisted HMBC correlations, we demonstrated that these products also contain 5TO resulting in a revision of the structure of oxazolin. We further provide evidence supporting a role for 5TO-containing peptides in copper detoxification and recommended this emerging class of <u>C</u>u-<u>a</u>ssociated <u>p</u>eptidic thio<u>o</u>xazole metallophores be referred to as captophorins. To further explore the captophorins, we reconstituted fontiphorin biosynthesis *in vitro* and investigated its enzymatic requirements. Using cell-free production of single-site, double-site, and naturally occurring variants, we examined enzyme-substrate interactions to determine key sites governing catalysis by 5TO-forming MNIOs. Through our detailed spectroscopic approach for 5TO assignment and investigation of enzyme-substrate interactions, our work unifies tens of thousands of MNIOs in the biosynthesis of captophorins.

## Introduction

Ribosomally synthesized and post-translationally modified peptides (RiPPs) comprise a large family of natural products whose activities are derived from enzymatically installed chemical modifications on gene-encoded precursor peptides.^1,2^ Canonical RiPP biosynthesis involves: *(i)* translation of the precursor peptide; *(ii)* substrate recognition by pathway-specific proteins; and *(iii)* enzymatic installation of post-translational modifications (PTMs; **Figure 1**). RiPPs exhibit diverse biological activities, including membrane disruption, transcription/translation inhibition, quorum sensing, and metal scavenging.^3^ To serve these myriad roles, RiPPs often receive unique PTMs, and many enzyme families have been discovered which perform unprecedented chemistry on RiPP substrates. In recent years, the multinuclear non-heme iron-dependent oxidative enzyme family (MNIO, formerly DUF692) has received increasing attention in the RiPP field.^4–11^

**Figure 1.**
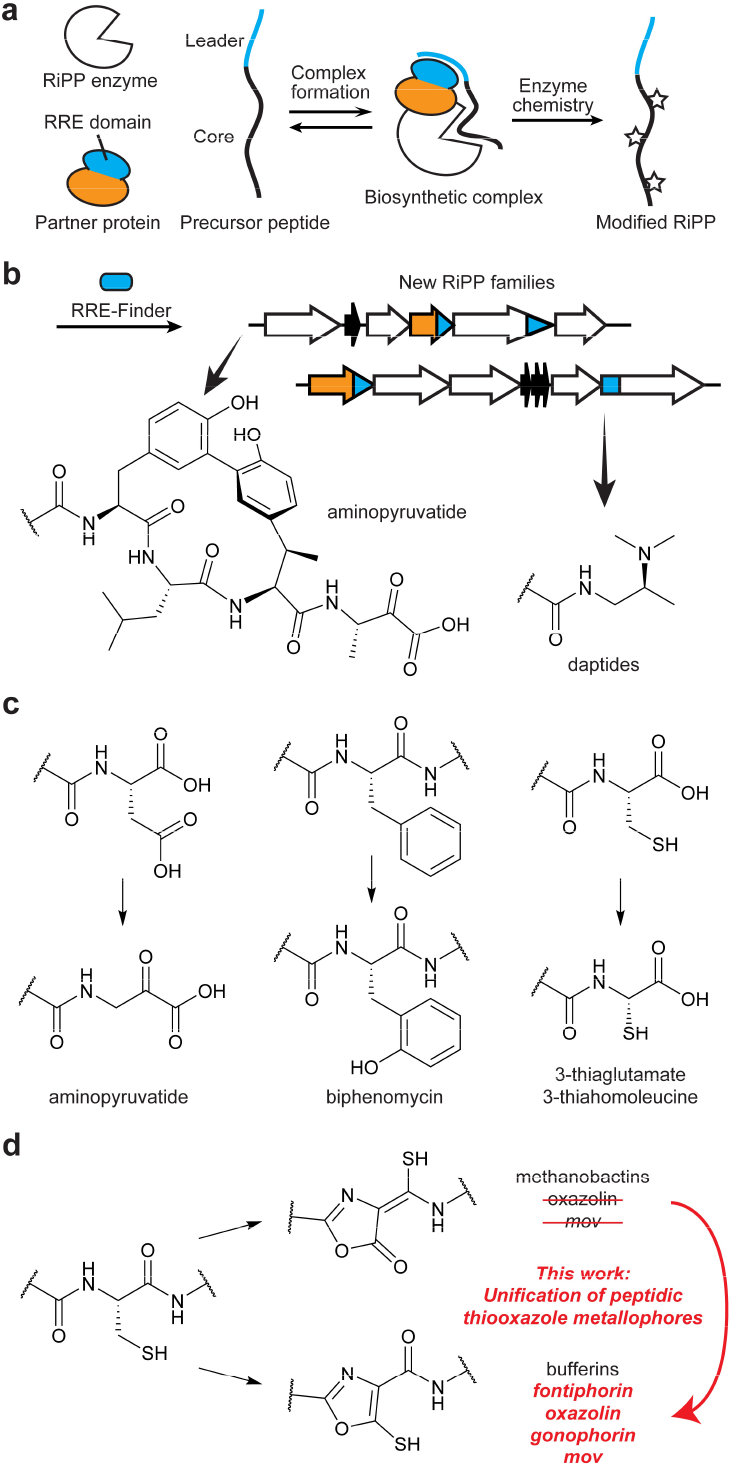
RRE-dependent biosynthesis by MNIOs. **a**, Scheme for RiPP biosynthesis with RRE-containing partner proteins. **b**, Examples of chemical transformations discovered using RRE-Finder. **c**, Example reactions catalyzed by RRE-dependent MNIOs. **d**, Oxazolone/thioamide (top) and 5-thiooxazole (bottom) PTMs installed by MNIOs.

MNIO enzymes use molecular oxygen to oxidatively reconfigure peptide substrates, often at cysteine residues.^9^ They catalyze a broad range of chemistries, including heterocycle formation, carbon excision, macrocyclization, C-N bond cleavage, and more. MNIOs adopt a triose-phosphate isomerase barrel fold with a metal-binding site accommodating multiple iron atoms. The available spectroscopic data suggest that MNIOs use mixed-valent Fe(III)–superoxo and Fe(IV)–oxo states for catalysis. While currently theoretical, mechanistic models postulate anchoring of a Cys thiolate to one iron while a neighboring ferrous site binds O_2_ to generate the oxidizing equivalents required for catalysis. The MNIO landscape contains thousands of phylogenetically diverse members which remain to be functionally characterized.

One common feature of MNIO biosynthetic gene clusters (BGCs) is the use of partner proteins, which often contain a RiPP Recognition Element (RRE; **Figure 1**).^9,12^ The RRE is the most common protein domain across the ∼50 described molecular classes of RiPPs, and examination of RREs can facilitate identification of new RiPP pathways. By systematically mining RRE domains using RRE-Finder,^13^ we have characterized multiple RiPPs containing new chemical transformations installed by various enzyme families (**Figure 1**).^8,14,15^

Through mining of RREs, we identified thousands of MNIOs putatively involved in heavy metal and oxidative stress response. To investigate this conserved gene cassette, we selected the *fon* BGC from *Fontimonas thermophila* and characterized its product, fontiphorin. Through rigorous spectrometric and spectroscopic techniques, we demonstrated that fontiphorin contains seven 5-thiooxazole (5TO) moieties. During our studies, multiple related biosynthetic gene clusters were reported with conflicting structural assignment of the products,^7,16,17^ and we theorized that some of these structures were possibly misassigned. Thus, we obtained spectroscopic data for one of these products, oxazolin, and a closely related new product from *Neisseria gonorrheae*, gonophorin. Our data conclusively demonstrate 5TO formation in both cases. We further affirm that fontiphorin binds copper, consistent with literature precedents and connect these MNIOs in biosynthesis of 5TO. Thus, we propose the name captophorins for this class of <u>C</u>u-<u>a</u>ssociated <u>p</u>eptidic thio<u>o</u>xazole metallophores. Following *in vitro* reconstitution of FonBC, we used cell-free approaches to investigate the substrate requirements for captophorin biosynthesis and identify factors controlling substrate specificity. Altogether, we unify the largest groups of MNIO enzymes by characterizing the spectroscopic features and biosynthetic requirements for captophorins.

## Results and Discussion

### Bioinformatic profiling of RRE-dependent MNIOs

The use of RRE-Finder has allowed rapid profiling of unexplored RiPP BGCs.^13^ We systematically analyzed data produced by one of our previous bioinformatic campaigns, which employed a divisive hierarchical clustering approach to sort RRE domains into BGC families.^14^ To do this, sequence logos for precursor peptides within each putative family were generated and local genomic context was analyzed for gene co-occurrence (**Supplementary Note, Figure S1**).^18,19^ Among the results of this procedure were BGC families with the following features: *(i)* MNIO enzyme; *(ii*) a PME1^20^ partner protein containing a DUF2063 domain and a predicted RRE (PME1, Type 1 <u>P</u>artner protein of <u>M</u>NIO <u>E</u>nzyme); and *(iii)* a Cys-rich precursor peptide. To explore this further, we sought to genomically characterize the global set of PME1-dependent MNIOs.

We gathered a comprehensive set of PME1 proteins in NCBI (National Center for Biotechnology Information) and analyzed their local genomic contexts using RODEO.^21^ Through these efforts, >24,000 genetic loci encoding both a MNIO and PME1 were identified (**Dataset S1**). We generated a sequence similarity network (SSN) for the MNIO family and identified all proteins with a local PME1, revealing that the majority of MNIO proteins co-occur with a PME1 (70.1%, **Figure S2**). MNIO-PME1 pairs represented the two largest groups; however, most characterized MNIO proteins occurred outside of these groups.^9,20^ We subsequently examined the converse relationship and showed that PME1s are almost universally encoded next to MNIO proteins, supporting their assignment as dedicated MNIO partner proteins (**Figure S3**). Among our dataset were proteins with solved structures, PDB codes 3BWW (MNIO, *Histophilus somni*) and 3DEE (PME1, *N. gonorrhoeae*; **Figures 2, S4**).^22^ Examination of their local genomic regions suggested both are involved in *bona fide* RiPP biosynthesis. Further analysis identified frequent co-occurrence of DoxX, DUF2282, and transcriptional regulators (**Table S1**). We also identified three general precursor peptide categories: *(i)* an identified HMM match for DUF2282; *(ii)* a 5-mer repeat containing a Lys-Cys motif; or *(iii)* multiple Cys-Lys motifs (**Dataset S2, Table S2**).

**Figure 2.**
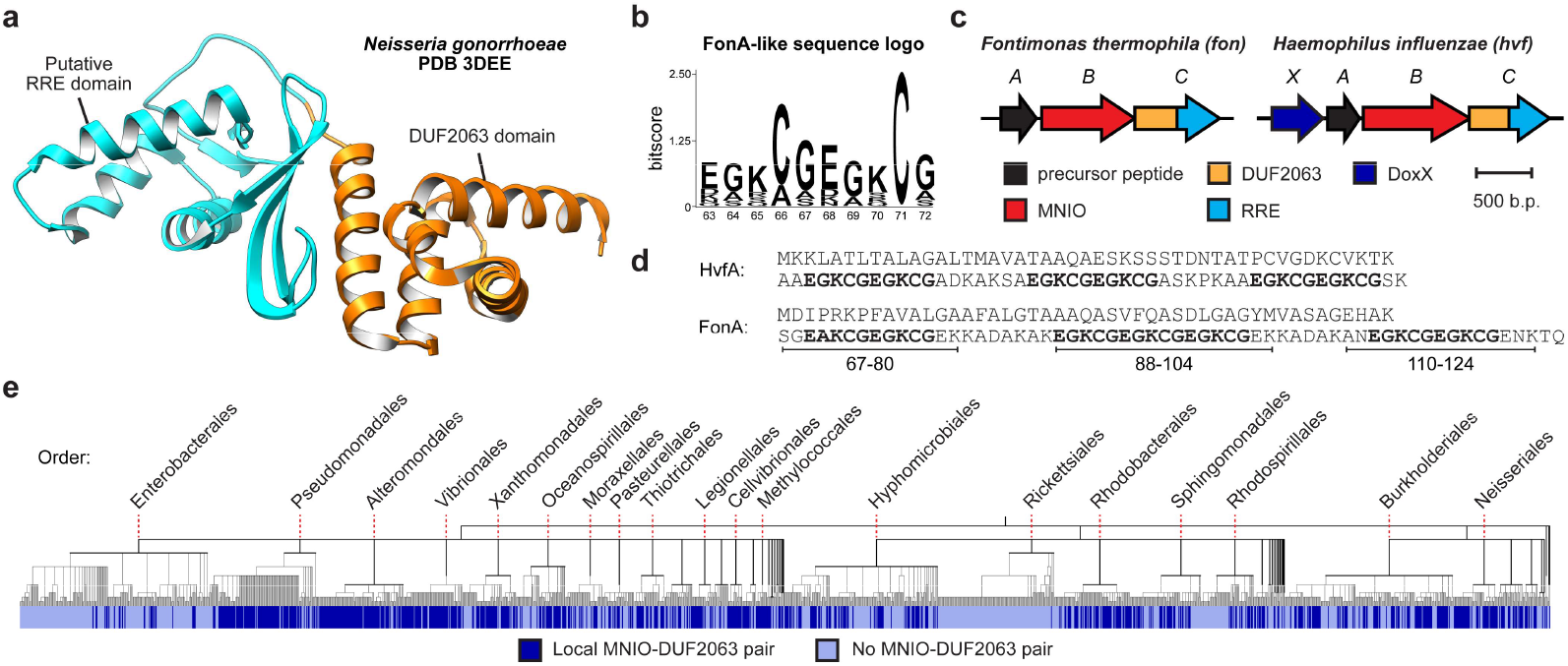
Bioinformatic identification of MNIO-PME1 BGCs. **a**, DUF2063-domain containing PME1 from *N. gonorrhoeae* (PDB code: 3DEE). **b**, Sequence logo (*n* = 100) for the EGKCG repeat motif identified within FonA-like sequences. **c**, Fontiphorin and oxazolin BGC diagrams. **d**, Amino acid sequences for FonA (fontiphorin precursor peptide) and HvfA (oxazolin precursor peptide). **e**, Distribution of MNIO-PME1 BGCs across Pseudomonadota NCBI reference genomes. Phylogenetic tree (*n* = 1,919) was generated from NCBI taxonomy data using phyloT and visualized with iTOL.

To explore the possible function(s) of these BGCs, we performed an extensive literature review.^23–30^ We discovered evidence connecting MNIO-PME1 BGCs to oxidative stress and metal response, so the distribution of these BGCs across bacterial taxa was investigated further (**Table S3**). While MNIOs are taxonomically diverse, the majority are encoded by Pseudomonadota. After gathering all Pseudomonadota reference genomes from NCBI, we searched for MNIOs and PME1 proteins occurring within the same locus and annotated a phylogenetic tree accordingly (**Figure 2**). MNIO-PME1 BGCs are distributed widely across Pseudomonadota, occurring in about half of reference genomes. They are depleted in certain bacterial orders (*e*.*g*., Enterobacterales, Rickettsiales) and enriched in others (*e*.*g*., Pseudomonadales, Vibrionales, Neisseriales). Additionally, MNIO-PME1 BGCs were identified in multiple human pathogenic organisms, including WHO (World Health Organization) bacterial priority pathogens *Pseudomonas aeruginosa, Neisseria gonorrhoeae*, and *Haemophilus influenzae*.^31^ Altogether, MNIO-PME1 BGCs are much more prevalent relative to other secondary metabolic pathways and likely play critical stress response roles across Pseudomonadota.

### Expression of modified fontiphorin

To begin our experimental investigation, we selected the *fon* BGC from *F. thermophila*, a moderately thermophilic Gammaproteobacterium isolated from a hot spring (**Figure 2**). Thermophilic organisms have been a consistent source of RiPP biosynthetic enzymes amenable for biochemical characterization.^15,32^ The *fon* BGC also represented a minimal case for exploration, containing only the predicted MNIO, PME1, and precursor peptide (**Table S4**). We obtained *E. coli* codon-optimized genes for *fonABC* and introduced a His_6_-tag before *fonA* (**Tables S5-7**). Cultures were supplemented with (NH_4_)_2_Fe(SO_4_)_2_, and products were purified by immobilized metal affinity chromatography (IMAC). Products of FonABC co-expression displayed unique local absorbance features at 254, 302, and 600 nm and were green upon visual inspection (**Figure S5**). Liquid-chromatography-mass spectrometry (LCMS) analysis showed a loss of ∼30 Da (**Figure S6**). Given the size of FonA, we next digested the sample with endoproteinase LysC. Using high-resolution mass spectrometry (HRMS), we detected peptides corresponding to FonA_67-80_, FonA_88-104_ and FonA_110-124_. FonA_67-80_ ([M+3H]^3+^: obs. *m/z*, 458.8502; exp. *m/z*, 458.8519; ppm error, −3.7) and FonA_110-124_ ([M+3H]^3+^: obs. *m/z*, 505.8681; exp. *m/z*, 505.8699; ppm error, −3.6) each displayed a loss of 8 H atoms, while FonA_88-104_ ([M+3H]^3+^: obs. *m/z*, 562.2085; exp. *m/z*, 562.2098; ppm error, −2.5) displayed a loss of 12 H atoms. This pattern of 4 H atoms lost per repeat suggested each repeat was similarly modified, and analysis of FonA_67-80_ using MALDI-LIFT-MS localized a 4 Da mass loss to each Lys-Cys motif (**Figure S7**).

Next, FonA_67-80_ was purified for detailed structural characterization. Purification required use of a polar group-functionalized C18 stationary phase, as we observed no retention using a standard C18 stationary phase. During purification, FonA_67-80_ displayed a strong UV absorption at 302 nm and was prone to aggregation (**Figure 3**). We hypothesized that any intact thiol groups in the modified structure may cause aggregation due to inter- and/or intra-molecular disulfide formation. Thus, we reacted the peptide with iodoacetamide (IAA) to alkylate free thiol with MALDI-TOF-MS analysis confirming two alkylation events (**Figure 3**). Upon purification of the alkylated FonA_67-80_, the local absorbance maximum at 302 nm was significantly diminished. High resolution tandem mass spectrometry (HRMS/MS) showed both Lys-Cys motifs had been alkylated, and that modification prevented backbone fragmentation (**Figure S8, Table S8**).

**Figure 3.**
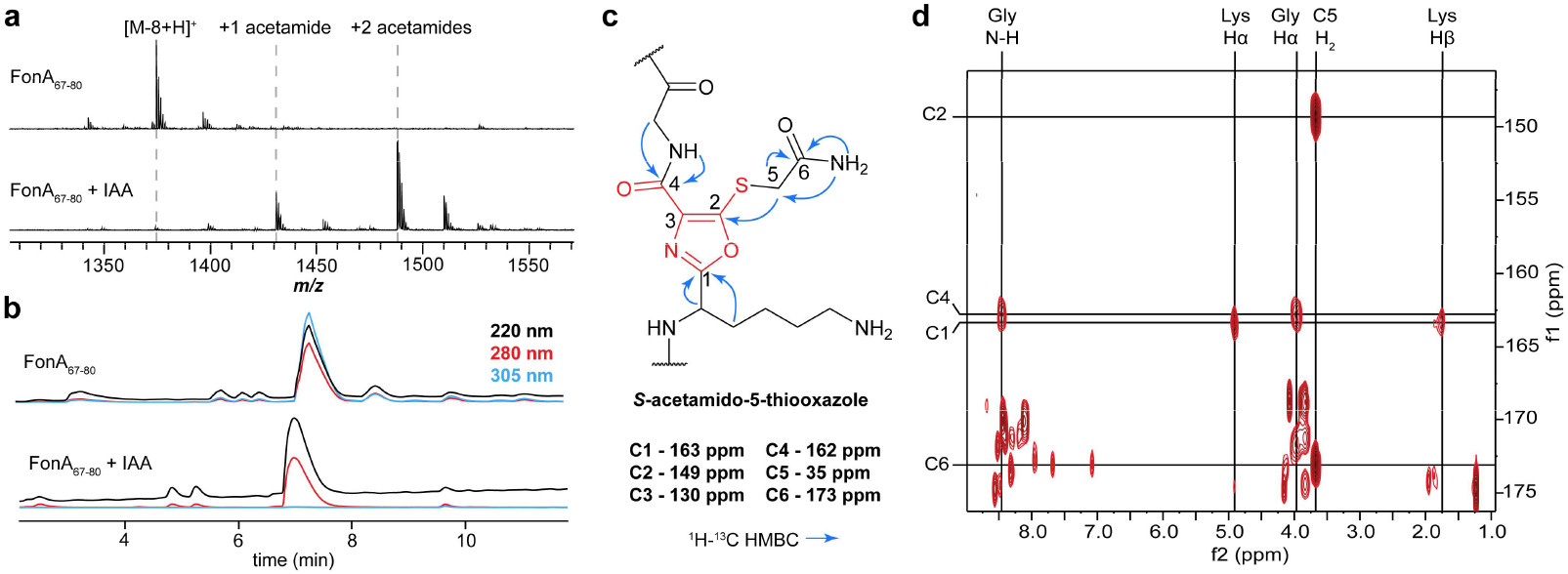
Structural elucidation for 5TO in fontiphorin. **a**, MALDI-TOF-MS spectra for FonA_67-80_ (SGEAKCGEGKCGEK) before and after alkylation by IAA. **b**, HPLC chromatograms for FonA_67-80_ before and after alkylation by IAA. Multiple wavelengths are overlaid for each analyte. **c**, Structural assignment for *S*-acetamido-5-thiooxazole with key HMBC correlations denoted. **d**, ^1^H-^13^C HMBC for alkylated FonA_67-80_ showing key differential correlations for assignment of 5TO. Abbreviation: IAA, Iodoacetamide.

### Spectroscopic evidence for 5-thiooxazoles in FonA_**67-80**_

NMR spectroscopy was performed to determine the chemical structure of alkylated FonA_67-80_ (**Figure S9**). We conducted a suite of 1D- and 2D-NMR experiments, including ^13^C, ^1^H-^1^H TOCSY, ^1^H-^13^C HSQC, and ^1^H-^13^C HMBC. Using these data, the spin systems for Ser, Gly, Glu, and Ala residues were first assigned (**Figure S10, Table S9**). From the TOCSY spectrum, it was determined that a multiplet at ∼5 ppm comprised two overlapping Lys Hα signals, with correlations observed to all canonical Lys side chain protons (**Figures S11-S12**). A third Lys residue was also identified, which was deduced to be unmodified Lys_80_. We attributed the downfield shift of the Lys_71/76_ Hα to deshielding effects from the modification. In the ^13^C spectrum, four signals at 130, 150, 162, and 163 ppm were observed which did not appear in the ^1^H-^13^C HSQC spectrum. Given the peptide sequence, we surmised this could only arise from installation of a proton-deficient heterocycle.

The ^1^H-^13^C HMBC data was examined to investigate this hypothesis and ultimately allowed us to assign the modification as 5TO. Correlations were observed from the modified Lys_71/76_ Hα and Hβ to the ^13^C signal at 163 ppm, showing connectivity between Lys and the 5TO C1. We further identified correlations connecting the acetamide CONH_2_ (173 ppm) and CH_2_ (35 ppm). From the acetamide CH_2_, analysis of the ^1^H-^13^C HMBC data identified a correlation to the unknown ^13^C signal at 150 ppm (**Figure 3**). Given the selective reactivity of IAA, we assigned this correlation as a ^3^J coupling travelling through the S atom to 5TO C2. Finally, the Gly_73/78_ N-H and Hα showed ^1^H-^13^C HMBC correlations to the ^13^C signal at 162 ppm, allowing assignment to 5TO C4.

Using the substructures above, we assembled putative heterocycles. As all proton signals had been assigned to non-Cys residues, we hypothesized that loss of Cys N-H, Hα, and Hβ resulted in the −4 Da mass shift of the modification. To account for the downfield ^13^C signal and loss of ^1^H-^13^C HSQC correlation at 5TO C2, we deduced that this may require a desaturation and a second heteroatom bond. Applying an α,β desaturation and bond formation from the Lys O atom to C2 would account for the lack of fragmentation, loss of HSQC correlations, downfield ^13^C chemical shift, and distinct ^1^H-^13^C HMBC correlations. To satisfy atomic valences and the missing Cys N-H, we arrived at 5TO (**Figures 3, S13**). Thus, the remaining ^13^C signal at 130 ppm was assigned to C3 of 5TO. Additionally, the 5TO C4 (162 ppm; formerly Cys C1) is retained, supported by fragmentation between 5TO and Gly in the peptide. While other heterocyclic ring structures were considered (*e*.*g*., oxazolone/thioamide [*vide infra*], azirine, oxazinone),^33^ 5TO was the only structure satisfying all lines of spectroscopic evidence.

### MNIO-PME1 BGCs broadly install 5TO

During our studies, multiple new MNIO products were reported.^7,16,17^ Bufferin 1 from *Caulobacter vibrioides* was assigned with 5TO moieties, while MovA and oxazolin were assigned with oxazolone/thioamide groups (OxT, **Figure S14**). Both structures are identical except for an exchange of O and S atom positions. As a result, differentiation of these two structures represents a significant challenge, and the proton-deficient ring systems obfuscated definitive NMR-based evidence. We examined the similarity of FonBC with characterized MNIOs and found that HvfB, BufB_1_, BufB_2_, and FonB are closely related and are encoded near PME1 proteins (**Figure S15**). MbnB and MovB are more distantly related and do not encode nearby PME1 proteins. Additionally, oxazolin and fontiphorin both contain EGKCG repeats (**Figure 2**). Altogether, we theorized that one or more of these structures could be misassigned.

We first re-examined our assignment of 5TO. The OxT moiety is well-precedented in MNIO-catalyzed biosynthesis of methanobactins, which have been characterized by UV, NMR, X-ray crystallography, chemical degradation, and comparison to synthetic standards.^34–36^ We did not identify reports of OxT susceptibility to IAA alkylation, but fontiphorin was readily alkylated. While methanobactins display absorbance maxima at ∼340 and ∼390 nm, oxazolin, bufferin 1, and MovA all displayed similar absorbance maxima to fontiphorin at ∼305 nm (**Figure S14**). Analysis of the same compounds revealed a lack of correlations within the putative heterocycles. To obtain our NMR data, we used the *S*-alkylated product, which provided critical HMBC correlations from the *S*-acetamide group to 5TO C2 and from the subsequent Gly residue to 5TO C4. The presence of OxT would result in correlations to the same ^13^C signal; therefore, FonA cannot possess OxT (**Figure 4**).

**Figure 4.**
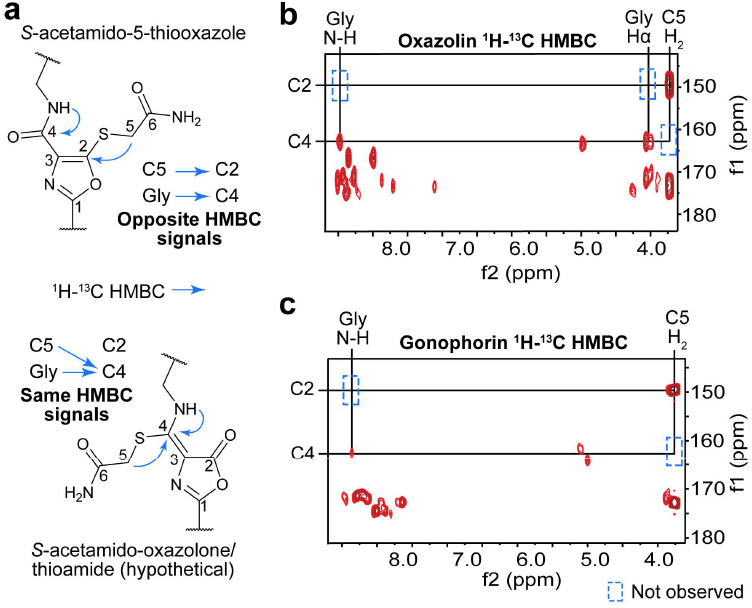
Spectroscopic evidence for 5TO formation in oxazolin and gonophorin. **a**, Structure for alkylated 5TO and hypothetical structure for alkylated OxT. ^1^H-^13^C HMBC for alkylated products of **b**, *hvf* and **c**, *ngo* focusing on Gly and *S*-acetamide ^1^H-^13^C HMBC correlations.

While the *fon* and *hvf* BGCs are highly similar, it remained possible that MNIO-PME1 BGCs could install multiple different PTMs. Thus, we investigated if alkylation-assisted HMBC correlations could resolve any structural discrepancies. *E. coli* codon-optimized genes for *hvfABC* were obtained, and the BGC was expressed (**Tables S4-S7**). Following IMAC purification of oxazolin, we noted the same λ_max_ at ∼305 nm and treated the peptide with IAA (**Figure S16**). Following digestion by trypsin, we purified peptide fragments by HPLC. MS analysis of the peptide fragments revealed each EGKCG repeat had been fully modified and alkylated (**Figure S16**). Following acquisition of 2D-NMR spectra for the modified peptides (**Figures 4, S17**), analysis of the ^1^H-^13^C HMBC data revealed that the *S*-acetamide groups and Gly residues were connected to distinct carbons at 150 ppm and 163 ppm, respectively. These data are incongruent with OxT and show that oxazolin possesses 5TO.

As additional evidence, we selected the *ngo* BGC from *N. gonorrhoeae*, from which the putative PME1 protein, NgoC, had been previously crystallized (PDB code: 3DEE; **Figures S4, S15**). After acquiring the requisite *ngo* genes, we repeated the procedure above, and we observed the same λ_max_ of ∼305 nm (**Figure S18**). Following IAA alkylation and purification, we collected NMR data for the full-length product (**Figures 4, S18-S19**). Once again, nearly identical ^1^H and ^1^H-^13^C HMBC signals were detected to those we had observed for fontiphorin and oxazolin. Once again, these data are incompatible with OxT formation and show the *ngo* product, gonophorin, also possesses 5TO groups.

The similar structures of 5TO and OxT render definitive NMR data difficult to obtain. While the reactivity of OxT moieties towards IAA is still unknown, 5TO readily undergoes alkylation. In the absence of high-resolution 3D diffraction data, the alkylation-assisted HMBC correlations reported here may represent the most reliable method for differentiating the two heterocycles (**Figure 4**). Additionally, we note the consistency of ∼305 nm absorbance maxima for 5TO containing compounds (fontiphorin, oxazolin, gonophorin, bufferins, modified SbtMa).^16,17,33^ While this is not unique to 5TO, confidently assigned methanobactins do not display this absorbance feature. Thus, UV spectral data is diagnostic for differentiating 5TO and OxT. While we did not experimentally characterize the *mov* product, its UV absorption profile supports 5TO rather than OxT.^7^ During the 2^nd^ International RiPPs Conference (October 2024), we publicly shared the alkylation-assisted HMBC approach to aid natural product researchers in assigning thiol-containing, proton-deficient heterocycles.

### Biological function of 5TO-containing peptides

A plethora of literature precedent suggested a role for MNIO-PME1 BGCs in oxidative stress and metal response. While methanobactin is induced by copper starvation, excess heavy metal was a frequent inducer of 5TO biosynthetic pathways.^23,25^ Accordingly, we suspected that 5TO metallophores could act in metal sequestration. Other reports have also recently shown copper binding functions for 5TO-containing peptides.^16,17,37^ Guided by these insights, we generated a co-expression vector containing FonBC and a maltose-binding protein (MBP)-fusion of FonA. Upon expression and purification of MBP-FonABC, we added Cu(II) in the presence of ascorbic acid to generate Cu(I) *in situ*. Following extensive buffer exchange, native MS analyses revealed binding of up to four Cu atoms, providing strong support for a role in Cu-binding (**Figure S20**).

Our attempts to purify Cu-loaded peptides for additional characterization faced significant challenges. We observed precipitation of fontiphorin upon *in vitro* loading with Cu(I) and a loss of chromatographic retention (**Figure S21**). We suspected that this behavior resulted from aggregation. Thus, we analyzed fontiphorin by dynamic light scattering, which showed Cu-induced formation of large-radius particles, most likely due to intermolecular coordination (**Figure S22**).^38^ Additionally, *in vivo* copper loading was investigated. We expressed *ngoABC* and added 50 μM CuSO_4_ 3 h post-induction. Immediately following IMAC purification and 500-fold buffer exchange, we analyzed the sample by LC-MS. Compared to a control, gonophorin expressed with supplemental CuSO_4_ was bound to either 1 or 2 Cu atoms (**Figure S23**); however, a complex mixture of species was also observed that was concomitant with precipitation and a time-dependent reduction of ion intensity.

Evidence from Leprevost et al.^16^ suggests that DUF2282 products bind Cu in an ordered fashion. In contrast, we suspect EGKCG products may bind Cu stochastically, which could explain the above results. We speculate that EGKCG peptides form a heterogeneous Cu-binding network, and that fully Cu-bound peptides aggregate over time. While we do not know if this behavior is biologically relevant, it plausibly fulfills a Cu sequestration role. Further studies will be required to reveal the physiological role of these peptides. For instance, it remains unclear whether the mature product of EGKCG pathways is a single modified polypeptide or smaller peptides generated by a host protease. There may also be additional factors involved in Cu-binding, such as DoxX proteins, which are not recapitulated by our heterologous expression or *in vitro* studies.

Altogether, our results show many common features for MNIO-PME1 BGCs. All characterized cases install 5TO moieties onto peptide substrates. Additionally, the products form during the microbial stress response and generally bind Cu. Given their mercaptan functional groups and Cu-binding functions, we propose the name captophorins (<u>C</u>u-<u>a</u>ssociated <u>p</u>eptidic thio<u>o</u>xazole-containing metallophores) for these compounds.

### *In vitro* reconstitution of FonBC

Having described captophorin biosynthesis, we next investigated the requirements for 5TO formation by reconstituting FonBC *in vitro*. We first generated a co-expression construct containing His_6_-FonB and FonC. After expression and purification, SDS-PAGE confirmed the presence of untagged FonC, suggesting a stable interaction with FonB (**Figures 5, S24**). Additionally, FonBC appeared brown in color, suggestive of Fe-binding. Native MS analysis identified that FonB and FonC formed a stable heterodimer and co-purified with 3 Fe atoms. MS analyses of purified FonA and NgoA revealed that both peptides co-purified with their respective BC heterodimers (**Figure S25**). Native MS analysis of the NgoA sample additionally revealed intact NgoBC and Ngo-ABC complexes, supporting a stable ternary interaction.

**Figure 5.**
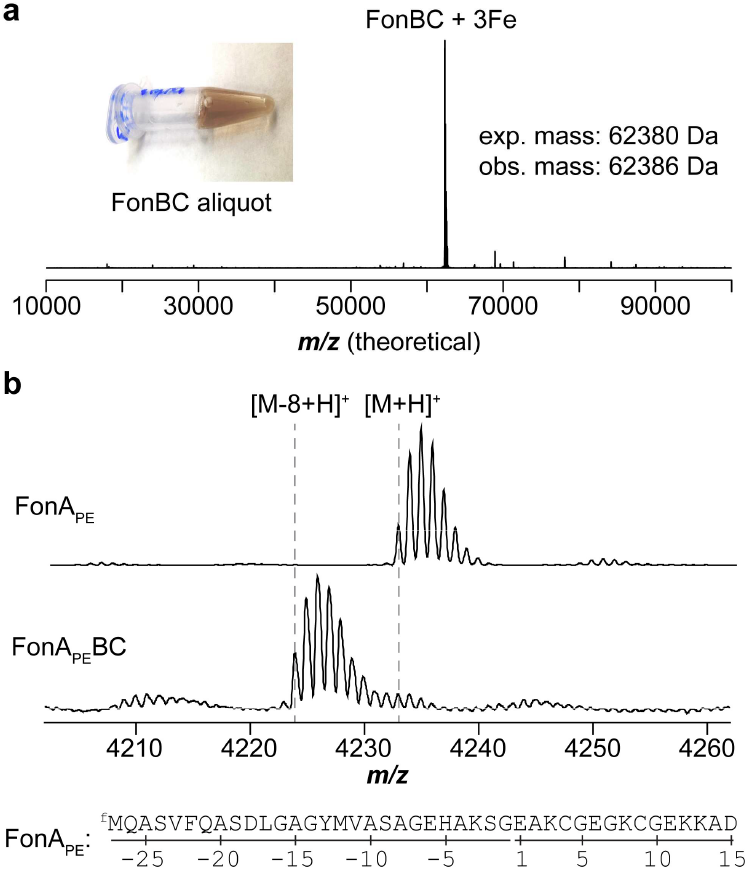
*In vitro* reconstitution of FonBC. **a**, Native MS data for FonBC and visual appearance of FonBC [∼1 mM]. **b**, Reconstitution of FonBC using *in vitro* translated FonA_PE_.

We next sought a minimal substrate for activity assays, as FonA contains seven sites of modification and is relatively large. Shortening the peptide length was expected to minimize aggregation and facilitate mass spectrometry analysis of enzymatic products. We generated DNA templates for *in vitro* transcription/translation of two shortened FonA peptides containing one or two EGKCG repeats, respectively (**Table S10, Figure S26**).^39^ After translation and aerobic reaction with purified FonBC, only the sequence with two repeats was modified, displaying a loss of 8 Da from the expected mass (**Figures 5, S26**). This double repeat sequence, hereafter FonA_PE_ (PE for PURExpress), was further characterized.

To investigate which residues of FonA_PE_ were critical for turnover, we performed a single-site Ala scan. DNA templates coding for these variants were synthesized by PCR and subjected to *in vitro* transcription/translation (**Table S10**). The resulting peptides were then reacted with FonBC and analyzed by MALDI-TOF-MS. These analyses revealed that FonBC was tolerant to most single-site Ala variants (**Figure S27**). All variants preceding the core region were tolerated, suggesting no individual position was strictly required for RRE engagement. Some core variants showed diminished turnover efficiency, such as FonA_PE_E6A. Both FonA_PE_E1A and FonA_PE_E11A could be modified twice, suggesting Glu was not strictly required preceding or following the modified Cys. Thus, the impaired turnover of FonA_PE_E6A suggests a more complex role for this site. These results could be explained by a model where Glu6 aids in active site repositioning, but this claim requires additional investigation.

Next, a panel of FonA_PE_ double variants, with equivalent amino acid substitutions to both repeats was examined (**Figure S28**). For example, variant FonA_PE_1-P places Pro at the first residue of each 5-mer repeat within FonA_PE_ to give <u>P</u>AKCG<u>P</u>GKCG in core residues 1-10, while FonA_PE_3-A yields EA<u>A</u>CGEG<u>A</u>CG (Peptide and DNA sequences, **Table S10**). Position 4 variants could not be processed, demonstrating the necessity of Cys modification. Across all sites, FonBC was intolerant to Pro, suggesting a requirement for substrate flexibility and/or backbone interactions. Among the non-Pro variants to positions 1 or 2, only FonA_PE_1-R (<u>R</u>AKCG<u>R</u>GKCG) could not be processed. Among position 3 variants, FonA_PE_3-A (EA<u>A</u>CGEG<u>A</u>CG), FonA_PE_3-R, and FonA_PE_3-T were tolerated, while FonA_PE_3-F and FonA_PE_3-E severely impacted processing. Position five processing was blocked by all variants except FonA_PE_5-A (EAKC<u>A</u>EGKC<u>A</u>), suggesting a strong preference for small residues. Altogether, FonBC retained activity against most substrate variants; however, variation of the residues flanking the critical Cys can result in diminished activity.

### Exploration of the captophorin biosynthetic landscape

To define the captophorins as a newly categorized RiPP class, we first profiled the putative precursor peptides. Our dataset of Cys-rich putative precursor peptides was analyzed for DUF2282 homology and Cys-containing motifs (**Dataset S2, Table S2**). All Cys-rich peptides were then used to generate an SSN, and nodes were annotated accordingly (**Figure S29**). This approach produced three large precursor peptide groups, specifically the DUF2282, Lys-Cys, and Cys-Lys groups. Among repeat-containing sequences, the most frequent motif was the EGKCG 5-mer found in fontiphorin, oxazolin, and gonophorin (**Figure 6, Dataset S2**).

**Figure 6.**
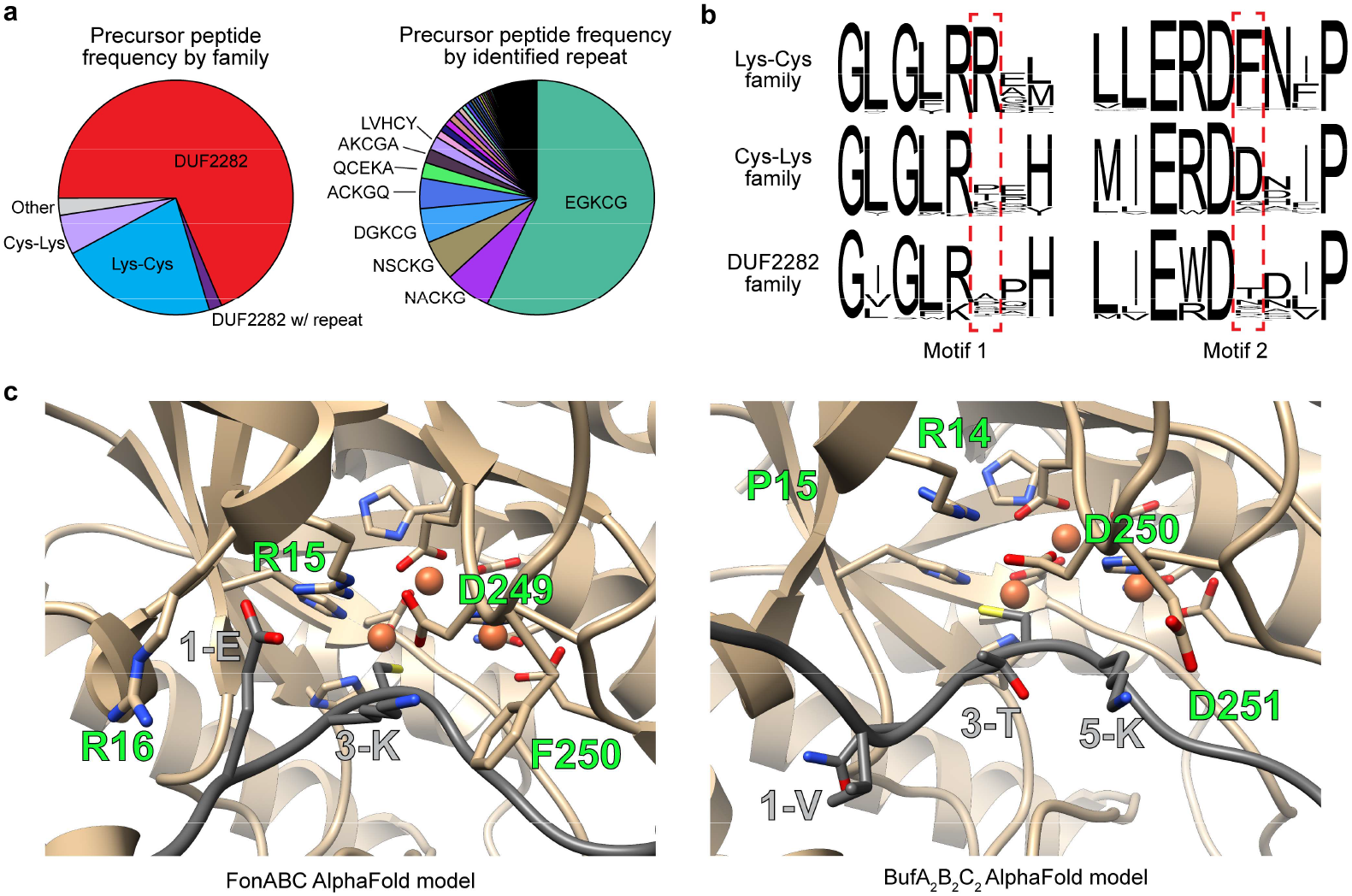
Factors affecting captophorin MNIO substrate specificity. **a**, Categorization of precursor peptides encoded in captophorin BGCs. Peptides were charted by general category (*n* = 21,424). Repeat containing precursor peptides were then charted by 5-mer repeat motif found within the sequence (*n* = 6,906). **b**, Sequence logos for captophorin MNIO enzymes (*n* = 100) focusing on putative determinants of substrate specificity. **c**, AlphaFold models for FonA_PE_BC (Lys-Cys) and BufA_2_B_2_C_2_ (Cys-Lys) active sites. Relevant residues involved in enzyme-substrate interaction or catalysis are shown as sticks.

While FonB (Lys-Cys group member) preferred a small residue in position five, we observed diverse residues surrounding Cys across other captophorin BGCs (**Figure S30**). To explore these cases, FonBC activity was screened against a set of commonly occurring 5-mer motifs with varying degrees of similarity to FonA (**Figure 6, Table S2**). For example, FonA_PE_AKCGA substitutes both repeats in FonA_PE_ to give a substrate with AKCGAAKGCGA in positions 1-10 (**Table S10**). Among the tested substrates, FonBC displayed varying degrees of turnover (**Figure S31**). The conservative FonA_PE_DGKCG variant could be fully modified, while FonA_PE_AKCGA, FonA_PE_GKCGT, and FonA_PE_NVCGG were modified once, consistent with our single- and double-site variant data above. In contrast to the previously observed preference for a Cys-Gly dyad, FonA_PE_LVHCY could be modified once, suggesting that the additional variation in the substrate allowed modification even with a Cys-Tyr sequence. Substrates containing a charged residue following Cys (FonA_PE_QCEKA, FonA_PE_HNDCK, and FonA_PE_NACKG) were not processed by FonBC, consistent with previously examined variants. Taken together, these data suggest family-specific enzyme-substrate interactions likely govern Cys selectivity as an independent feature from 5TO installation.

### Substrate selectivity among captophorin MNIOs

To investigate the determinants of specificity, we examined a FonA_PE_BC AlphaFold model (**Figures 6, S32**).^40^ This showed conservation of the MNIO metal-binding residues and proposed catalytic base (FonB-Asp249).^41,42^ The side chain of FonA_PE_Glu1 was placed directly in between FonB-Arg15 and FonB-Arg16, which likely explains the loss of modification in FonA_PE_1-R due to charge-charge repulsion. Furthermore, interactions were observed between FonA_PE_Lys3, FonA_PE_Gly5, and FonB-Phe250. We generated a sequence logo for 100 Lys-Cys MNIOs, which showed conservation of Arg and Phe at these positions (**Figures 6, S33**). We then generated sequence logos for Cys-Lys and DUF2282 MNIOs, respectively. Both logos showed a loss of conservation at the FonB-Arg16 site. At the FonB-Phe250 site, we observed a substitution of Phe for Asp in the Cys-Lys MNIOs. An AlphaFold model of the Cys-Lys BufA_2_B_2_C_2_ additionally suggested an interaction between the substrate Lys of BufA_2_ and BufB_2_-Asp251 (**Figure 6**). Within DUF2282 sequences, there is little conservation for the residues surrounding Cys, and there was a corresponding lack of conservation at this site of the MNIO sequence logo.

Using site-directed mutagenesis, vectors containing variants at these sites were generated. Co-expression of FonB-D249A with FonAC yielded unmodified peptide, consistent with its proposed role as a catalytic base (**Figure S34**).^41^ In contrast, Ala variants of FonB at Arg15, Arg16, and Phe250 yielded modified products containing a ∼305 nm absorbance maximum, consistent with 5TO installation. Similarly, the FonB-F250D variant also yielded 5TO-containing product. We subsequently purified the variant FonBC complexes and examined their activity. In most cases, these variants displayed similar activity to wild-type enzyme; however, we observed some changes in substrate tolerance. FonB-R15A could fully process FonA_PE_1-R (<u>R</u>AKCG<u>R</u>GKCG) unlike wildtype FonB (**Figure S35**). FonB-R16A, on the other hand, showed diminished turnover with either acidic or basic residues in position 1. Taken together, this suggests that Arg15 may play a gatekeeping role to prevent positively charged residues from binding with the wrong register, while Arg16 likely provides a productive electrostatic interaction with Glu. This could also explain why the second Arg residue is only conserved in Cys-Lys family MNIOs, which predominantly contain Glu at position 1 of their substrates. For the selectivity around Cys, FonB-F250D could fully process FonA_PE_3-F, while the wild-type enzyme could not (**Figure S36**). This suggests that the loss of Phe steric occupancy is compensated for by the variation to the substrate sequence. While additional factors are likely at play, our data show that these active site residues govern selectivity for the residues surrounding the modified Cys.

### Catalytic proposals for 5TO biosynthesis

Based on recent mechanistic models of MNIO reactions, we interpreted captophorin biosynthesis through a similar catalytic cycle as proposed for OxT formation in methanobactins (**Figure S37**).^9^ Briefly, OxT biosynthesis is proposed to start with a mixed-valent Fe(II)/Fe(III). The Fe(II) site then binds molecular oxygen to generate an Fe(III)– superoxo species capable of β-hydrogen abstraction on the coordinated cysteine residue of the substrate peptide. This is followed by oxidation to a thioaldehyde. In the proposed OxT pathway, the C-terminal N-H is deprotonated to give an amidate (**Figure S37**, step 1c), which attacks the Cys thioaldehyde and yields a 4-thioxo-2-azetidinone ring. Deprotonation of the N-terminal Cys N-H then provides an imidate nucleophile to attack C2, followed by tautomerization to give OxT. In doing so, this returns the active site Fe atoms to a mixed Fe(II)/Fe(III) state.

Given the above, two general mechanistic routes could be drawn for 5TO formation. In one, the mechanism could proceed to the azetidinone intermediate, followed by imidate nucleophile attack at C4 rather than C2 (step 2e). After tautomerization, this would then restore the amide bond and yield 5TO. Alternatively, the reaction could directly proceed from the thioaldehyde to a five-membered ring by imidate formation at the N-terminal peptide bond (step 3c). This imidate would then directly attack the thioaldehyde, bypassing the azetidinone intermediate and ultimately giving 5TO. As drawn, we raise the possibility that amide bond deprotonation could be controlled by Cys and O_2_ coordination to alternate Fe atoms. As with the OxT mechanism, both proposed 5TO mechanisms return the net Fe oxidation state to the beginning of the catalytic cycle.

Many questions remain regarding MNIO catalysis. Both OxT- and 5TO-forming MNIOs display increased turnover in the presence of ascorbate. It remains unclear if MNIOs require external reductants to perform multiple turnovers. Ascorbate may generate the initial Fe(II) to start the catalytic cycle but could also serve a “salvage” role by reducing labile or off-pathway Fe(III). Further, MNIOs have been reported with either two or three active site Fe atoms, and the role of the third Fe atom is also unclear. Beyond the metal center, little investigation of substrate scope has been shown for MNIOs. Evaluation of FonBC suggests enzymatic preference for a small residue after Cys. Accordingly, we speculate that mechanism two, which directly involves this site in 4-thioxo-2-azetidinone formation, could explain this selectivity. Of course, detailed mechanistic investigation will be required to confirm or refute these proposals.

## Conclusions

In this work, we identified and characterized a widespread family of MNIOs responsible for captophorin biosynthesis. Using various spectroscopic and spectrometric data, we show that the MNIO enzyme, FonB, installs seven 5TO moieties onto its peptide substrate to produce fontiphorin. We further characterized oxazolin and gonophorin as additional examples and demonstrate that these contain 5TO moieties rather than oxazolin’s previously reported OxT.^17^ Together with Leprevost et al.,^16^ this suggests that PME1-dependent MNIOs are broadly involved in 5TO formation. The shared 5TO moiety suggests that bufferins, oxazolin, fontiphorin, bulbicupramide, and gonophorin all belong to a single RiPP class, which we propose naming captophorins (Cu-associated peptidic thiooxazole metallophores). While differentiation of proton-deficient heterocycles, such as 5TO, represents a common challenge in natural product chemistry,^43,44^ we describe a replicable approach for differentiation of 5TO and OxT using UV absorbance, chemical reactivity, and NMR spectroscopy. With clear distinctions now drawn between methanobactins and captophorins, further efforts can be directed toward enzyme mechanisms and biological functions.

Despite sharing common chemistry, captophorin BGCs encode diverse substrate sequences and architectures. Thus, we investigated their substrate selectivity using cell-free enzyme assays. In this case, our substrate scope data for FonB revealed that residues within the repeat motif dictate whether catalysis can proceed. Based on these data, we bioinformatically investigated putative active site residues and hypothesized sites of enzyme-substrate interaction. Using additional cell-free enzyme assays, we investigated these proposed sites within 5TO-forming MNIOs and rationalized sequence features across different MNIO groups. As further hypotheses are generated, cell-free assays will enable rapid generation of additional data to accelerate our understanding of enzyme function.

Through our efforts to connect 5TO-forming MNIOs, captophorins are now among the most populous RiPP classes encoded in public genomic data, with more than 20,000 identified examples in NCBI RefSeq data alone. The abundance of captophorin BGCs suggests they provide a strong selective advantage, most likely in heavy metal and oxidative stress response. Further biological studies will be required to link peptidic 5TO-containing metallophores to a specific molecular function. We note that captophorins are depleted from certain taxa, suggesting 5TO biosynthesis does not provide a universal advantage for these organisms. While this could amount to infrequent exposure to oxidative stress or heavy metals, there are likely additional factors to consider. Investigation of captophorins may reveal hidden relationships between microbial physiology, oxidative stress response, and lifestyles of diverse Pseudomonadota. Broadly, genomics-guided exploration of biosynthesis will continue to offer an exciting entryway to explore physiology across the tree of life.

## ASSOCIATED CONTENT

### Supporting Information

Materials, Methods, Supplementary Note, Tables S1-S10, and Figures S1-S37 showing additional bioinformatic, spectral, and experimental data.

Dataset S1. Captophorin BGCs identified in NCBI RefSeq data.

Dataset S2. Captophorin precursor peptides.

## ACKNOWLEDGMENT

The authors thank the Mass Spectrometry and NMR Core Facilities at University of Illinois Urbana-Champaign and Vanderbilt University for assistance with data acquisition. The authors further thank the Vanderbilt Center for Structural Biology for instrumentation access and Grace Kenney (Univ. of Pittsburgh) for helpful early discussions.

## ABBREVIATIONS

MNIO: multinuclear nonheme iron-dependent oxidative enzyme
5TO: 5-thiooxazole
RiPP: ribosomally synthesized and post-translationally modified peptide
PTM: post-translational modification
BGC: biosynthetic gene cluster
RRE: RiPP recognition element
PME1: Type 1 partner protein of MNIO
SSN: sequence similarity network
WHO: World Health Organization
IMAC: immobilized metal affinity chromatography
MS: mass spectrometry
LCMS: liquid chromatography-mass spectrometry
HRMS: high resolution mass spectrometry
MALDI-TOF: matrix assisted laser desorption/ionization-time of flight
IAA: iodoacetamide
HRMS/MS: high resolution mass spectrometry/mass spectrometry
OxT: oxazolone/thioamide
MBP: maltose binding protein
HPLC: high performance liquid chromatrography

## For Table of Contents graphic

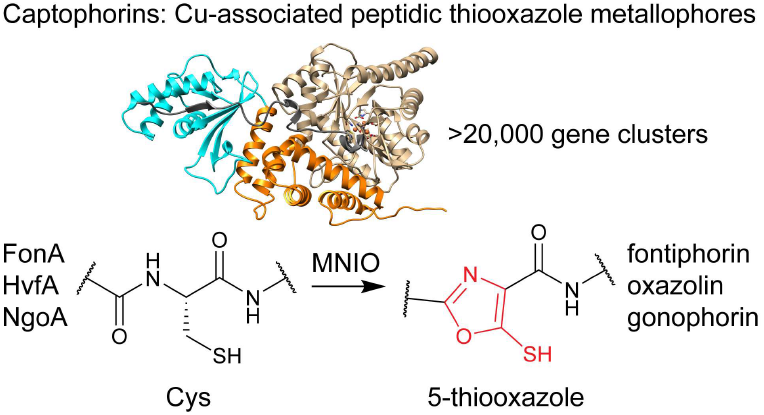

