## Supporting Information for "Biosynthesis of peptidic thiooxazole metallophores installed by multinuclear nonheme iron enzymes"

### Table of Contents

Materials and Methods .......... S4
Note S1: cluster_to_logo.py. .......... S8
Table S1. Protein co-occurrence in MNIO-PME1 BGCs. .......... S9
Table S2. Precursor peptide classification across MNIO-PME1 BGCs. .......... S10
Table S3. Taxonomic distribution of MNIO-PME1 BGCs with identified precursor peptides. .......... S11
Table S4. Proteins encoded in the *fon*, *hvf*, and *ngo* BGCs. .......... S13
Table S5. Key gene sequences from the *fon*, *hvf*, and *ngo* BGCs. .......... S14
Table S6. Key protein sequences from the *fon*, *hvf*, and *ngo* BGCs. .......... S16
Table S7. Vectors used in this study. .......... S17
Table S8. Table of daughter ion assignments for HR-MS/MS analysis of alkylated $FonA_{67-80}$. .......... S18
Table S9. NMR assignments for alkylated $FonA_{67-80}$. .......... S19
Table S10. Constructs generated for *in vitro* transcription/translation. .......... S20
Figure S1. Sequence logos for putative MNIO-PME1 families identified from RRE-Finder data. .......... S23
Figure S2. Sequence similarity network of MNIO family. .......... S26
Figure S3. Sequence similarity network for PME1 family. .......... S27
Figure S4. MNIO BGCs from *H. somni* and *N. gonorrhoeae*. .......... S28
Figure S5. UV-Vis spectral comparison of unmodified FonA to fontiphorin. .......... S29
Figure S6. Deconvoluted mass spectrum of full-length fontiphorin. .......... S30
Figure S7. MALDI-LIFT-MS for $FonA_{67-80}$. .......... S31
Figure S8. HRMS/MS analysis of alkylated $FonA_{67-80}$. .......... S32
Figure S9. NMR spectra for $FonA_{67-80}$. .......... S33
Figure S10. Spin system assignments for unmodified residues in $FonA_{67-80}$. .......... S37
Figure S11. Zoom-in showing anomalous $FonA_{67-80}$ $^{1}H$ and $^{13}C$ signals. .......... S38
Figure S12. Assignment/differentiation of $FonA_{67-80}$ Lys residues. .......... S39
Figure S13. $FonA_{67-80}$ structure. .......... S40
Figure S14. Comparison of NMR and UV absorbance data for MNIO products. .......... S41
Figure S15. Comparison of MNIO and PME1 proteins. .......... S42
Figure S16. Purification and alkylation of oxazolin peptides. .......... S43
Figure S17. NMR spectra of the proteolytically digested oxazolin sample. .......... S44
Figure S18. Gonophorin UV and MS data. .......... S46
Figure S19. Gonophorin NMR data. .......... S47
Figure S20. Native MS of MBP-FonABC. .......... S49
Figure S21. LC retention time shift during fontiphorin Cu-binding assay. .......... S50
Figure S22. Dynamic light scattering for fontiphorin. .......... S51
Figure S23. LC-MS spectra of gonophorin with Cu loaded *in vivo* .......... S52
Figure S24. Purification of FonBC dimer. .......... S53
Figure S25. FonA and NgoA co-purify with their respective MNIO and PME1 .......... S54
Figure S26. Initial reconstitution of FonBC. .......... S55
Figure S27. FonB substrate tolerance for single-site variants. .......... S56
Figure S28. FonB substrate tolerance for double-site variants. .......... S57
Figure S29. Sequence similarity network for putative precursor peptides. .......... S59
Figure S30. Biosynthetic protein sequence similarity networks colored by precursor peptide family. .......... S60
Figure S31. FonB substrate tolerance for naturally occurring 5-mer repeats. .......... S62
Figure S32. AlphaFold model for $FonA_{PE}BC$. .......... S63

Figure S33. MNIO sequence logos by substrate family. .......... S64
Figure S34. FonB variant heterologous co-expressions. .......... S67
Figure S35. *In vitro* activity of FonB Ala variants. .......... S68
Figure S36. FonB-F250D substrate tolerance. .......... S69
Figure S37. Mechanistic proposals for 5-thiooxazole installation. .......... S70
Supplementary References. .......... S72

### Materials and Methods

**Materials.** All chemicals were used without further purification unless stated otherwise. Antibiotics were purchased from Goldbio. Enzymes were purchased from New England Biolabs (NEB). DNA oligonucleotides were purchased from Integrated DNA Technologies, Inc. or Twist Biosciences. Sources for experiment-specific materials and reagents are provided in the relevant sections below.

**General molecular biology methods.** General PCR was performed using Q5 Polymerase (NEB) following manufacturer-recommended protocols. New plasmids were assembled using 2x HiFi DNA Assembly Master Mix (NEB), and site-directed mutagenesis was performed using the Quikchange (Agilent) protocol. Where appropriate, PCR reactions were treated with DpnI (NEB) to digest template DNA. Gibson assembly and site-directed mutagenesis reactions were transformed into chemically competent *E. coli* DH5α, and successful transformants were selected using solid LB-Miller medium (10 g/L tryptone, 5 g/L yeast extract, 10 g/L NaCl, 20 g/L agar) supplemented with 50 µg/mL kanamycin. Colonies were then individually grown in 5 mL liquid LB-Miller. Plasmids were then purified from these cultures by alkaline lysis, and plasmid sequences were confirmed by Oxford nanopore sequencing (Plasmidsaurus).

**Bioinformatics.** RRE-containing BGC families identified in Ren et al.[1] were gathered and hypothetical precursor peptides were converted to FASTA format. We adapted a script developed by He et al.[2,3] to generate a sequence logo for each set of putative precursor peptides. FASTA files of precursor peptides were aligned using command line mafft and alignments were converted into sequence logos with low-occupancy columns removed for clarity (**Supplementary Note**). Using matplotlib, sequence logos were output directly for user analysis. Using RODEO,[4] general co-occurrence features were examined for these RRE-containing BGCs. Comparison of gene co-occurrence and precursor sequence logos allowed prioritization of MNIO-containing BGCs for further analysis.

Sequence similarity networks (SSNs) were generated using the EFI-EST[5] webpage (https://efi.igb.illinois.edu/efi-est/) and visualized using Cytoscape. The MNIO SSN (**Figure S2**) was generated using the family query option for Interprofam IPR007801.[6] The PME1 SSN (**Figure S3**) was generated using the family query option for PfamID PF09836. Annotations of these SSNs were calculated using the EFI-GNT tool, after which local protein family matches were downloaded for each accession, and nodes were colored in Cytoscape, accordingly.

Using NCBI protein family models, all RefSeq accession IDs retrieved by NCBIFam NF021364 were downloaded. These accessions were used as input for RODEO and local genomic neighborhoods were analyzed to identify local pairs of PME1 and MNIO proteins. For loci containing matches to both PF09836 and PF05114, co-occurrence of local proteins was calculated (**Table S1**). Following RODEO analysis, the complete set of open reading frames (ORFs) encoded in MNIO-PME1 BGCs were analyzed for Cys content, and ORFs containing <2.5% Cys content or fewer than 2 Cys residues were removed. The complete set of ORFs was also examined for retrieval by the DUF2282 HMM (PfamID: PF09836). Using these criteria, we identified 24,105 Cys-rich ORFs. Using EFI-EST, we generated a precursor peptide SSN with individual sequences conflated above 70% identity. Each ORF was analyzed for 5-mer repeat motifs containing a Cys. The most abundant 5-mer motif within the sequence was listed. Nodes within the SSN were then analyzed by the presence and content of the 5-mer motif identified for the representative ORF. Subsequently, SSNs were generated for MNIOs and PME1s found in Refseq data. These were additionally annotated by the identified family of precursor peptide encoded within the local genomic region.

**Taxonomic analysis of MNIO-PME1 BGCs.** Retrieved NCBI RefSeq accession IDs for MNIOs were mapped to organism lineage using a local install of the NCBI Entrez utilities. In summary, files containing lists of accession IDs were posted to NCBI, and an XML record was retrieved and parsed using the command below. The resulting parsed XML data was then used to match individual accession IDs to their taxonomy.

Entrez command: epost -db protein -input mnio_ids.txt | efetch -db protein -format xml | xtract -pattern Seq-entry -element Textseq-id_accession,OrgName_lineage > mnio_xml.txt

To analyze abundance in reference genomes, all NCBI Pseudomonadota reference genomes ($n$ = 1,919) were downloaded. Genome records were then analyzed for encoded PME1 proteins. Corresponding accession IDs were used as input for RODEO analysis, and local MNIO proteins were identified. The NCBI taxIDs for the reference genomes were used to generate a phylogenetic tree using phyloT. The tree was then visualized with iTOL and annotated by the presence or absence of an MNIO-PME1 local pair within the reference genome.

**Expression and purification for MNIO-PME1 BGCs.** Vectors encoding the protein(s) of interest were transformed into chemically competent *E. coli* BL21(DE3) cells. Cells were grown on solid LB-Miller media containing 50 μg/mL kanamycin overnight. Single colonies were used to inoculate 10 mL of LB medium containing 50 μg/mL kanamycin grown at 37 °C with shaking at 250 rpm. After 18 h, 1 L of LB was inoculated with the starter culture after addition of 50 μg/mL kanamycin. Cells were grown to an optical density ($OD_{600}$) of 0.6-0.8 then placed on ice for 30 min. Isopropyl β-D-1-thiogalactopyranoside (IPTG) and ferrous ammonium sulfate were added to final concentrations of 0.5 mM and 200 μM, respectively. The cultures were grown for an additional 18 h at 18 °C with shaking at 250 rpm. Cells were harvested by centrifugation at 4,500 × g for 15 min, washed once with phosphate-buffered saline and re-centrifuged at 4,500 × g for 15 min. Cell pellets were flash-frozen in liquid nitrogen and stored at -80 °C.

For purification, cell pellets were thawed on ice and resuspended in 30 mL of lysis buffer [50 mM HEPES, 500 mM NaCl, 10 mM imidazole, 5 mM β-mercaptoethanol, 2.5% glycerol (*v/v*), and 0.1% triton X-100 (*v/v*), pH 7.5] with additional lysozyme at 4 mg/mL. The mixture was then subjected to sonication for 45 s followed by gentle rocking for 10 minutes while being kept at 4 °C. This process was repeated a total of three times. Cellular debris was removed by centrifugation (20,000 × g, 90 min) and the supernatant was applied to Ni-NTA resin pre-equilibrated with lysis buffer. The column was washed with 30 column volumes (CV) of wash buffer [50 mM HEPES, 500 mM NaCl, 20 mM imidazole, 5 mM β-mercaptoethanol, 2.5% glycerol (*v/v*), pH 7.5]. $His_6$-tagged peptides/proteins were eluted using 5 CV of elution buffer [50 mM HEPES, 500 mM NaCl, 250 mM imidazole, 5 mM β-mercaptoethanol, 2.5% glycerol (*v/v*), pH 7.5]. Eluent was concentrated using an Amicon ultracentrifugal filter (EMD Millipore) with an appropriate molecular weight cutoff. Proteins were buffer exchanged with 10× volume of protein storage buffer [50 mM HEPES, 500 mM NaCl, 2.5% glycerol (*v/v*), and 0.5 mM TCEP, pH 7.5] before being flash frozen in liquid nitrogen and stored at -80 °C.

**Matrix-assisted laser desorption/ionization time-of-flight mass spectrometry (MALDI-TOF-MS).** Matrix-assisted laser desorption/ionization time-of-flight mass spectrometry (MALDI-TOF-MS) analysis and MALDI-LIFT-TOF/TOF-MS was performed using a Bruker UltrafleXtreme MALDI TOF-TOF mass spectrometer (Bruker Daltonics) in reflector positive mode at the University of Illinois School of Chemical Sciences Mass Spectrometry Laboratory and in the Mass Spectrometry Research Core at Vanderbilt University. All samples were analyzed in reflector positive mode. After collection of MS/MS data, fragment ions were annotated using the Interactive Peptide Spectral Annotator webtool.

**Liquid chromatography-mass spectrometry.** Samples were analyzed by a Shimadzu LC-MS-2020 single quadrupole system under positive mode electrospray ionization. For the liquid chromatography component, a ThermoFisher Scientific Hypersil GOLD column (150 × 2.1 mm) was employed. Solvents A ($H_2O$) and B (acetonitrile) both contained 0.1% formic acid. The LC method began with equilibration at 10% B for 5 min following injection, followed by a gradient from 5-70% B over 20 min, held at 90% B for 5 min, ramped down to 5% B over 1 min, and finally held at 5% B for 10 min to re-equilibrate the column.

**High resolution and tandem mass spectrometry.** An ESI mix composed of 80% acetonitrile, 19% water, and 1% acetic acid was used to resuspend purified peptides. Samples were analyzed via a ThermoFisher Scientific Orbitrap Fusion ESI-MS using an Advion TriVersa Nanomate 100. MS calibration was performed with Pierce LTQ Velos ESI Positive Ion Calibration Solution (ThermoFisher). The spectrometer was operated using the following parameters: 100,000 resolution, 1 m/z isolation width (MS/MS), 35 normalized collision energy (MS/MS), 0.4 activation q value (MS/MS), and 30 ms activation time (MS/MS). Fragmentation was performed using collision-induced dissociation (CID) at 30% and 70%. Data analysis was conducted using the Qualbrowser application of Xcalibur software (ThermoFisher Scientific). The Interactive Peptide Spectral Annotator (webtool developed by Coon Laboratory, University of Wisconsin) was used to annotate MS/MS spectra.

**Mass spectral deconvolution.** Low-resolution mass spectrometry data were deconvoluted using the UniDec software package (version 8.0.3; downloaded from https://github.com/michaelmarty/UniDec)[7]. Raw $MS^1$ spectra were exported from the instrument as centroided mass-to-charge (m/z) versus intensity text files and imported into UniDec without additional preprocessing. Spectra were subjected to background subtraction and mild smoothing using the default UniDec settings, and intensities were normalized to the most intense peak in each spectrum. Deconvolution was carried out over a mass range chosen to encompass the expected analyte mass ± ~50% and a charge-state range broad enough to include all observed charge states (typically z = 1–30 for intact proteins/peptides), using a Gaussian peak shape appropriate for quadrupole analyzers. The "Sample Mass Every" (bin size) parameter was set to 1 Da. Regularization parameters (charge smooth width, mass smooth width, and point smooth width) were kept at their default values. Deconvoluted zero-charge mass spectra were exported as comma-separated values and used for figure generation.

**General HPLC purification of modified compounds.** Upon isolation and concentration of modified peptides by Ni-NTA purification, formic acid and acetonitrile were added to generate final compositions of 0.1% and 20% (*v/v*), respectively. This mixture was subjected to centrifugation at 17,000 × g for 10 min. The clarified supernatant was then subjected to HPLC purification on a ThermoFisher Scientific Vanquish instrument equipped with an Hypersil GOLD $C_{18}$ column (150 × 4.6 mm). Solvents A ($H_2O$) and B (acetonitrile) both contained 0.1% formic acid. The following general method was used: equilibration for 5 min at 10% B, gradient of 10-50% B for 15 min, ramp up to 90% B over 1 min, held at 90% B for 5 min, ramp down to 10% B over 1 min, and then held at 10% B for 10 min. Fractions were collected based on absorbance at 305 nm.

**Iodoacetamide alkylation.** To a solution of purified modified peptide in alkylation buffer [25 mM HEPES, 50 mM NaCl, 2 mM TCEP, pH 8], a 10x molar excess of iodoacetamide was added. The reaction was allowed to proceed for 1 h at RT while being protected from light.

**UV-Vis spectroscopy.** UV-Vis absorption features were monitored by HPLC using a ThermoFisher Vanquish instrument equipped with a Photodiode Array Detector (see above). For compounds not undergoing HPLC purification, UV-Vis absorption spectra were collected using a NanoDrop 2000 spectrophotometer with no baseline correction and following a buffer-only blank sample.

**NMR spectroscopy.** Dried, purified peptide sample was resuspended in an appropriate volume of NMR solvent mixture (9:1 $H_2O$:$D_2O$ + 0.1% formic acid-$d_2$) prior to analysis. All one and two-dimensional NMR data were collected on a Bruker Avance NEO 600 MHz spectrometer with a 5-mm prodigy BBO probe using Topspin 4.1.4 software. Typical acquisition parameters included: 1.5 s relaxation delay, 32–64 scans, 16 dummy scans, 2048 × 256–400 data points, 11 ppm 1H spectral width and 220 ppm spectral width in $^{13}C$. A mixing time of 80 ms was employed for TOCSY. The dpfgse (double pulsed field gradient spin echo) pulse sequence was used for water suppression (if required). All spectra were processed in MestReNova 14.3.0.

**High resolution native mass spectrometry.** Native mass spectrometry experiments were performed on a Thermo Scientific Q Exactive Orbitrap mass spectrometer operated in positive-ion nanoelectrospray ionization (nanoESI) mode and configured for high–mass range analysis. Protein samples were buffer-exchanged into 150–200 mM ammonium acetate (pH 6.8–7.5) using Amicon molecular weight cutoff centrifugal filters immediately prior to analysis. Typically, 2–3 µL of sample were loaded into gold-coated borosilicate capillaries (1.0–1.2 mm outer diameter) and introduced by static nanoESI. Source conditions were tuned to maximize desolvation while preserving native-like charge state distributions, with a spray voltage of 1.0–1.6 kV, capillary temperature of 150–200 °C. Spectra were acquired in full-scan mode over an m/z range of 1,000–8,000 (up to 25,000 m/z for higher-mass complexes), with Orbitrap resolving power typically set between 15,000 and 60,000 at m/z 200, a trapping gas pressure setting of 3–7, an automatic gain control (AGC) target of ~$1 \times 10^6$, a maximum injection time of 100–200 ms, and 5–10 microscans per transient. Instrument calibration was performed using native protein standards according to the manufacturer's recommendations. Raw data were inspected in Xcalibur (Thermo Scientific), exported as centroided spectra, and deconvoluted to zero-charge mass distributions using UniDec with parameters optimized for native protein spectra as described above.

**Copper-binding assays.** To purified peptides/proteins in protein storage buffer, a solution of Cu-glycinate and sodium ascorbate was added to final concentrations of 10 mM and 20 mM, respectively. The mixture was placed on ice for 30 min, followed by mass spectral analysis.

*In vivo* copper-loading experiments were conducted as above. Following induction of control and test cultures with IPTG, $CuSO_4$ was added to the test culture to a final concentration of 50 μM. After overnight expression, purification proceeded immediately without freezing of cell pellets. Eluents following Ni-NTA purification were buffer exchanged 500-fold into protein storage buffer using a 3 kDa molecular weight cutoff centrifugal filter. Subsequently, samples were analyzed by LCMS.

**Dynamic light scattering.** Dynamic light scattering (DLS) data was gathered using a Wyatt DynaPro NanoStar instrument. Measurements were taken using NanoStar disposable microcuvettes after equilibration to 25 °C for 3 min. Measurements were obtained following laser auto-attenuation to avoid detector saturation. Ten measurements were taken for each sample and averaged, excluding any samples with poor signal-to-noise ratio. This was repeated three times, and data represents the average of the three individual replicates. Lyophilized samples of fontiphorin were dissolved in 25 mM PIPES buffer (pH 6.8) containing 5 mM dithiothreitol (DTT) to a starting concentration of 250 μM. Fontiphorin samples were then diluted 5-fold to obtain good signal-to-noise ratios. Cu solutions were generated by preparing a 50 mM $CuSO_4$ stock solution in 25 mM PIPES buffer without DTT. A 250 mM sodium ascorbate stock solution was also prepared and added in a 5:1 ratio to give each $CuSO_4$ solution. Resulting Cu(I) solutions were then diluted with PIPES buffer and an equal volume was added to the fontiphorin sample for measurement. Each combination was prepared immediately before mixing with fontiphorin. Control Cu sample was prepared by adding DTT-containing PIPES buffer without fontiphorin.

**Production of DNA templates for in vitro transcription/translation.** Linear double-stranded DNA templates for cell-free reactions were generated by overlap-extension PCR of smaller single-stranded primer sequences. DNA sequences of all templates used for in vitro transcription/translation reactions are provided in Table S10.

**Enzyme assays using in vitro transcribed/translated substrate peptides.** In vitro transcription/translation reactions were performed using the commercially available PURExpress kit (NEB). Typical transcription/translation reactions were set up at a 5 μL scale by mixing 2 μL Solution A, 1.5 μL Solution B, and 1.5 μL of a DNA template dissolved in water at concentrations of > 200 ng/μL. Transcription/translation reactions were allowed to proceed for 3 h at 37 °C. Next, 20 μL of a FonBC reaction mixture was added to the transcription/translation reaction. This FonBC reaction mixture consisted of 25 mM HEPES, 50 mM NaCl, 1 mM Fe(II) ammonium sulfate, 10 mM sodium ascorbate, 5 mM DTT, and 100 μM FonBC. Following this, the tube containing the mixture was shaken in a thermomixer at 400 rpm and 30 °C for 1 h. Afterwards, 3 μL of a 1% formic acid solution in water was added to the mixture to induce precipitation of proteins (final concentration: ~0.1% formic acid). Following centrifugation, the clarified supernatant was transferred to a fresh tube, desalted by a $C_{18}$ ZipTip following standard operating protocols, and analyzed by MALDI-TOF-MS using SDHB as the matrix.

**Supplementary Note: cluster_to_logo.py.** Script for logo generation from FASTA files. Adapted from He et al.[2]

```
import numpy as np
import pandas as pd
import matplotlib.pyplot as plt
import logomaker as lm
import os, time, argparse, concurrent.futures
from Bio.Align.Applications import MafftCommandline
from Bio import AlignIO

try:
    import StringIO
except ImportError:
    from io import StringIO
start = time.perf_counter()

def makelogo(fasta_file,out_dir):
    mafft_cline = MafftCommandline(input=fasta_file)
    mafft_cline.op = 3.0
    mafft_cline.ep = 3.0
    stdout, stderr = mafft_cline()
    prefix = os.path.basename(fasta_file)
    aligned_path = out_dir + f'/{prefix}.aln'
    with open(aligned_path, "w") as handle:
        handle.write(stdout)
    align = AlignIO.read(aligned_path, "fasta")
    seqs = []
    for seq_record in align:
        seqs.append(str(seq_record.seq))

    pre_aln_df = lm.alignment_to_matrix(sequences=seqs, to_type='counts', characters_to_ignore='.X')
    pre_aln_df = lm.transform_matrix(pre_aln_df, from_type='counts', to_type='information')
    temp_aln_df = pre_aln_df.drop('-', axis=1)
    scalar_df = pre_aln_df.sum(axis=1)/temp_aln_df.sum(axis=1)
    scalar_df = scalar_df.replace([np.inf, -np.inf], 1.0)
    scalar_df = scalar_df.mask(scalar_df < 1.0, 1.0)
    new_aln_df = pre_aln_df
    new_aln_df = new_aln_df.apply(lambda row: row.where(row >= row.mean(), 0), axis=1)
    new_aln_df = new_aln_df.drop('-', axis=1)
    new_aln_df = new_aln_df.loc[(new_aln_df!=0).any(axis=1)]
    new_aln_df.reset_index(inplace=True)
    new_aln_df = new_aln_df.drop('pos', axis=1)
    print(len(new_aln_df))
    pre_aln_logo = lm.Logo(new_aln_df, font_name='sans',
                           figsize=(18,1.5),  # <= 50AA (15, 2)  <=100 (25,2)
                           stack_order='big_on_top', vpad=0.02,
                           color_scheme='weblogo_protein',
fade_probabilities=False,baseline_width=1,show_spines=True)
    plt.savefig(out_dir+ f'/{prefix}.png', bbox_inches='tight', dpi=600)
    plt.close()

if __name__ == '__main__':
    parser = argparse.ArgumentParser(description='Perform sequence alignment and then generate logo')
    parser.add_argument('-i', '--input_fasta', type=str, metavar='', required=False,
                        help='Folder contains to-be-processed protein or nucleotide sequence files')
    parser.add_argument('-o', '--outdir', type=str, metavar='', required=True,
                        help='A  path to save all files, e.g., data/output')
    parser.add_argument('-n', '--num_threads', type=int, metavar='', required=False,default=1,
                        help='<-n 4> Number of threads,(default n=1)')
    args = parser.parse_args()

    if not os.path.exists(args.outdir):
        os.makedirs(args.outdir)

    with concurrent.futures.ProcessPoolExecutor(args.num_threads) as executor:  # 24 threads
        for fasta_file in os.listdir(args.input_fasta):
            print(f'Processing {fasta_file}')
            makelogo(args.input_fasta+fasta_file, args.outdir)
            print(f'Finished {fasta_file}')

    finish = time.perf_counter()
    print(f'finished in {round(finish - start, 3)} seconds')
```

**Table S1. Protein co-occurrence in MNIO-PME1 BGCs.**

| PfamID | Pfam Name | Pfam Description | Frequency |
|---|---|---|---|
| PF05114 | DUF692 | Multinuclear non-heme iron dependent oxidative enzyme (MNIO) | 99.8% |
| PF09836 | DUF2063 | MNIO-binding domain of partner protein | 99.5% |
| PF07681 | DoxX | DoxX | 67.8% |
| PF10048 | DUF2282 | Non-repeat precursor peptide | 55.1% |
| PF03466 | LysR_substrate | LysR substrate binding domain | 22.9% |
| PF06532 | NrsF | Negative regulator of sigma F | 14.1% |
| TIGR02937 | TIGR02937 | RNA polymerase sigma-70 factor | 9.3% |
| TIGR02154 | TIGR02154 | PhoB transcriptional regulatory protein | 8.6% |
| TIGR02943 | TIGR02943 | RNA polymerase sigma-70 factor | 8.6% |
| TIGR01386 | TIGR01386 | CopS-like heavy metal sensor kinase | 8.4% |

**Table S2. Precursor peptide classification across MNIO-PME1 BGCs.**

| Precursor Family | # BGCs | Repeat Family | # BGCs |
|---|---|---|---|
| DUF2282 | 15,077 | None | 14,518 |
| | | Cys-Lys | 358 |
| | | Lys-Cys | 21 |
| | | Other repeat | 180 |
| Repeat-containing ORFs | 6,347 | Lys-Cys | 4,667 |
| | | Cys-Lys | 1,172 |
| | | Other repeat | 508 |
| Other Cys-rich ORFs | 2,681 | N/A | |

**Table S3. Taxonomic distribution of MNIO-PME1 BGCs with identified precursor peptides.**

| Phylum | # BGCs | Class | # BGCs | Order | # BGCs |
|---|---|---|---|---|---|
| Pseudo-monadota | 20,131 | Gamma-proteobacteria | 12,235 | Pseudomonadales | 3,760 |
| | | | | Vibrionales | 2,117 |
| | | | | Alteromonadales | 1,473 |
| | | | | Lysobacterales | 1,351 |
| | | | | Aeromonadales | 1,013 |
| | | | | Moraxellales | 525 |
| | | | | Oceanospirillales | 497 |
| | | | | Cellvibrionales | 257 |
| | | | | Pasteurellales | 231 |
| | | | | Legionellales | 220 |
| | | | | Enterobacterales | 195 |
| | | | | Chromatiales | 168 |
| | | | | Thiotrichales | 149 |
| | | | | Methylococcales | 144 |
| | | | | Nevskiales | 31 |
| | | | | Gammaproteobacteria incertae sedis | 19 |
| | | | | Steroidobacterales | 15 |
| | | | | Kangiellales | 13 |
| | | | | Acidiferrobacterales | 8 |
| | | | | Immundisolibacterales | 6 |
| | | | | Thiohalomonadales | 5 |
| | | | | Thiohalobacterales | 5 |
| | | | | Candidatus Berkiellales | 4 |
| | | | | Arenicellales | 4 |
| | | | | Cardiobacteriales | 4 |
| | | | | Thiohalorhabdales | 3 |
| | | | | Salinisphaerales | 2 |
| | | | | Xanthomonadales | 2 |
| | | | | Methylohalomonadales | 1 |
| | | | | Thiohalospirales | 1 |
| | | Alpha-proteobacteria | 4,549 | Hyphomicrobiales | 2,113 |
| | | | | Rhodobacterales | 1,271 |
| | | | | Sphingomonadales | 615 |
| | | | | Rhodospirillales | 229 |
| | | | | Caulobacterales | 196 |
| | | | | Maricaulales | 25 |
| | | | | Kordiimonadales | 19 |
| | | | | Geminicoccales | 15 |
| | | | | Acetobacterales | 14 |
| | | | | Sneathiellales | 12 |
| | | | | Emcibacterales | 10 |
| | | | | Iodidimonadales | 6 |
| | | | | Rickettsiales | 5 |
| | | | | Hyphomonadales | 5 |
| | | | | Minwuiales | 2 |
| | | | | Holosporales | 1 |
| | | | | Parvularculales | 1 |
| | | Beta-proteobacteria | 3,279 | Burkholderiales | 2,400 |
| | | | | Neisseriales | 523 |
| | | | | Nitrosomonadales | 198 |
| | | | | Rhodocyclales | 102 |
| | | | | Betaproteobacteria incertae sedis | 48 |
| | | | | Ferrovales | 4 |

**Table S3 (cont.). Taxonomic distribution of MNIO-PME1 BGCs with identified precursor peptides.**

| Phylum | # BGCs | Class | # BGCs | Order | # BGCs |
|---|---|---|---|---|---|
| Pseudo-monadota (cont.) | 20,131 | Acidithiobacillia | 52 | Acidithiobacillales | 52 |
| | | Candidatius Mariprofundia | 3 | Mariprofundales | 3 |
| | | Hydrogenophilia | 2 | Hydrogenophilales | 1 |
| | | | | Hydrogenophilia incertae sedis | 1 |
| | | Magnetococcia | 1 | Magnetococcales | 1 |
| Bacteroidota | 173 | Flavobacteriia | 76 | Flavobacteriales | 76 |
| | | Chitinophagia | 60 | Chitinophagales | 60 |
| | | Sphingobacteriia | 16 | Sphingobacteriales | 16 |
| | | Cytophagia | 15 | Cytophagales | 15 |
| | | Bacteroidia | 6 | Marinilabiliales | 6 |
| Myxococcota | 74 | Polyangia | 41 | Nannocystales | 22 |
| | | | | Polyangiales | 19 |
| | | Myxococcia | 33 | Myxococcales | 33 |
| Acido-bacteriota | 69 | Terriglobia | 64 | Terriglobales | 64 |
| | | Holophagae | 4 | Holophagales | 3 |
| | | | | Acanthopleuribacterales | 1 |
| | | Vicinamibacteria | 1 | Vicinamibacterales | 1 |
| Plancto-mycetota | 54 | Planctomycetia | 54 | Planctomycetales | 29 |
| | | | | Pirellulales | 22 |
| | | | | Isosphaerales | 2 |
| | | | | Gemmatales | 1 |
| Spirochaetota | 52 | Spirochaetia | 52 | Leptospirales | 52 |
| Cyano-bacteriota | 42 | Cyanophyceae | 42 | Oscillatoriales | 13 |
| | | | | Coleofasciculales | 10 |
| | | | | Synechococcales | 10 |
| | | | | Acaryochloridales | 7 |
| | | | | Spirulinales | 2 |
| Actino-mycetota | 33 | Actinomycetes | 33 | Kitasatosporales | 16 |
| | | | | Streptosporangiales | 8 |
| | | | | Pseudonocardiales | 7 |
| | | | | Micrococcales | 1 |
| | | | | Mycobacteriales | 1 |
| Verruco-microbiota | 33 | Verrucomicrobiia | 19 | Verrucomicrobiales | 19 |
| | | Opitutia | 6 | Opitutales | 6 |
| | | Methylacidiphilae | 5 | Methylacidiphilales | 5 |
| | | Spartobacteria | 2 | Chthoniobacterales | 2 |
| | | Terrimicrobiia | 1 | Terrimicrobiales | 1 |
| Campylo-bacterota | 22 | Epsilon-proteobacteria | 22 | Campylobacterales | 22 |
| Bdello-vibrionota | 17 | Bacteriovoracia | 9 | Bacteriovoracales | 9 |
| | | Bdellovibrionia | 4 | Bdellovibrionales | 4 |
| | | Oligoflexia | 4 | Oligoflexales | 3 |
| | | | | Silvanigrellales | 1 |
| Chlamydiota | 11 | Chlamydiia | 11 | Parachlamydiales | 11 |
| Nitrospirota | 5 | Nitrospiria | 5 | Nitrospirales | 5 |
| Aquificota | 4 | Aquificia | 4 | Aquificales | 4 |
| Ignavi-bacteriota | 2 | Ignavibacteria | 2 | Ignavibacteriales | 2 |
| Candidatus Binatota | 2 | Candidatus Binatia | 2 | Candidatus Binatales | 2 |
| Gemmati-monadota | 1 | Gemmatimonadia | 1 | Gemmatimonadales | 1 |
| Calditrichota | 1 | Calditrichia | 1 | Calditrichales | 1 |

**Table S4. Proteins encoded in the *fon*, *hvf*, and *ngo* BGCs.**

| Name | Accession Code | PfamID | Pfam Annotation | Assigned function |
|---|---|---|---|---|
| FonA | SFF26396.1 | - | - | Lys-Cys-family 5-thiooxazole precursor peptide |
| FonB | SFF26386.1 | PF05114 | DUF692 | Multinuclear non-heme iron-dependent oxidative enzyme (MNIO) |
| FonC | SFF26371.1 | PF09836 | DUF2063 | Partner protein of MNIO enzyme, type 1 (PME1) |
| HvfX | WP_005653243.1 | PF07681 | DoxX | Unknown |
| HvfA | WP_005691265.1 | - | - | Lys-Cys-family 5-thiooxazole precursor peptide |
| HvfB | WP_032826769.1 | PF05114 | DUF692 | Multinuclear non-heme iron-dependent oxidative enzyme (MNIO) |
| HvfC | WP_005691267.1 | PF09836 | DUF2063 | Partner protein of MNIO enzyme, type 1 (PME1) |
| NgoX | WP_003692059.1 | PF07681 | DoxX | Unknown |
| NgoA | WP_003692062.1 | - | - | Lys-Cys-family 5-thiooxazole precursor peptide |
| NgoB | WP_010355671.1 | PF05114 | DUF692 | Multinuclear non-heme iron-dependent oxidative enzyme (MNIO) |
| NgoC | WP_003705168.1 | PF09836 | DUF2063 | Partner protein of MNIO enzyme, type 1 (PME1) |

**Table S5. Key gene sequences from the *fon*, *hvf*, and *ngo* BGCs.**

| Name | Sequence |
|---|---|
| *fonA* | ATGGACATTCCACGTAAACCCTTTGCAGTGGCGCTGGGCGCCGCGTTCGCACTCGGCACCGCCGCTGCCCAGGCGTCTGTGTTCCAGGCAT<br>CCGACCTCGGCGCCGGCTACATGGTCGCATCGGCTGGTGAGCACGCCAAATCCGGCGAGGCCAAATGTGGCGAGGGCAAGTGTGGCGAAAA<br>AAAGGCCGATGCCAAGGCCAAAGAGGGCAAGTGCGGCGAGGGTAAGTGCGGCGAAGGCAAGTGCGGTGAAAAGAAGGCCGACGCCAAGGCC<br>AACGAGGGCAAATGCGGCGAGGGCAAGTGCGGCGAAAACAAGACCCAGTGA |
| *fonA* (codon-optimized) | ATGGATATACCGCGTAAACCGTTCGCTGTAGCACTCGGGGCTGCATTTGCTCTTGGTACGGCAGCGGCGCAAGCATCCGTATTTCAAGCGA<br>GTGATCTTGGTGCAGGGTATATGGTTGCTTCTGCGGGCGAACATGCAAAGTCTGGTGAAGCTAAGTGCGGTGAAGGGAAATGCGGGGAGAA<br>GAAAGCTGACGCTAAAGCGAAGGAAGGTAAATGTGGGGAAGGAAAATGTGGTGAGGGTAAATGTGGCGAGAAGAAAGCGGATGCTAAAGCA<br>AATGAAGGAAAGTGTGGGGAAGGGAAATGTGGTGAGAATAAAACTCAATAA |
| *fonB* | ATGAGCCTGCACCCGACCATTGCCGGCGCCGGTCTTGGCCTGCGCCGGGCCTTGCTCGAACCACTGGGTGAATCTGCGCCCTGCGTCGATT<br>TCCTCGAGATCGCGCCGGAGAACTGGATGCGCGTCGGCGGGCGCCTGGGCCGTGCATTTCGCGCGCTGACCGAACGCTATCCGTTCGCGGC<br>GCACGGCCTTTCGCTGTCGATCGGCGCGCCGGCGCCGCTGGATGAGGATTTCGTCCGTGAGCTCAAGGCTTTCCTCGACCTGCATGGGATT<br>GCCGACTACAGCGAGCATCTGAGCTATTGTTCCGACGACGGGCATCTCTACGACTTGATGCCGATGCCGTTCACCGCCGAGGCGGTGGAGT<br>ACGTCGCCGCACGCGTGCGCCGCGTGCAGGACATCCTCGAGCGCCGCATTGCGCTGGAAAACATCAGCTATTACGCGCCGCTGTGCACCGA<br>ACTGACGGAACTGGAGTTCATCACGGCGGTGATCGAGCGCGCCGACTGCGATCTGCTGCTCGATGTCAACAACATCTACGTCAACAGCATC<br>AACCATCGCTACGACGCCCAGGCCTTTCTCGACGCATTGCCCGGCGAGCGCGTGCGCTGGATCCACGTCGCCGGTCATTACGTCGAGGCCG<br>AGGACTTGCGCGTGGATACACACGGCGCCGACGTGATCGATCCGGTGTGGGCCCTGCTCGACCGCGCCTATGCACGCTTCGGCGTGCGGCC<br>GACGCTGCTCGAGCGCGATTTCAACATTCCGCCGCTGGACGTGCTGCTCGGCGAAGTCGGGCACATCCGTGATATCCAGGCGCGGCATCGA<br>TCGCTGCAGCTCGCCTATGGCTAA |
| *fonB* (codon-optimized) | ATGTCCTTACATCCCACGATAGCAGGAGCAGGATTAGGGTTACGTCGTGCACTGTTGGAGCCCTTAGGCGAGAGCGCCCCTTGTGTTGACT<br>TTCTGGAAATTGCACCAGAAAATTGGATGCGTGTAGGTGGCCGGCTCGGAAGAGCTTTCCGAGCCTTAACGGAGCGTTACCCATTTGCCGC<br>TCATGGTTTGTCACTTTCAATTGGTGCCCCTGCCCCTTTAGACGAAGACTTTGTACGCGAATTAAAAGCGTTTCTTGATTTGCACGGTATA<br>GCAGATTATAGTGAACACTTGAGTTACTGCAGCGATGATGGTCACCTGTATGATCTTATGCCCATGCCATTTACAGCAGAAGCAGTCGAAT<br>ATGTGGCGGCCAGAGTTCGTCGTGTTCAAGATATTCTGGAACGTCGGATCGCTTTGGAGAATATATCCTACTATGCACCTCTTTGTACGGA<br>GCTCACCGAGTTGGAATTTATAACAGCTGTCATTGAACGAGCAGATTGTGACTTACTTCTGGACGTAAACAATATATATGTGAATTCGATT<br>AATCACCGTTATGATGCACAAGCGTTCCTGGATGCGCTTCCTGGTGAACGTGTTCGTTGGATTCATGTAGCAGGACACTATGTAGAAGCTG<br>AAGATCTTAGAGTTGACACGCATGGTGCGGATGTTATTGACCCTGTCTGGGCGTTGCTGGATAGAGCGTACGCCCGGTTTGGAGTTCGCCC<br>CACTCTCCTGGAAAGAGACTTTAATATCCCACCCCTCGATGTTTTGTTGGGAGAGGTGGGCCATATACGGGACATTCAAGCAAGACACCGC<br>TCATTGCAATTGGCGTACGGATGA |
| *fonC* | ATGGCTAACGATCAGCGCTGCGCGTCCGCAACTGAGCGCGACTGGCGTGCGCGCCAGCGCGCGTTCGCCGCGCATCTGCGCGACCCGCGCA<br>ATGCCTTCGCTGGCATCGAGGAGCGGCGGCTGCGGATTTACCGCGAGCTGTTCTACAACAATGTCGAGAGCTGCCTGGCCAGTGCATTCCC<br>GGTGCTACGCAAACTGTCGAGCGATGCGGTCTGGCATGCGCGCGTGCGCGATTTTTTCACGCGCCACCGCTGCACCGCACCGCAGTTCCAT<br>CGTCTGCCCGAGGAGTTCCTGCGCTATCTCGAGGCGGAGCGCGGCGAACACGCCGACGATCCGCCGTTCCTGCGCGAGCTCGCGCACTATG<br>AATGGGTCGAGCTCGCGCTCGCGATCGCGCCAGAGGATTTGCCCATCGAGCAGGCCGACCCGGAAGGTGATCTGCTCGCCGGATGCCCGCT<br>GCTGTCACCGCTGGCGTGGCCGCTGGTCTATGCCTATCCGGTGCACCGCATCGGGCCGCAATTCCAGCCGCAGGCACCCGGCGACTCGCCC<br>ACGTATCTGATCGTCAACCGCGACCGTGAAGACCGCGTCCGCTTCCTGGAGATCAATGCCGTCACCGCGCGTCTGGTCGCGCTGATCGAGG<br>CCGAACCCGCTGCGAGCGGACGTGCGCTGCTGCTGCGCATCGCTGCCGAACTCGCCCATCCCGATCCTCAATCCGTCGTCGCCCAGGGCGC<br>GCGCATCCTCGCCGATCTGCGCGCGCGCGACATCGTGCTCGGCACGCGGCGCTGA |
| *fonC* (codon-optimized) | ATGGCGAATGACCAACGATGTGCCAGTGCGACGGAAAGAGATTGGAGAGCCCGTCAACGGGCATTTGCGGCCCACTTACGTGATCCCAGAA<br>ACGCTTTTGCGGGAATTGAAGAAAGACGTCTTCGCATCTATCGAGAATTATTTTATAATAACGTTGAATCTTGTTTGGCTTCAGCCTTTCC<br>CGTACTGCGGAAGTTGAGCTCCGACGCAGTTTGGCACGCTCGTGTTCGGGACTTCTTTACCCGTCATCGATGTACTGCTCCTCAATTTCAC<br>CGCTTACCGGAAGAATTTCTTCGATACTTAGAAGCAGAACGTGGTGAGCATGCAGATGACCCGCCCTTTCTGCGTGAACTTGCACATTACG<br>AGTGGGTGGAATTGGCACTGGCAATTGCCCCTGAAGACTTACCGATTGAACAAGCTGATCCTGAGGGCGACTTGCTGGCGGGCTGTCCCCT<br>CTTATCCCCCTTGGCATGGCCCTTAGTTTACGCATACCCCGTTCATCGTATTGGTCCTCAGTTTCAACCTCAAGCTCCGGGTGATTCTCCA<br>ACCTACCTTATAGTAAATCGAGATCGCGAGGATAGAGTACGGTTCTTAGAAATTAACGCTGTAACTGCACGGCTCGTGGCTTTAATAGAAG<br>CGGAGCCTGCAGCTAGTGGTCGGGCTCTTTTGTTACGAATAGCGGCAGAGCTGGCGCACCCTGACCCCCAGAGCGTGGTAGCGCAAGGTGC<br>TAGAATTCTTGCTGACTTACGGGCTCGTGATATAGTTCTGGGAACTCGTAGATAA |
| *hvfA* | ATGAAAAAATTAGCAACATTAACAGCATTAGCAAGTGCATTAACGATGGCAGTAGCGACAGCAGTACAGGCTGAATCAAAGTCAAGCAATA<br>CTGATAATACAGCAACACCTTGTGTAGGCGATAAATGTGTCAAAACGAAAGCCGCAGAAGGTAAGTGTGGTGAAGGCAAATGCGGTGCAGA<br>TAAAGCTAAATCTGCTGAAGGTAAATGTGGCGAGGGTAAATGTGGTGCGAGCAAACCGAAAGCTGCTGAAGGCAAATGTGGCGAAGGTAAA<br>TGCGGTTCTAAATAA |
| *hvfA* (codon-optimized) | ATGAAAAAATTAGCTACCCTGACCGCATTGGCAGGAGCACTGACCATGGCGGTTGCAACGGCTGCTCAGGCCGAGTCTAAAAGCAGCTCAA<br>CTGATAATACCGCGACTCCATGTGTCGGCGACAAATGCGTCAAAACGAAAGCGGCTGAGGGTAAATGCGGTGAAGGAAAATGCGGAGCGGA<br>CAAGGCGAAAAGCGCCGAGGGAAAATGTGGAGAAGGAAAGTGTGGAGCATCTAAACCTAAGGCGGCGGAAGGGAAATGCGGAGAGGGAAAA<br>TGCGGCTCAAAATAA |
| *hvfB* | ATGAAATTACAAGGCGCTGGACTAGGTTATCGTCGGAATTTAGCTGAGGATTTTTTACAACTTCCCTCAAATAACGCTATTCAATTTATTG<br>AAGTCGCACCAGAAAATTGGAGTAAAATGGGTGGAATGGCACGTTATCAATTTGATCAAGCAGCAGAAAGATTTCCGTTAGCAGTACACGG<br>TCTTTCACTTTCTTTAGGTGGGCAAGCACCACTTGATCGCGAATTACTCCGTAATACGAAAGCATTAATTAATCAATATAATTCGTCTTTC<br>TTTTCTGAACATTTAAGTTATTGTGAATGTGAAGGGCATTTATATGATTTATTACCTATGCCATTTACAGAAGAAGCAGTAAAACACGTTG<br>CACAACGAATACGTGATGTGCAAGATTTCTTAGGATTACAAATTTCATTAGAAAATACTTCTTATTATTTGCATTCTCCTACTAGCACAAT<br>GAATGAAGTGGAATTTTTAAATGCTATCGCACAAGAGGCTGATTGTGGCATTCATTTAGATGTGAATAATATTTATGTTAATGGTGTTAAT<br>CACGGATTGCTTGATCCTTATATCTTTTTAGATCAAGTTGATGTTAAACGTGTTAATTACATTCATATTGCAGGGCACGATGAAGAACATT<br>CTGCTGCACAAGTTGTAGAGAATTCAGCAAATGAATCATTTAATAAAGTAAAAGGAGCATATCGCCATTTACCTGAATTATTAATTGATAC<br>ACACGGAGAAGCTGTAAAAGGCACTGTATGGGATTTACTCGAATATGCCTATCAACGACTACCTACGATTCCTCCAACATTATTGGAGCGT<br>GATTTCAACTTCCCACCATTTGCAGAACTTTACGCCGAAGTTGAGCATATTGCACAATTACAGCAAAAATACGCTCACACAGAGGTAATGT<br>CTTATGCAGCCTAA |
| *hvfB* (codon-optimized) | ATGAAATTACAAGGCGCCGGATTAGGATATCGCCGTAACTTGGCGGAAGACTTTCTGCAGCTGCCTAGCAATAACGCGATCCAATTCATCG<br>AAGTCGCACCTGAGAACTGGAGTAAAATGGGCGGAATGGCACGCTACCAGTTCGACCAGGCGGCAGAGCGCTTTCCGCTGGCAGTTCATGG<br>GCTTAGTCTGTCCTTAGGGGGGCAAGCTCCCTTAGACCGTGAATTGCTTCGCAATACTAAGGCACTTATTAACCAGTACAACTCCTCTTTT<br>TTCTCGGAGCACCTGTCCTATTGTGAGTGTGAAGGCCACTTGTACGACCTTCTGCCCATGCCATTCACTGAGGAAGCGGTGAAGCATGTTG<br>CACAACGCATCCGCGATGTGCAGGATTTCTTGGGGCTTCAAATTAGTCTTGAGAATACATCCTATTATTTGCACTCCCCAACGTCAACCAT |

| | |
|---|---|
| *hvfB* (codon-optimized, cont.) | GAATGAAGTTGAATTTCTTAACGCAATCGCACAAGAGGCCGACTGTGGGATCCACTTGGATGTTAACAATATTTATGTAAATGGCGTGAAC<br>CACGGGTTACTTGATCCATATATCTTCTTGGACCAGGTGGATGTGAAGCGTGTAAATTACATCCACATCGCTGGTCATGACGAAGAGCACT<br>CAGCCGCACAGGTAGTAGAAAATAGTGCCAATGAATCCTTCAACAAAGTAAAAGGCGCTTATCGTCATCTGCCAGAGCTGTTAATTGACAC<br>ACATGGGGAAGCCGTTAAAGGCACGGTTTGGGATTTACTGGAATATGCTTACCAGCGTTTGCCTACCATCCCTCCAACATTATTAGAGCGT<br>GATTTTAACTTCCCTCCATTCGCAGAACTTTACGCGGAGGTAGAGCATATTGCACAGCTTCAGCAAAAATACGCACATACAGAGGTCATGA<br>GCTACGCCGCCTGA |
| *hvfC* | ATGCAGCCTAAGTCATCATTGAAAGAAACTCAGCAAGCATTGGCAAATGCTATTCGGTTAGGTAATGCAGATCCTTTAAATGGTTATGCTG<br>AAAGCCGTTTAGCGGTATATACACGTTTAGTTCGTAATAATGCTTTTGGTTTTATTGATCGTTGTTTTGTTGAAGCACCATTGCATATTGA<br>GCTAGAGTATTGGAAAAATGCTAAAGAAAATTTTGTGCAAAATGGTAATGCCCATTCTCCTTATTTCCAAGATATTGCAGGAGAATTTCTC<br>TTGTTTTGTCAAGAAAAAGAGATATTTGATACAAATATATTGGCATTGATGGATTTTGAAAATACACAATTACTCGCAGAAGTTTCTTTAG<br>CCAAAGTGCCTGAAAAATTTGAGTGGAATAGACATAGCGTAATGCAATTATCTGGTGCAGCTTATTTAAAAAGTTATGAAGTTGATTTTTT<br>ATCAAGTGACTTTAAACAATTTGATGATACGCCAATTCAAGTTATTATATGGCGTGATAGTGATTTTAGGATTCAGCAACAAATCCTTTCA<br>GAGTTAGATTACTGGTTGTTAAGTTATCTACAAGAACAACCAAATTCATTAGAAAATGTTTTGTCTGCTCTTAATACAATGGTTGAAGATA<br>GTACGTCAATAATCCCATTATTAGAGCAAGTATGGATGAAATGGGTTACATCTGAAGTAATTTACCCCGAACAAAGATAA |
| *hvfC* (codon-optimized) | ATGCTGCCCAAGTCTAGCTTGAAAGAGACGCAGCAAGCATTGGCCAACGCGATCCGCCTTGGCAATGCTGATCCGCTGAATGGATATGCAG<br>CTTCCCGCTTAGCGGTCTATACCCGTTTGGTACGTAACAACGCTTTTGGGTTTATCGACCGCTGTTTTGTGGAAGCCCCGTTACACATCGA<br>ACCCGAGTATTGGAAAAATGCGAAAGAGAACTTTGTTCAAAACGGCAATGCACACTCTCCGTACTTTCAAGATATCGCCGGGGAATTCTTA<br>TTGCTTTGTCAAGAGAAAGAAATCTTTGACACTAATATTCTGGCGTTGATGGACTTCGAGAACACACAATTGTTGGCTGAGGTATCTCTTG<br>CAAAGGTACCGGAGAAGTTCGAGTGGAACCGCCATAGCGTGATGCAACTGTCTGGCGCCGCATATTTAAAGAGCTACGACGTGGACTTCTT<br>ATCATCCGACTTCAAGCAGTTCGACGATACCCCCATCCAAGCCATTATCTGGCGCGATTCCGATTTCCGCATTCAACAACAAATTCTGTCG<br>GAACTTGACTACTGGCTGTTGTCATACTTACAGGAACAACCTAATAGTTTGGAAAATGTACTTAGCGCTCTGAACACTATGGTTGAAGATT<br>CGACAAGTATTGTCCCCCTGTTAGAGCAGGTGTGGATGAAGTGGGTAACCTCCGAGGTGATTTACCCTGAACAGCGCTGA |
| *ngoA* | ATGAACAAAAATATTGCTGCCGCACTCGCCGGTGCTTTATCCCTGTCTCTGGCCGCCGGCGCCGTTGCCGCCCACAAACCGGCAAGCAACG<br>CAACAGGCGTTCAAAAATCCGCCCAAGGCTCTTGCGGCGCATCCAAATCTGCCGAAGGTTCGTGCGGCGCATCCAAATCTGCCGAAGGTTC<br>GTGCGGCGCGGCTGCTTCTAAAGCAGGCGAAGGCAAATGCGGCGAGGGCAAATGCGGTGCAACTGTAAAAAAAGCCCACAAACACACCAAA<br>GCATCTAAAGCCAAAGCCAAATCTGCCGAAGGCAAATGCGGCGAAGGCAAATGCGGTTCTAAATAA |
| *ngoA* (codon-optimized) | ATGAACAAGAACATTGCAGCAGCTTTAGCAGGAGCTTTGTCCTTATCTTTAGCCGCCGGCGCCGTGGCTGCGCACAAACCCGCATCTAACG<br>CGACTGGCGTTCAAAAGAGCGCCCAAGGGTCGTGCGGTGCATCTAAAAGTGCTGAAGGTAGCTGCGGGGCCAGTAAATCAGCTGAGGGCTC<br>CTGCGGGGCGGCCGCCAGCAAAGCAGGTGAGGGTAAATGCGGAGAGGGCAAATGCGGAGCAACGGTGAAGAAAGCCCACAAACACACCAAA<br>GCGTCAAAAGCAAAAGCCAAATCTGCCGAAGGCAAATGCGGGGAAGGAAAATGCGGCTCTAAATAA |
| *ngoB* | ATGATTCAACACGCAGGCTTGGGCTACCGCCGCGACTTGGCGGAAGACTTTCTCTCGCTTTCCGAAAACAGCCCGATATGCTTTATCGAAG<br>CCGCACCGGAAAACTGGCTGAAAATGGGCGGCAGGGCGCGCAAACAGTTTGACCGTGTGGCGGAACGGCTGCCGCTGGCGTTGCACGGATT<br>ATCTATGTCGCTGGGCGGACAAGCCCCGCTGGATACCGATTTGATAGACGGCATCAAAGAAATGATGTGCCGTTATGACTGCACGTTTTTC<br>TCCGATCATTTGAGCTACTGCCACGACGGCGGTCATCTTTACGATTTGTTGCCGCTGCCTTTCACTGAAGAAATGGTGCATCATACGGCGC<br>GGCGTATCCGCGAAGTGCAAGACCGTTTGGGCTGCCGCATCGCCGTGGAAAACACGTCCTACTACCTGCATTCTCCGCTTGCCGAGATGAA<br>CGAGGTCGAGTTCCTCAACGCGGTTGCCCGTGAAGCCGACTGCGGTATTCATTTGGATGTGAACAATATTTACGTCAACGCCGTCAATCAC<br>GGCCTGCTGTCGCCTGAGGCCTTTTTAGAGAATGTGGACGCAGGGCGCGTGTGCTACATCCACATCGCCGGACACGATGCCGAAACCCCCG<br>AGCTTTTGATTGATACCCACGGCGCGGCGGTTTTACCGACCGTTTGGGACTTGCTCGAACTCGCCTACACCAAGCTTCCCACCATCCCTCC<br>CACCCTGTTGGAACGCGATTTCAATTTCCCACCTTTTGCCGAACTCGAAGCCGAAGTTGCCAAAATTGCCGATTATCAAACGCGTGCCGGA<br>AAGGAATACCGCCGTGCAGCCTGA |
| *ngoB* (codon-optimized) | ATGATACAGCATGCTGGATTAGGGTATCGTCGGGACTTAGCCGAGGATTTCCTTAGTTTAAGCGAGAATTCGCCTATTTGTTTCATTGAGG<br>CGGCGCCTGAGAATTGGTTAAAGATGGGCGGTCGTGCACGGAAGCAATTCGATCGAGTCGCCGAGAGATTACCACTTGCATTACATGGTCT<br>CTCAATGTCCTTAGGTGGTCAGGCACCACTTGACACGGACCTGATTGATGGGATAAAGGAGATGATGTGTCGGTACGATTGTACTTTCTTT<br>TCAGACCACCTGAGTTATTGTCATGATGGCGGCCACTTGTATGACTTACTCCCCCTTCCATTTACGGAAGAGATGGTACACCACACAGCCC<br>GACGGATACGTGAGGTCCAGGATCGACTGGGGTGTCGTATTGCAGTAGAGAATACCAGCTATTATTTGCACTCGCCCTTAGCAGAAATGAA<br>TGAAGTGGAATTTCTCAATGCCGTCGCAAGAGAGGCAGATTGTGGAATCCACCTGGACGTCAATAACATCTATGTGAATGCGGTGAACCAT<br>GGACTTCTCTCTCCAGAAGCATTCCTCGAAAACGTCGATGCCGGAAGAGTTTGTTATATTCATATAGCGGGTCATGACGCAGAGACACCGG<br>AACTGCTCATCGACACGCATGGAGCCGCAGTACTGCCTACGGTATGGGATCTGCTGGAGCTGGCGTATACAAAACTGCCGACAATTCCCCC<br>GACTTTGCTGGAGCGGGACTTTAACTTTCCGCCGTTCGCGGAGTTAGAGGCAGAGGTGGCTAAGATAGCAGACTACCAGACCAGAGCGGGC<br>AAAGAGTATCGTCGCGCCGCTTAA |
| *ngoC* | GTGCAGCCTGAAACCTCCGCCCAATACCAGCACCGTTTCTCCCAAGCCATACGCGGGGGCGAAGCCGCAGACGGTCTGCCGCAAGACCGAT<br>TGAACGTCTATATCCGCCTGATACGCAACAATATCCACAGCTTTATCGACCGCTGCTATACCGAAACGCGGCAATACTTTGACAGCAAAGA<br>ATGGAGCCGTCTGAAAGAAGGTTTCGTCCGCGACGCGCGTGCCCAAACGCCCTATTTTCAAGAAATCCCCGGCGAGTTCCTACAATATTGC<br>CAAAGCCCGCCGCTTTCAGACGGCATTTTGGCGCTGATGGATTTTGAATATACCCAATTGCTGGCAGAAGTTGCACAAATTCCGGATATTC<br>CCGACATTCATTATTCAAATGACAGCAAATACACGCCCTCCCCTGCGGCCTTTATCCGGCAATATCGATACGATGTTACCCATGATTTGCA<br>GGAAGCGGAAACAGCCTTGTTAATATGGCGAAACGCCGAAGATGATGTGATGTACCAAACATTGGACGGCTTCGATATGATGCTGCTGGAG<br>ATAATGGGTTCCTCCGCGCTTTCGTTTGACACCCTCGCCCAAACCCTTGTCGAATTTATGCCTAAAGCCGATAATTGGAAAAATATTTTGC<br>TTGGGAAATGGTCAGGCTGGATTGAACAAAGGATTATCATCCCCTCCTTGTCCGCCATATCCGAAAATATGGAAGGCAATTCCCCGAGCCA<br>AAACCATCTATCCGCATAA |
| *ngoC* (codon-optimized) | ATGCAACCGGAGACGTCAGCTCAGTATCAACATCGCTTTAGTCAGGCTATTAGAGGCGGAGAGGCAGCCGATGGCTTGCCACAGGATCGCC<br>TGAATGTTTACATTAGACTTATTCGTAATAACATTCATTCTTTCATTGATCGGTGTTACACAGAGACCCGTCAGTATTTCGATTCGAAGGA<br>GTGGTCCCGCCTCAAAGAGGGGTTTGTAAGAGATGCCCGAGCGCAGACACCGTACTTCCAGGAGATACCGGGTGAATTTCTCCAGTACTGT<br>CAGTCTCCCCCTTTATCTGATGGGATCCTGGCACTCATGGACTTCGAGTACACGCAGCTGTTGGCGGAAGTCGCCCAGATCCCAGACATAC<br>CAGATATACACTACAGCAACGATTCTAAGTATACACCTTCGCCCGCCGCTTTCATTCGACAGTACCGCTATGACGTCACGCACGACCTTCA<br>AGAGGCAGAGACTGCGCTGCTGATTTGGCGGAATGCGGAAGACGACGTCATGTATCAGACCTTAGATGGTTTTGACATGATGTTGCTCGAA<br>ATTATGGGCAGCAGTGCTCTCTCATTCGATACACTTGCTCAGACTTTAGTTGAGTTCATGCCAAAGGCAGACAACTGGAAGAACATACTCT<br>TGGGTAAGTGGTCTGGATGGATCGAGCAGCGAATCATTATTCCTAGCCTGTCTGCAATCTCAGAGAACATGGAAGGTAACTCTCCATCACA<br>GAATCACTTAAGCGCGTGA |

**Table S6. Key protein sequences from the *fon*, *hvf*, and *ngo* BGCs.**

| Name | Sequence |
|---|---|
| FonA | MDIPRKPFAVALGAAFALGTAAAQASVFQASDLGAGYMVASAGEHAKSGEAKCGEGKCGEKKADAKAKEGKCGEGKCGEG<br>KCGEKKADAKANEGKCGEGKCGENKTQ |
| FonB | MSLHPTIAGAGLGLRRALLEPLGESAPCVDFLEIAPENWMRVGGRLGRAFRALTERYPFAAHGLSLSIGAPAPLDEDFVR<br>ELKAFLDLHGIADYSEHLSYCSDDGHLYDLMPMPFTAEAVEYVAARVRRVQDILERRIALENISYYAPLCTELTELEFIT<br>AVIERADCDLLLDVNNIYVNSINHRYDAQAFLDALPGERVRWIHVAGHYVEAEDLRVDTHGADVIDPVWALLDRAYARFG<br>VRPTLLERDFNIPPLDVLLGEVGHIRDIQARHRSLQLAYG |
| FonC | MANDQRCASATERDWRARQRAFAAHLRDPRNAFAGIEERRLRIYRELFYNNVESCLASAFPVLRKLSSDAVWHARVRDFF<br>TRHRCTAPQFHRLPEEFLRYLEAERGEHADDPPFLRELAHYEWVELALAIAPEDLPIEQADPEGDLLAGCPLLSPLAWPL<br>VYAYPVHRIGPQFQPQAPGDSPTYLIVNRDREDRVRFLEINAVTARLVALIEAEPAASGRALLLRIAAELAHPDPQSVVA<br>QGARILADLRARDIVLGTRR |
| HvfA | MKKLATLTALAGALTMAVATAAQAESKSSSTDNTATPCVGDKCVKTKAAEGKCGEGKCGADKAKSAEGKCGEGKCGASKP<br>KAAEGKCGEGKCGSK |
| HvfB | MKLQGAGLGYRRNLAEDFLQLPSNNAIQFIEVAPENWSKMGGMARYQFDQAAERFPLAVHGLSLSLGGQAPLDRELLRNT<br>KALINQYNSSFFSEHLSYCECEGHLYDLLPMPFTEEAVKHVAQRIRDVQDFLGLQISLENTSYYLHSPTSTMNEVEFLNA<br>IAQEADCGIHLDVNNIYVNGVNHGLLDPYIFLDQVDVKRVNYIHIAGHDEEHSAAQVVENSANESFNKVKGAYRHLPELL<br>IDTHGEAVKGTVWDLLEYAYQRLPTIPPTLLERDFNFPPFAELYAEVEHIAQLQQKYAHTEVMSYAA |
| HvfC | MLPKSSLKETQQALANAIRLGNADPLNGYAASRLAVYTRLVRNNAFGFIDRCFVEAPLHIEPEYWKNAKENFVQNGNAHS<br>PYFQDIAGEFLLLCQEKEIFDTNILALMDFENTQLLAEVSLAKVPEKFEWNRHSVMQLSGAAYLKSYDVDFLSSDFKQFD<br>DTPIQAIIWRDSDFRIQQQILSELDYWLLSYLQEQPNSLENVLSALNTMVEDSTSIVPLLEQVWMKWVTSEVIYPEQR |
| NgoA | MNKNIAAALAGALSLSLAAGAVAAHKPASNATGVQKSAQGSCGASKSAEGSCGASKSAEGSCGAAASKAGEGKCGEGKCG<br>ATVKKAHKHTKASKAKAKSAEGKCGEGKCGSK |
| NgoB | MIQHAGLGYRRDLAEDFLSLSENSPICFIEAAPENWLKMGGRARKQFDRVAERLPLALHGLSMSLGGQAPLDTDLIDGIK<br>EMMCRYDCTFFSDHLSYCHDGGHLYDLLPLPFTEEMVHHTARRIREVQDRLGCRIAVENTSYYLHSPLAEMNEVEFLNAV<br>AREADCGIHLDVNNIYVNAVNHGLLSPEAFLENVDAGRVCYIHIAGHDAETPELLIDTHGAAVLPTVWDLLELAYTKLPT<br>IPPTLLERDFNFPPFAELEAEVAKIADYQTRAGKEYRRAA |
| NgoC | MQPETSAQYQHRFSQAIRGGEAADGLPQDRLNVYIRLIRNNIHSFIDRCYTETRQYFDSKEWSRLKEGFVRDARAQTPYF<br>QEIPGEFLQYCQSPPLSDGILALMDFEYTQLLAEVAQIPDIPDIHYSNDSKYTPSPAAFIRQYRYDVTHDLQEAETALLI<br>WRNAEDDVMYQTLDGFDMMLLEIMGSSALSFDTLAQTLVEFMPKADNWKNILLGKWSGWIEQRIIIPSLSAISENMEGNS<br>PSQNHLSA |

**Table S7. Vectors used in this study.**

| Name | Source | Resistance Marker |
| --- | --- | --- |
| pET-28a-*fonABC* | this work | Kanamycin |
| pET-28a-*hvfABC* | this work | Kanamycin |
| pET-28a-*ngoABC* | this work | Kanamycin |
| pET-28a-(MBP)*fonABC* | this work | Kanamycin |
| pET-28a-*fonBC* | this work | Kanamycin |
| pET-28a-*fonAB*(F250D)*C* | this work | Kanamycin |
| pET-28a-*fonB*(F250D)*C* | this work | Kanamycin |
| pET-28a-*fonAB*(F250A)*C* | this work | Kanamycin |
| pET-28a-*fonB*(F250A)*C* | this work | Kanamycin |
| pET-28a-*fonAB*(D249A)*C* | this work | Kanamycin |
| pET-28a-*fonB*(D249A)*C* | this work | Kanamycin |
| pET-28a-*fonAB*(R15A)*C* | this work | Kanamycin |
| pET-28a-*fonB*(R15A)*C* | this work | Kanamycin |
| pET-28a-*fonAB*(R16A)*C* | this work | Kanamycin |
| pET-28a-*fonB*(R16A)*C* | this work | Kanamycin |
| pET-28a-*fonAB*(R15A,R16A)*C* | this work | Kanamycin |

**Table S8. Table of daughter ion assignments for HR-MS/MS analysis of alkylated FonA$_{67\text{-}80}$.**

| Fragment Type | Fragmented Bond Number | Fragment Charge | Experimental *m/z* | Theoretical *m/z* | Mass Error (ppm) |
|---|---|---|---|---|---|
| b | 3 | 1 | 274.1032 | 274.1034 | -0.6 |
| b | 7 | 1 | 686.2559 | 686.2563 | -0.6 |
| b | 8 | 1 | 815.2990 | 815.2989 | 0.2 |
| b | 12 | 2 | 607.2215 | 607.2217 | -0.4 |
| b | 13 | 2 | 671.7426 | 671.743 | -0.6 |
| y | 2 | 1 | 276.1553 | 276.1554 | -0.3 |
| y | 3 | 1 | 333.1767 | 333.1769 | -0.5 |
| y | 5 | 1 | 617.271 | 617.2712 | -0.3 |
| y | 6 | 2 | 337.6498 | 337.6500 | -0.5 |
| y | 6 | 1 | 674.2923 | 674.2927 | -0.5 |
| y | 7 | 1 | 803.3352 | 803.3353 | -0.1 |
| y | 8 | 2 | 430.6817 | 430.682 | -0.7 |
| y | 8 | 1 | 860.3568 | 860.3567 | 0.1 |
| y | 10 | 2 | 572.7288 | 572.7292 | -0.7 |
| y | 11 | 3 | 405.8339 | 405.8342 | -0.9 |
| y | 11 | 2 | 608.2473 | 608.2477 | -0.7 |
| y | 12 | 3 | 448.8482 | 448.8484 | -0.5 |
| y | 12 | 2 | 672.7686 | 672.769 | -0.6 |
| y | 13 | 3 | 467.8553 | 467.8556 | -0.6 |
| y | 13 | 2 | 701.2794 | 701.2798 | -0.5 |

**Table S9. NMR assignments for alkylated FonA$_{67\text{-}80}$.**

| Number | Residue | Amide NH | αH | αC | βH | βC | γH | γC | δH | δC | εH | εC |
|---|---|---|---|---|---|---|---|---|---|---|---|---|
| **67** | **Ser** | N/A | 4.08 | 54.4 | 3.91 | 60.2 | | | | | | |
| **68** | **Gly** | 8.6-8.7 | 3.8-4.1 | 42.5 | | | | | | | | |
| **69** | **Glu** | 8.34 | 4.1-4.2 | 54 | 1.89, 2.01 | 26.7 | 2.3 | 31.6 | | 179.2 | | |
| **70** | **Ala** | 8.28 | 4.15 | 49.9 | 1.24 | 16.3 | | | | | | |
| **71** | **Lys** | 8.55 | 4.91 | 47.3 | 1.82 | 30.8 | 1.53 | 21.9 | 1.25 | 26.1 | 2.84 | 39.1 |
| **73** | **Gly** | 8.4-8.5 | 3.8-4.1 | 42.5 | | | | | | | | |
| **74** | **Glu** | 8.48 | 4.1-4.2 | 54 | 1.84, 2.00 | 26.7 | 2.3 | 31.6 | | 179.2 | | |
| **75** | **Gly** | 8.6-8.7 | 3.82, 3.94 | 42.5 | | | | | | | | |
| **76** | **Lys** | 8.19 | 4.96 | 47.3 | 1.82 | 30.8 | 1.53 | 21.9 | 1.25 | 26.1 | 2.84 | 39.1 |
| **78** | **Gly** | 8.4-8.5 | 3.8-4.1 | 42.5 | | | | | | | | |
| **79** | **Glu** | 8.3 | 4.23 | 54 | 1.84, 2.00 | 26.7 | 2.25 | 31.6 | | 180.2 | | |
| **80** | **Lys** | 8.01 | 4.05 | 54.9 | 1.69 | 30.8 | 1.53 | 21.9 | 1.25 | 26.1 | 2.84 | 39.1 |

| | | C1 (Lys C=O) | C2 (Cys βC) | C3 (Cys αC) | C4 (Cys C=O) | C5 (acetamide $CH_2$) | C5-$H_2$ | C6 (acetamide C=O) | C6-$NH_2$ |
|---|---|---|---|---|---|---|---|---|---|
| **72** | **S-acetamido - 5-thiooxazole** | 163 | 149 | 131 | 162 | 35 | 3.67 | 173 | 7.1, 7.7 |
| **77** | **S-acetamido - 5-thiooxazole** | 163 | 149 | 131 | 162 | 35 | 3.67 | 173 | 7.1, 7.7 |

**Table S10. Constructs generated for *in vitro* transcription/translation.** Bolded letters indicate variant residues.

| Name | Peptide sequence | DNA template |
|---|---|---|
| FonA$_{PE}$_WT | MQASVFQASDLGAG<br>YMVASAGEHAKSGE<br>AKCGEGKCGEKKAD | GGCGTAATACGACTCACTATAGGGTTAACTTTAACAAGGAGAAAAACATGCAGGCGAGCGT<br>GTTTCAGGCCAGCGATCTGGGCGCGGGTTATATGGTGGCGAGCGCGGGCGAACATGCCAAA<br>AGCGGCGAAGCGAAATGCGGCGAAGGCAAATGCGGCGAAAAAAAAGCGGATTAAGCTTCG |
| FonA$_{PE}$Q(-26)A | M**A**ASVFQASDLGAG<br>YMVASAGEHAKSGE<br>AKCGEGKCGEKKAD | GGCGTAATACGACTCACTATAGGGTTAACTTTAACAAGGAGAAAAACATGGCGGCGAGCGT<br>GTTTCAGGCCAGCGATCTGGGCGCGGGTTATATGGTGGCGAGCGCGGGCGAACATGCCAAA<br>AGCGGCGAAGCGAAATGCGGCGAAGGCAAATGCGGCGAAAAAAAAGCGGATTAAGCTTCG |
| FonA$_{PE}$A(-25)G | MQ**G**SVFQASDLGAG<br>YMVASAGEHAKSGE<br>AKCGEGKCGEKKAD | GGCGTAATACGACTCACTATAGGGTTAACTTTAACAAGGAGAAAAACATGCAGGGCAGCGT<br>GTTTCAGGCCAGCGATCTGGGCGCGGGTTATATGGTGGCGAGCGCGGGCGAACATGCCAAA<br>AGCGGCGAAGCGAAATGCGGCGAAGGCAAATGCGGCGAAAAAAAAGCGGATTAAGCTTCG |
| FonA$_{PE}$S(-24)A | MQA**A**VFQASDLGAG<br>YMVASAGEHAKSGE<br>AKCGEGKCGEKKAD | GGCGTAATACGACTCACTATAGGGTTAACTTTAACAAGGAGAAAAACATGCAGGCGGCGGT<br>GTTTCAGGCCAGCGATCTGGGCGCGGGTTATATGGTGGCGAGCGCGGGCGAACATGCCAAA<br>AGCGGCGAAGCGAAATGCGGCGAAGGCAAATGCGGCGAAAAAAAAGCGGATTAAGCTTCG |
| FonA$_{PE}$V(-23)A | MQAS**A**FQASDLGAG<br>YMVASAGEHAKSGE<br>AKCGEGKCGEKKAD | GGCGTAATACGACTCACTATAGGGTTAACTTTAACAAGGAGAAAAACATGCAGGCGAGCGC<br>GTTTCAGGCCAGCGATCTGGGCGCGGGTTATATGGTGGCGAGCGCGGGCGAACATGCCAAA<br>AGCGGCGAAGCGAAATGCGGCGAAGGCAAATGCGGCGAAAAAAAAGCGGATTAAGCTTCG |
| FonA$_{PE}$F(-22)A | MQASV**A**QASDLGAG<br>YMVASAGEHAKSGE<br>AKCGEGKCGEKKAD | GGCGTAATACGACTCACTATAGGGTTAACTTTAACAAGGAGAAAAACATGCAGGCGAGCGT<br>GGCGCAGGCCAGCGATCTGGGCGCGGGTTATATGGTGGCGAGCGCGGGCGAACATGCCAAA<br>AGCGGCGAAGCGAAATGCGGCGAAGGCAAATGCGGCGAAAAAAAAGCGGATTAAGCTTCG |
| FonA$_{PE}$Q(-21)A | MQASVF**A**ASDLGAG<br>YMVASAGEHAKSGE<br>AKCGEGKCGEKKAD | GGCGTAATACGACTCACTATAGGGTTAACTTTAACAAGGAGAAAAACATGCAGGCGAGCGT<br>GTTTGCGGCCAGCGATCTGGGCGCGGGTTATATGGTGGCGAGCGCGGGCGAACATGCCAAA<br>AGCGGCGAAGCGAAATGCGGCGAAGGCAAATGCGGCGAAAAAAAAGCGGATTAAGCTTCG |
| FonA$_{PE}$A(-20)G | MQASVFQ**G**SDLGAG<br>YMVASAGEHAKSGE<br>AKCGEGKCGEKKAD | GGCGTAATACGACTCACTATAGGGTTAACTTTAACAAGGAGAAAAACATGCAGGCGAGCGT<br>GTTTCAGGGCAGCGATCTGGGCGCGGGTTATATGGTGGCGAGCGCGGGCGAACATGCCAAA<br>AGCGGCGAAGCGAAATGCGGCGAAGGCAAATGCGGCGAAAAAAAAGCGGATTAAGCTTCG |
| FonA$_{PE}$S(-19)A | MQASVFQA**A**DLGAG<br>YMVASAGEHAKSGE<br>AKCGEGKCGEKKAD | GGCGTAATACGACTCACTATAGGGTTAACTTTAACAAGGAGAAAAACATGCAGGCGAGCGT<br>GTTTCAGGCCGCGGATCTGGGCGCGGGTTATATGGTGGCGAGCGCGGGCGAACATGCCAAA<br>AGCGGCGAAGCGAAATGCGGCGAAGGCAAATGCGGCGAAAAAAAAGCGGATTAAGCTTCG |
| FonA$_{PE}$D(-18)A | MQASVFQAS**A**LGAG<br>YMVASAGEHAKSGE<br>AKCGEGKCGEKKAD | GGCGTAATACGACTCACTATAGGGTTAACTTTAACAAGGAGAAAAACATGCAGGCGAGCGT<br>GTTTCAGGCCAGCGCGCTGGGCGCGGGTTATATGGTGGCGAGCGCGGGCGAACATGCCAAA<br>AGCGGCGAAGCGAAATGCGGCGAAGGCAAATGCGGCGAAAAAAAAGCGGATTAAGCTTCG |
| FonA$_{PE}$L(-17)A | MQASVFQASD**A**GAG<br>YMVASAGEHAKSGE<br>AKCGEGKCGEKKAD | GGCGTAATACGACTCACTATAGGGTTAACTTTAACAAGGAGAAAAACATGCAGGCGAGCGT<br>GTTTCAGGCCAGCGATGCAGGCGCGGGTTATATGGTGGCGAGCGCGGGCGAACATGCCAAA<br>AGCGGCGAAGCGAAATGCGGCGAAGGCAAATGCGGCGAAAAAAAAGCGGATTAAGCTTCG |
| FonA$_{PE}$G(-16)A | MQASVFQASDL**A**AG<br>YMVASAGEHAKSGE<br>AKCGEGKCGEKKAD | GGCGTAATACGACTCACTATAGGGTTAACTTTAACAAGGAGAAAAACATGCAGGCGAGCGT<br>GTTTCAGGCCAGCGATCTGGCAGCGGGTTATATGGTGGCGAGCGCGGGCGAACATGCCAAA<br>AGCGGCGAAGCGAAATGCGGCGAAGGCAAATGCGGCGAAAAAAAAGCGGATTAAGCTTCG |
| FonA$_{PE}$A(-15)G | MQASVFQASDLG**G**G<br>YMVASAGEHAKSGE<br>AKCGEGKCGEKKAD | GGCGTAATACGACTCACTATAGGGTTAACTTTAACAAGGAGAAAAACATGCAGGCGAGCGT<br>GTTTCAGGCCAGCGATCTGGGCGGTGGTTATATGGTGGCGAGCGCGGGCGAACATGCCAAA<br>AGCGGCGAAGCGAAATGCGGCGAAGGCAAATGCGGCGAAAAAAAAGCGGATTAAGCTTCG |
| FonA$_{PE}$G(-14)A | MQASVFQASDLGA**A**<br>YMVASAGEHAKSGE<br>AKCGEGKCGEKKAD | GGCGTAATACGACTCACTATAGGGTTAACTTTAACAAGGAGAAAAACATGCAGGCGAGCGT<br>GTTTCAGGCCAGCGATCTGGGCGCGGCATATATGGTGGCGAGCGCGGGCGAACATGCCAAA<br>AGCGGCGAAGCGAAATGCGGCGAAGGCAAATGCGGCGAAAAAAAAGCGGATTAAGCTTCG |
| FonA$_{PE}$Y(-13)A | MQASVFQASDLGAG<br>**A**MVASAGEHAKSGE<br>AKCGEGKCGEKKAD | GGCGTAATACGACTCACTATAGGGTTAACTTTAACAAGGAGAAAAACATGCAGGCGAGCGT<br>GTTTCAGGCCAGCGATCTGGGCGCGGGTGCAATGGTGGCGAGCGCGGGCGAACATGCCAAA<br>AGCGGCGAAGCGAAATGCGGCGAAGGCAAATGCGGCGAAAAAAAAGCGGATTAAGCTTCG |
| FonA$_{PE}$E1A | MQASVFQASDLGAG<br>YMVASAGEHAKSG**A**<br>AKCGEGKCGEKKAD | GGCGTAATACGACTCACTATAGGGTTAACTTTAACAAGGAGAAAAACATGCAGGCGAGCGT<br>GTTTCAGGCCAGCGATCTGGGCGCGGGTTATATGGTGGCGAGCGCGGGCGAACATGCCAAA<br>AGCGGCGCAGCGAAATGCGGCGAAGGCAAATGCGGCGAAAAAAAAGCGGATTAAGCTTCG |
| FonA$_{PE}$A2G | MQASVFQASDLGAG<br>YMVASAGEHAKSGE<br>**G**KCGEGKCGEKKAD | GGCGTAATACGACTCACTATAGGGTTAACTTTAACAAGGAGAAAAACATGCAGGCGAGCGT<br>GTTTCAGGCCAGCGATCTGGGCGCGGGTTATATGGTGGCGAGCGCGGGCGAACATGCCAAA<br>AGCGGCGAAGGTAAATGCGGCGAAGGCAAATGCGGCGAAAAAAAAGCGGATTAAGCTTCG |
| FonA$_{PE}$K3A | MQASVFQASDLGAG<br>YMVASAGEHAKSGE<br>A**A**CGEGKCGEKKAD | GGCGTAATACGACTCACTATAGGGTTAACTTTAACAAGGAGAAAAACATGCAGGCGAGCGT<br>GTTTCAGGCCAGCGATCTGGGCGCGGGTTATATGGTGGCGAGCGCGGGCGAACATGCCAAA<br>AGCGGCGAAGCGGCATGCGGCGAAGGCAAATGCGGCGAAAAAAAAGCGGATTAAGCTTCG |
| FonA$_{PE}$C4A | MQASVFQASDLGAG<br>YMVASAGEHAKSGE<br>AK**A**GEGKCGEKKAD | GGCGTAATACGACTCACTATAGGGTTAACTTTAACAAGGAGAAAAACATGCAGGCGAGCGT<br>GTTTCAGGCCAGCGATCTGGGCGCGGGTTATATGGTGGCGAGCGCGGGCGAACATGCCAAA<br>AGCGGCGAAGCGAAAGCAGGCGAAGGCAAATGCGGCGAAAAAAAAGCGGATTAAGCTTCG |
| FonA$_{PE}$G5A | MQASVFQASDLGAG<br>YMVASAGEHAKSGE<br>AKC**A**EGKCGEKKAD | GGCGTAATACGACTCACTATAGGGTTAACTTTAACAAGGAGAAAAACATGCAGGCGAGCGT<br>GTTTCAGGCCAGCGATCTGGGCGCGGGTTATATGGTGGCGAGCGCGGGCGAACATGCCAAA<br>AGCGGCGAAGCGAAATGCGCAGAAGGCAAATGCGGCGAAAAAAAAGCGGATTAAGCTTCG |
| FonA$_{PE}$E6A | MQASVFQASDLGAG<br>YMVASAGEHAKSGE<br>AKCG**A**GKCGEKKAD | GGCGTAATACGACTCACTATAGGGTTAACTTTAACAAGGAGAAAAACATGCAGGCGAGCGT<br>GTTTCAGGCCAGCGATCTGGGCGCGGGTTATATGGTGGCGAGCGCGGGCGAACATGCCAAA<br>AGCGGCGAAGCGAAATGCGGCGCAGGCAAATGCGGCGAAAAAAAAGCGGATTAAGCTTCG |

**Table S10 (cont.). Constructs generated for *in vitro* transcription/translation.**

| Name | Peptide sequence | DNA template |
|---|---|---|
| FonA$_{PE}$G7A | MQASVFQASDLGAG<br>YMVASAGEHAKSGE<br>AKCGE**A**KCGEKKAD | GGCGTAATACGACTCACTATAGGGTTAACTTTAACAAGGAGAAAAACATGCAGGCGAGCGT<br>GTTTCAGGCCAGCGATCTGGGCGCGGGTTATATGGTGGCGAGCGCGGGCGAACATGCCAAA<br>AGCGGCGAAGCGAAATGCGGCGAAGCCAAATGCGGCGAAAAAAAAGCGGATTAAGCTTCG |
| FonA$_{PE}$K8A | MQASVFQASDLGAG<br>YMVASAGEHAKSGE<br>AKCGEG**A**CGEKKAD | GGCGTAATACGACTCACTATAGGGTTAACTTTAACAAGGAGAAAAACATGCAGGCGAGCGT<br>GTTTCAGGCCAGCGATCTGGGCGCGGGTTATATGGTGGCGAGCGCGGGCGAACATGCCAAA<br>AGCGGCGAAGCGAAATGCGGCGAAGGCGCATGCGGCGAAAAAAAAGCGGATTAAGCTTCG |
| FonA$_{PE}$C9A | MQASVFQASDLGAG<br>YMVASAGEHAKSGE<br>AKCGEGK**A**GEKKAD | GGCGTAATACGACTCACTATAGGGTTAACTTTAACAAGGAGAAAAACATGCAGGCGAGCGT<br>GTTTCAGGCCAGCGATCTGGGCGCGGGTTATATGGTGGCGAGCGCGGGCGAACATGCCAAA<br>AGCGGCGAAGCGAAATGCGGCGAAGGCAAAGCAGGCGAAAAAAAAGCGGATTAAGCTTCG |
| FonA$_{PE}$G10A | MQASVFQASDLGAG<br>YMVASAGEHAKSGE<br>AKCGEGKC**A**EKKAD | GGCGTAATACGACTCACTATAGGGTTAACTTTAACAAGGAGAAAAACATGCAGGCGAGCGT<br>GTTTCAGGCCAGCGATCTGGGCGCGGGTTATATGGTGGCGAGCGCGGGCGAACATGCCAAA<br>AGCGGCGAAGCGAAATGCGGCGAAGGCAAATGTGCAGAAAAAAAAGCGGATTAAGCTTCG |
| FonA$_{PE}$E11A | MQASVFQASDLGAG<br>YMVASAGEHAKSGE<br>AKCGEGKCG**A**KKAD | GGCGTAATACGACTCACTATAGGGTTAACTTTAACAAGGAGAAAAACATGCAGGCGAGCGT<br>GTTTCAGGCCAGCGATCTGGGCGCGGGTTATATGGTGGCGAGCGCGGGCGAACATGCCAAA<br>AGCGGCGAAGCGAAATGCGGCGAAGGCAAATGCGGCGCAAAAAAAGCGGATTAAGCTTCG |
| FonA$_{PE}$K12A | MQASVFQASDLGAG<br>YMVASAGEHAKSGE<br>AKCGEGKCGE**A**KAD | GGCGTAATACGACTCACTATAGGGTTAACTTTAACAAGGAGAAAAACATGCAGGCGAGCGT<br>GTTTCAGGCCAGCGATCTGGGCGCGGGTTATATGGTGGCGAGCGCGGGCGAACATGCCAAA<br>AGCGGCGAAGCGAAATGCGGCGAAGGCAAATGCGGCGAAGCAAAAGCGGATTAAGCTTCG |
| FonA$_{PE}$1-A | MQASVFQASDLGAG<br>YMVASAGEHAKSG**A**<br>AKCG**A**GKCGEKKAD | GGCGTAATACGACTCACTATAGGGTTAACTTTAACAAGGAGAAAAACATGCAGGCGAGCGT<br>GTTTCAGGCCAGCGATCTGGGCGCGGGTTATATGGTGGCGAGCGCGGGCGAACATGCCAAA<br>AGCGGCGCGGCGAAATGCGGCGCAGGCAAATGCGGCGAAAAAAAAGCGGATTAAGCTTCG |
| FonA$_{PE}$1-D | MQASVFQASDLGAG<br>YMVASAGEHAKSG**D**<br>AKCG**D**GKCGEKKAD | GGCGTAATACGACTCACTATAGGGTTAACTTTAACAAGGAGAAAAACATGCAGGCGAGCGT<br>GTTTCAGGCCAGCGATCTGGGCGCGGGTTATATGGTGGCGAGCGCGGGCGAACATGCCAAA<br>AGCGGCGATGCGAAATGCGGCGATGGCAAATGCGGCGAAAAAAAAGCGGATTAAGCTTCG |
| FonA$_{PE}$1-F | MQASVFQASDLGAG<br>YMVASAGEHAKSG**F**<br>AKCG**F**GKCGEKKAD | GGCGTAATACGACTCACTATAGGGTTAACTTTAACAAGGAGAAAAACATGCAGGCGAGCGT<br>GTTTCAGGCCAGCGATCTGGGCGCGGGTTATATGGTGGCGAGCGCGGGCGAACATGCCAAA<br>AGCGGCTTTGCGAAATGCGGCTTTGGCAAATGCGGCGAAAAAAAAGCGGATTAAGCTTCG |
| FonA$_{PE}$1-P | MQASVFQASDLGAG<br>YMVASAGEHAKSG**P**<br>AKCG**P**GKCGEKKAD | GGCGTAATACGACTCACTATAGGGTTAACTTTAACAAGGAGAAAAACATGCAGGCGAGCGT<br>GTTTCAGGCCAGCGATCTGGGCGCGGGTTATATGGTGGCGAGCGCGGGCGAACATGCCAAA<br>AGCGGCCCTGCGAAATGCGGCCCTGGCAAATGCGGCGAAAAAAAAGCGGATTAAGCTTCG |
| FonA$_{PE}$1- | MQASVFQASDLGAG<br>YMVASAGEHAKSG**Q**<br>AKCG**Q**GKCGEKKAD | GGCGTAATACGACTCACTATAGGGTTAACTTTAACAAGGAGAAAAACATGCAGGCGAGCGT<br>GTTTCAGGCCAGCGATCTGGGCGCGGGTTATATGGTGGCGAGCGCGGGCGAACATGCCAAA<br>AGCGGCCAGGCGAAATGCGGCCAGGGCAAATGCGGCGAAAAAAAAGCGGATTAAGCTTCG |
| FonA$_{PE}$1-R | MQASVFQASDLGAG<br>YMVASAGEHAKSG**R**<br>AKCG**R**GKCGEKKAD | GGCGTAATACGACTCACTATAGGGTTAACTTTAACAAGGAGAAAAACATGCAGGCGAGCGT<br>GTTTCAGGCCAGCGATCTGGGCGCGGGTTATATGGTGGCGAGCGCGGGCGAACATGCCAAA<br>AGCGGCCGTGCGAAATGCGGCCGTGGCAAATGCGGCGAAAAAAAAGCGGATTAAGCTTCG |
| FonA$_{PE}$1-T | MQASVFQASDLGAG<br>YMVASAGEHAKSG**T**<br>AKCG**T**GKCGEKKAD | GGCGTAATACGACTCACTATAGGGTTAACTTTAACAAGGAGAAAAACATGCAGGCGAGCGT<br>GTTTCAGGCCAGCGATCTGGGCGCGGGTTATATGGTGGCGAGCGCGGGCGAACATGCCAAA<br>AGCGGCACCGCGAAATGCGGCACTGGCAAATGCGGCGAAAAAAAAGCGGATTAAGCTTCG |
| FonA$_{PE}$2-E | MQASVFQASDLGAG<br>YMVASAGEHAKSGE<br>**E**KCGE**E**KCGEKKAD | GGCGTAATACGACTCACTATAGGGTTAACTTTAACAAGGAGAAAAACATGCAGGCGAGCGT<br>GTTTCAGGCCAGCGATCTGGGCGCGGGTTATATGGTGGCGAGCGCGGGCGAACATGCCAAA<br>AGCGGCGAAGAAAAATGCGGCGAAGAAAAATGCGGCGAAAAAAAAGCGGATTAAGCTTCG |
| FonA$_{PE}$2-F | MQASVFQASDLGAG<br>YMVASAGEHAKSGE<br>**F**KCGE**F**KCGEKKAD | GGCGTAATACGACTCACTATAGGGTTAACTTTAACAAGGAGAAAAACATGCAGGCGAGCGT<br>GTTTCAGGCCAGCGATCTGGGCGCGGGTTATATGGTGGCGAGCGCGGGCGAACATGCCAAA<br>AGCGGCGAATTTAAATGCGGCGAATTTAAATGCGGCGAAAAAAAAGCGGATTAAGCTTCG |
| FonA$_{PE}$2-P | MQASVFQASDLGAG<br>YMVASAGEHAKSGE<br>**P**KCGE**P**KCGEKKAD | GGCGTAATACGACTCACTATAGGGTTAACTTTAACAAGGAGAAAAACATGCAGGCGAGCGT<br>GTTTCAGGCCAGCGATCTGGGCGCGGGTTATATGGTGGCGAGCGCGGGCGAACATGCCAAA<br>AGCGGCGAACCGAAATGCGGCGAACCTAAATGCGGCGAAAAAAAAGCGGATTAAGCTTCG |
| FonA$_{PE}$2-R | MQASVFQASDLGAG<br>YMVASAGEHAKSGE<br>**R**KCGE**R**KCGEKKAD | GGCGTAATACGACTCACTATAGGGTTAACTTTAACAAGGAGAAAAACATGCAGGCGAGCGT<br>GTTTCAGGCCAGCGATCTGGGCGCGGGTTATATGGTGGCGAGCGCGGGCGAACATGCCAAA<br>AGCGGCGAACGTAAATGCGGCGAACGTAAATGCGGCGAAAAAAAAGCGGATTAAGCTTCG |
| FonA$_{PE}$2-T | MQASVFQASDLGAG<br>YMVASAGEHAKSGE<br>**T**KCGE**T**KCGEKKAD | GGCGTAATACGACTCACTATAGGGTTAACTTTAACAAGGAGAAAAACATGCAGGCGAGCGT<br>GTTTCAGGCCAGCGATCTGGGCGCGGGTTATATGGTGGCGAGCGCGGGCGAACATGCCAAA<br>AGCGGCGAAACGAAATGCGGCGAAACTAAATGCGGCGAAAAAAAAGCGGATTAAGCTTCG |
| FonA$_{PE}$3-A | MQASVFQASDLGAG<br>YMVASAGEHAKSGE<br>A**A**CGEG**A**CGEKKAD | GGCGTAATACGACTCACTATAGGGTTAACTTTAACAAGGAGAAAAACATGCAGGCGAGCGT<br>GTTTCAGGCCAGCGATCTGGGCGCGGGTTATATGGTGGCGAGCGCGGGCGAACATGCCAAA<br>AGCGGCGAAGCGGCGTGCGGCGAAGGCGCATGCGGCGAAAAAAAAGCGGATTAAGCTTCG |
| FonA$_{PE}$3-E | MQASVFQASDLGAG<br>YMVASAGEHAKSGE<br>A**E**CGEG**E**CGEKKAD | GGCGTAATACGACTCACTATAGGGTTAACTTTAACAAGGAGAAAAACATGCAGGCGAGCGT<br>GTTTCAGGCCAGCGATCTGGGCGCGGGTTATATGGTGGCGAGCGCGGGCGAACATGCCAAA<br>AGCGGCGAAGCGGAATGCGGCGAAGGCGAATGCGGCGAAAAAAAAGCGGATTAAGCTTCG |
| FonA$_{PE}$3-F | MQASVFQASDLGAG<br>YMVASAGEHAKSGE<br>A**F**CGEG**F**CGEKKAD | GGCGTAATACGACTCACTATAGGGTTAACTTTAACAAGGAGAAAAACATGCAGGCGAGCGT<br>GTTTCAGGCCAGCGATCTGGGCGCGGGTTATATGGTGGCGAGCGCGGGCGAACATGCCAAA<br>AGCGGCGAAGCGTTTTGCGGCGAAGGCTTTTGCGGCGAAAAAAAAGCGGATTAAGCTTCG |

**Table S10 (cont.). Constructs generated for *in vitro* transcription/translation.**

| Name | Peptide sequence | DNA template |
|---|---|---|
| FonA$_{PE}$3-P | MQASVFQASDLGAG<br>YMVASAGEHAKSGE<br>A**P**CGEG**P**CGEKKAD | GGCGTAATACGACTCACTATAGGGTTAACTTTAACAAGGAGAAAAACATGCAGGCGAGCGTG<br>TTTCAGGCCAGCGATCTGGGCGCGGGTTATATGGTGGCGAGCGCGGGCGAACATGCCAAAAG<br>CGGCGAAGCGCCGTGCGGCGAAGGCCCTTGCGGCGAAAAAAAAGCGGATTAAGCTTCG |
| FonA$_{PE}$3-R | MQASVFQASDLGAG<br>YMVASAGEHAKSGE<br>A**R**CGEG**R**CGEKKAD | GGCGTAATACGACTCACTATAGGGTTAACTTTAACAAGGAGAAAAACATGCAGGCGAGCGTG<br>TTTCAGGCCAGCGATCTGGGCGCGGGTTATATGGTGGCGAGCGCGGGCGAACATGCCAAAAG<br>CGGCGAAGCGCGCTGCGGCGAAGGCCGTTGCGGCGAAAAAAAAGCGGATTAAGCTTCG |
| FonA$_{PE}$3-T | MQASVFQASDLGAG<br>YMVASAGEHAKSGE<br>A**T**CGEG**T**CGEKKAD | GGCGTAATACGACTCACTATAGGGTTAACTTTAACAAGGAGAAAAACATGCAGGCGAGCGTG<br>TTTCAGGCCAGCGATCTGGGCGCGGGTTATATGGTGGCGAGCGCGGGCGAACATGCCAAAAG<br>CGGCGAAGCGACCTGCGGCGAAGGCACTTGCGGCGAAAAAAAAGCGGATTAAGCTTCG |
| FonA$_{PE}$4-A | MQASVFQASDLGAG<br>YMVASAGEHAKSGE<br>AK**A**GEGK**A**GEKKAD | GGCGTAATACGACTCACTATAGGGTTAACTTTAACAAGGAGAAAAACATGCAGGCGAGCGTG<br>TTTCAGGCCAGCGATCTGGGCGCGGGTTATATGGTGGCGAGCGCGGGCGAACATGCCAAAAG<br>CGGCGAAGCGAAAGCGGGCGAAGGCAAAGCAGGCGAAAAAAAAGCGGATTAAGCTTCG |
| FonA$_{PE}$4-D | MQASVFQASDLGAG<br>YMVASAGEHAKSGE<br>AK**D**GEGK**D**GEKKAD | GGCGTAATACGACTCACTATAGGGTTAACTTTAACAAGGAGAAAAACATGCAGGCGAGCGTG<br>TTTCAGGCCAGCGATCTGGGCGCGGGTTATATGGTGGCGAGCGCGGGCGAACATGCCAAAAG<br>CGGCGAAGCGAAAGATGGCGAAGGCAAAGATGGCGAAAAAAAAGCGGATTAAGCTTCG |
| FonA$_{PE}$4-M | MQASVFQASDLGAG<br>YMVASAGEHAKSGE<br>AK**M**GEGK**M**GEKKAD | GGCGTAATACGACTCACTATAGGGTTAACTTTAACAAGGAGAAAAACATGCAGGCGAGCGTG<br>TTTCAGGCCAGCGATCTGGGCGCGGGTTATATGGTGGCGAGCGCGGGCGAACATGCCAAAAG<br>CGGCGAAGCGAAAATGGGCGAAGGCAAAATGGGCGAAAAAAAAGCGGATTAAGCTTCG |
| FonA$_{PE}$5-A | MQASVFQASDLGAG<br>YMVASAGEHAKSGE<br>AKC**A**EGKC**A**EKKAD | GGCGTAATACGACTCACTATAGGGTTAACTTTAACAAGGAGAAAAACATGCAGGCGAGCGTG<br>TTTCAGGCCAGCGATCTGGGCGCGGGTTATATGGTGGCGAGCGCGGGCGAACATGCCAAAAG<br>CGGCGAAGCGAAATGCGCAGAAGGCAAATGCGCAGAAAAAAAAGCGGATTAAGCTTCG |
| FonA$_{PE}$5-E | MQASVFQASDLGAG<br>YMVASAGEHAKSGE<br>AKC**E**EGKC**E**EKKAD | GGCGTAATACGACTCACTATAGGGTTAACTTTAACAAGGAGAAAAACATGCAGGCGAGCGTG<br>TTTCAGGCCAGCGATCTGGGCGCGGGTTATATGGTGGCGAGCGCGGGCGAACATGCCAAAAG<br>CGGCGAAGCGAAATGCGAAGAAGGCAAATGCGAAGAAAAAAAAGCGGATTAAGCTTCG |
| FonA$_{PE}$5-F | MQASVFQASDLGAG<br>YMVASAGEHAKSGE<br>AKC**F**EGKC**F**EKKAD | GGCGTAATACGACTCACTATAGGGTTAACTTTAACAAGGAGAAAAACATGCAGGCGAGCGTG<br>TTTCAGGCCAGCGATCTGGGCGCGGGTTATATGGTGGCGAGCGCGGGCGAACATGCCAAAAG<br>CGGCGAAGCGAAATGCTTTGAAGGCAAATGCTTTGAAAAAAAAGCGGATTAAGCTTCG |
| FonA$_{PE}$5-P | MQASVFQASDLGAG<br>YMVASAGEHAKSGE<br>AKC**P**EGKC**P**EKKAD | GGCGTAATACGACTCACTATAGGGTTAACTTTAACAAGGAGAAAAACATGCAGGCGAGCGTG<br>TTTCAGGCCAGCGATCTGGGCGCGGGTTATATGGTGGCGAGCGCGGGCGAACATGCCAAAAG<br>CGGCGAAGCGAAATGCCCTGAAGGCAAATGCCCTGAAAAAAAAGCGGATTAAGCTTCG |
| FonA$_{PE}$5-R | MQASVFQASDLGAG<br>YMVASAGEHAKSGE<br>AKC**R**EGKC**R**EKKAD | GGCGTAATACGACTCACTATAGGGTTAACTTTAACAAGGAGAAAAACATGCAGGCGAGCGTG<br>TTTCAGGCCAGCGATCTGGGCGCGGGTTATATGGTGGCGAGCGCGGGCGAACATGCCAAAAG<br>CGGCGAAGCGAAATGCCGTGAAGGCAAATGCCGTGAAAAAAAAGCGGATTAAGCTTCG |
| FonA$_{PE}$5-T | MQASVFQASDLGAG<br>YMVASAGEHAKSGE<br>AKC**T**EGKC**T**EKKAD | GGCGTAATACGACTCACTATAGGGTTAACTTTAACAAGGAGAAAAACATGCAGGCGAGCGTG<br>TTTCAGGCCAGCGATCTGGGCGCGGGTTATATGGTGGCGAGCGCGGGCGAACATGCCAAAAG<br>CGGCGAAGCGAAATGCACTGAAGGCAAATGCACTGAAAAAAAAGCGGATTAAGCTTCG |
| FonA$_{PE}$AKCGA | MQASVFQASDLGAG<br>YMVASAGEHAKSG**A**<br>**KCGAAKCGA**EKKAD | GGCGTAATACGACTCACTATAGGGTTAACTTTAACAAGGAGAAAAACATGCAGGCGAGCGTG<br>TTTCAGGCCAGCGATCTGGGCGCGGGTTATATGGTGGCGAGCGCGGGCGAACATGCCAAAAG<br>CGGCGCGAAATGCGGTGCGGCGAAATGTGGTGCGGAAAAAAAAGCGGATTAAGCTTCG |
| FonA$_{PE}$DGKCG | MQASVFQASDLGAG<br>YMVASAGEHAKSG**D**<br>**GKCGDGKCG**EKKAD | GGCGTAATACGACTCACTATAGGGTTAACTTTAACAAGGAGAAAAACATGCAGGCGAGCGTG<br>TTTCAGGCCAGCGATCTGGGCGCGGGTTATATGGTGGCGAGCGCGGGCGAACATGCCAAAAG<br>CGGCGATGGCAAATGCGGCGATGGCAAATGCGGCGAAAAAAAAGCGGATTAAGCTTCG |
| FonA$_{PE}$GKCGT | MQASVFQASDLGAG<br>YMVASAGEHAKSG**G**<br>**KCGTGKCGT**EKKAD | GGCGTAATACGACTCACTATAGGGTTAACTTTAACAAGGAGAAAAACATGCAGGCGAGCGTG<br>TTTCAGGCCAGCGATCTGGGCGCGGGTTATATGGTGGCGAGCGCGGGCGAACATGCCAAAAG<br>CGGCGGCAAATGCGGCACCGGCAAATGTGGCACCGAAAAAAAAGCGGATTAAGCTTCG |
| FonA$_{PE}$HNDCK | MQASVFQASDLGAG<br>YMVASAGEHAKSG**H**<br>**NDCKHNDCK**EKKAD | GGCGTAATACGACTCACTATAGGGTTAACTTTAACAAGGAGAAAAACATGCAGGCGAGCGTG<br>TTTCAGGCCAGCGATCTGGGCGCGGGTTATATGGTGGCGAGCGCGGGCGAACATGCCAAAAG<br>CGGCCACAACGATTGCAAACACAACGACTGCAAAGAAAAAAAAGCGGATTAAGCTTCG |
| FonA$_{PE}$LVHCY | MQASVFQASDLGAG<br>YMVASAGEHAKSG**L**<br>**VHCYLVHCY**EKKAD | GGCGTAATACGACTCACTATAGGGTTAACTTTAACAAGGAGAAAAACATGCAGGCGAGCGTG<br>TTTCAGGCCAGCGATCTGGGCGCGGGTTATATGGTGGCGAGCGCGGGCGAACATGCCAAAAG<br>CGGCCTGGTGCATTGCTATCTGGTGCATTGCTATGAAAAAAAAGCGGATTAAGCTTCG |
| FonA$_{PE}$NACKG | MQASVFQASDLGAG<br>YMVASAGEHAKSG**N**<br>**ACKGNACKG**EKKAD | GGCGTAATACGACTCACTATAGGGTTAACTTTAACAAGGAGAAAAACATGCAGGCGAGCGTG<br>TTTCAGGCCAGCGATCTGGGCGCGGGTTATATGGTGGCGAGCGCGGGCGAACATGCCAAAAG<br>CGGCAACGCGTGCAAAGGCAACGCGTGCAAAGGCGAAAAAAAAGCGGATTAAGCTTCG |
| FonA$_{PE}$NVCGG | MQASVFQASDLGAG<br>YMVASAGEHAKSG**N**<br>**VCGGNVCGG**EKKAD | GGCGTAATACGACTCACTATAGGGTTAACTTTAACAAGGAGAAAAACATGCAGGCGAGCGTG<br>TTTCAGGCCAGCGATCTGGGCGCGGGTTATATGGTGGCGAGCGCGGGCGAACATGCCAAAAG<br>CGGCAATGTGTGTGGCGGCAATGTGTGCGGCGGCGAAAAAAAAGCGGATTAAGCTTCG |
| FonA$_{PE}$QCEKA | MQASVFQASDLGAG<br>YMVASAGEHAKSG**Q**<br>**CEKAQCEKA**EKKAD | GGCGTAATACGACTCACTATAGGGTTAACTTTAACAAGGAGAAAAACATGCAGGCGAGCGTG<br>TTTCAGGCCAGCGATCTGGGCGCGGGTTATATGGTGGCGAGCGCGGGCGAACATGCCAAAAG<br>CGGCCAGTGCGAAAAAGCGCAGTGCGAAAAAGCGGAAAAAAAAGCGGATTAAGCTTCG |
| FonA$_{PE}$1Cys | MQASVFQASDLGAG<br>YMVASAGEHAKSG**E**<br>**AKCG**EGK | GGCGTAATACGACTCACTATAGGGTTAACTTTAACAAGGAGAAAAACATGCAGGCGAGCGTG<br>TTTCAGGCCAGCGATCTGGGCGCGGGTTATATGGTGGCGAGCGCGGGCGAACATGCCAAAAG<br>CGGCGAAGCGAAATGCGGCGAAGGCAAATAAGCTTCG |

Logo 1

Logo 2

Logo 3

Logo 4

Logo 5

Logo 6

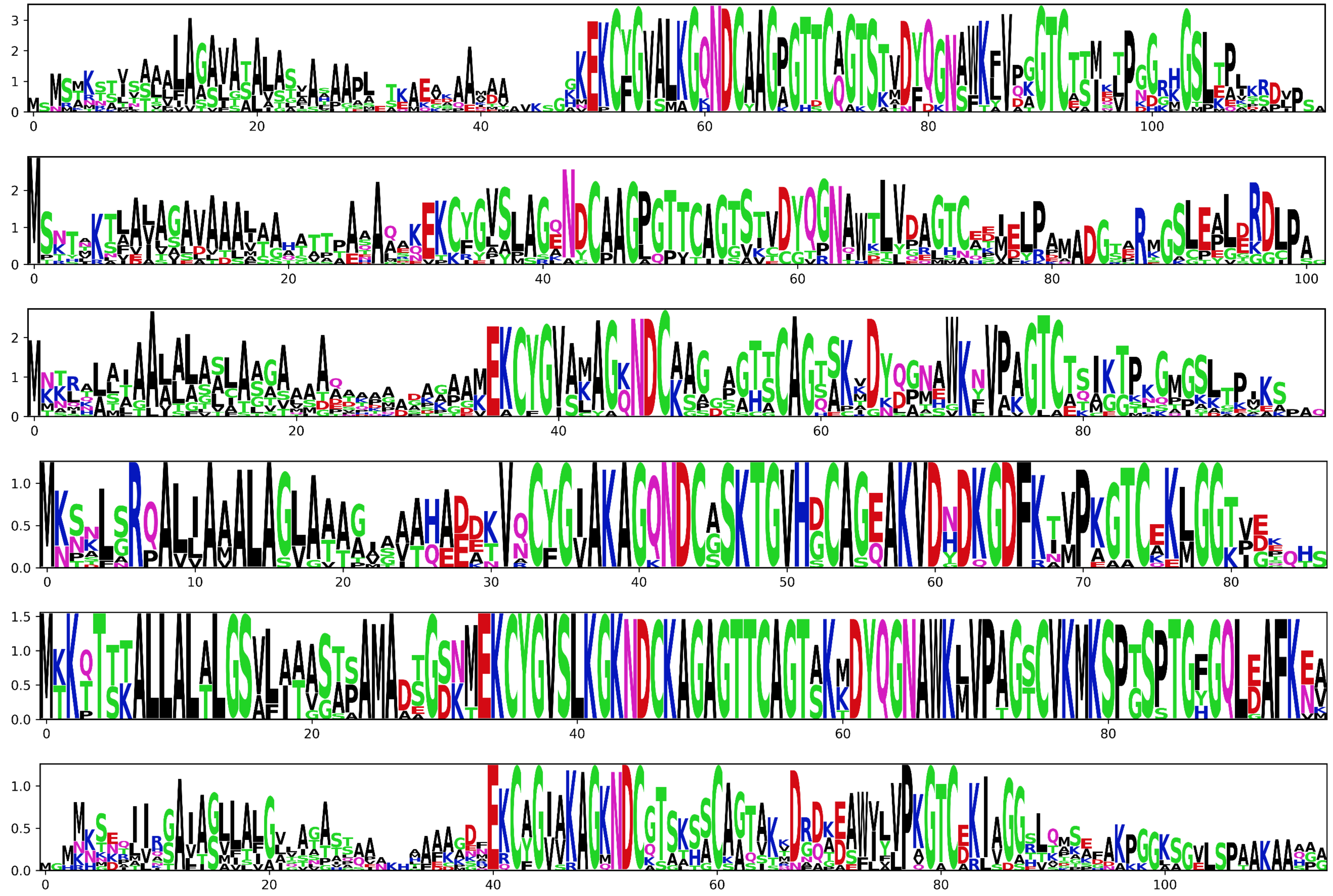

**Figure S1. Sequence logos for putative MNIO-PME1 families identified from RRE-Finder data.** Logos were generated using a customized version of the script developed for SPECO. The y-axis represents pseudobits, and the x-axis represents position within alignment. Logos represent differing numbers of input sequences as follows: 1, $n = 540$; 2, $n = 367$; 3, $n = 462$; 4, $n = 35$; 5, $n = 41$; 6, $n = 36$; 7, $n = 63$; 8, $n = 85$; 9, $n = 300$; 10, $n = 391$; 11, $n = 151$; 12, $n = 77$; 13, $n = 26$; 14, $n = 191$; 15, $n = 99$; 16, $n = 118$; 17, $n = 479$; 18, $n = 37$.

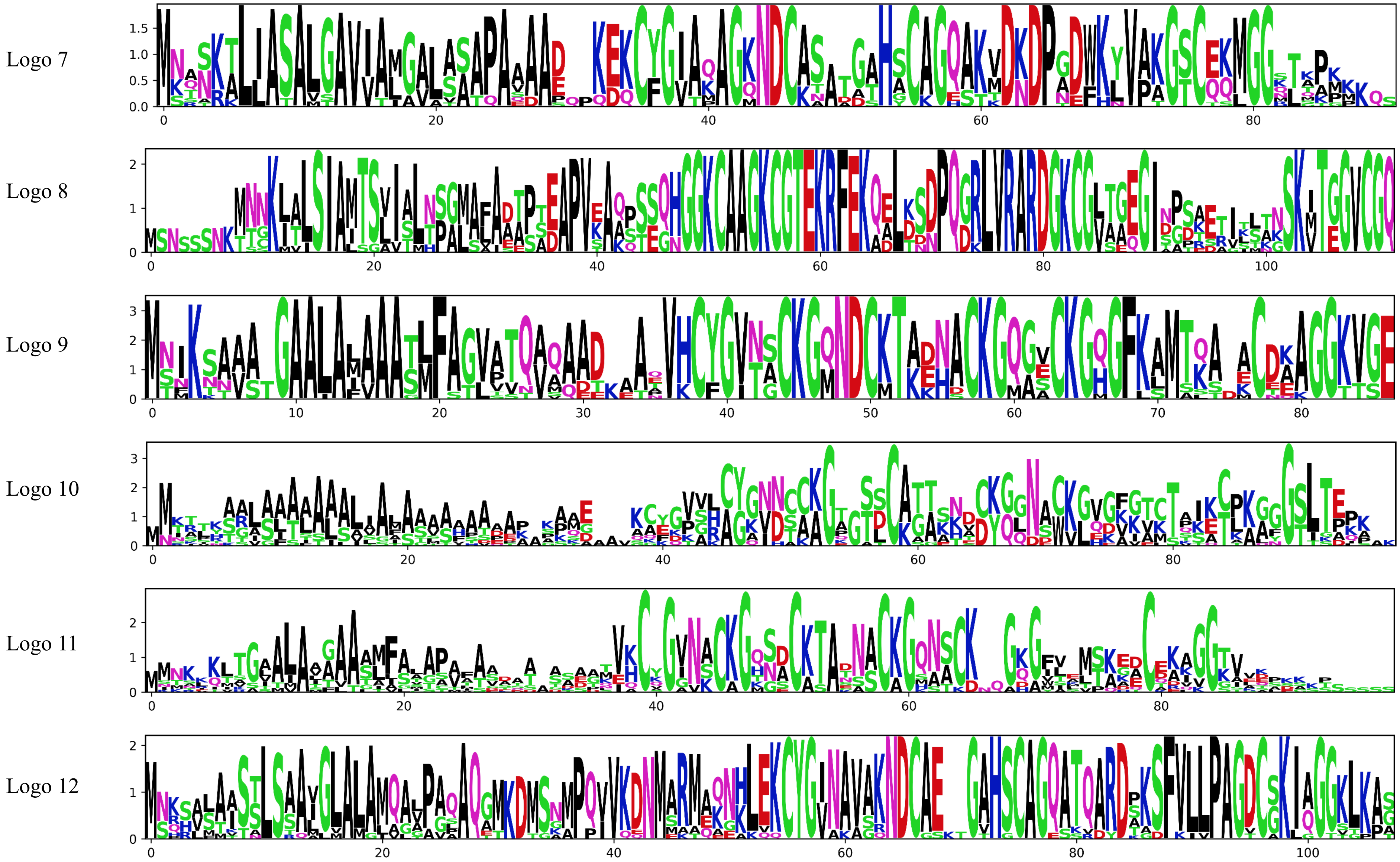


**Figure S1 (cont.). Sequence logos for putative MNIO-PME1 families identified from RRE-Finder data.**

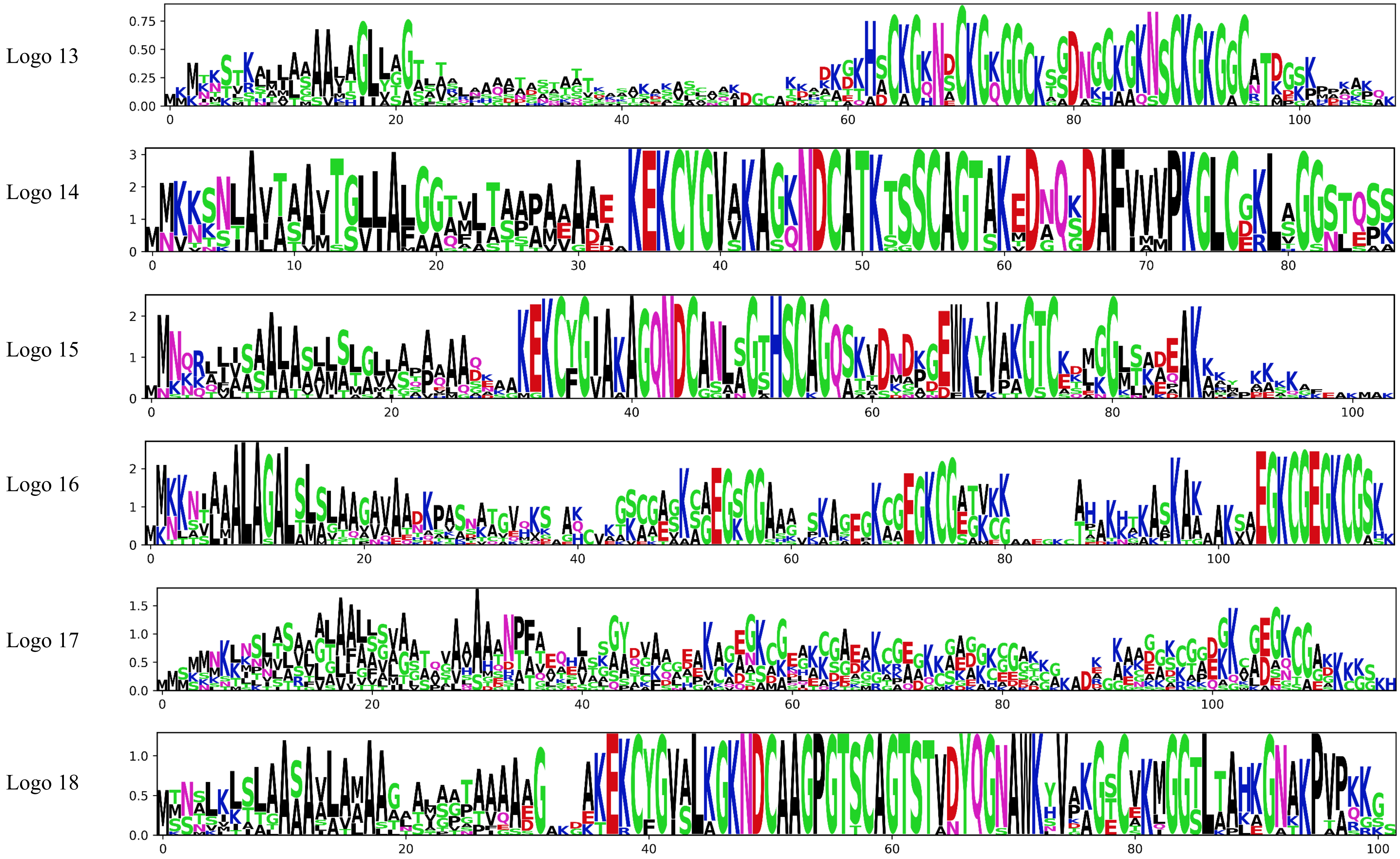


**Figure S1 (cont.). Sequence logos for putative MNIO-PME1 families identified from RRE-Finder data.**

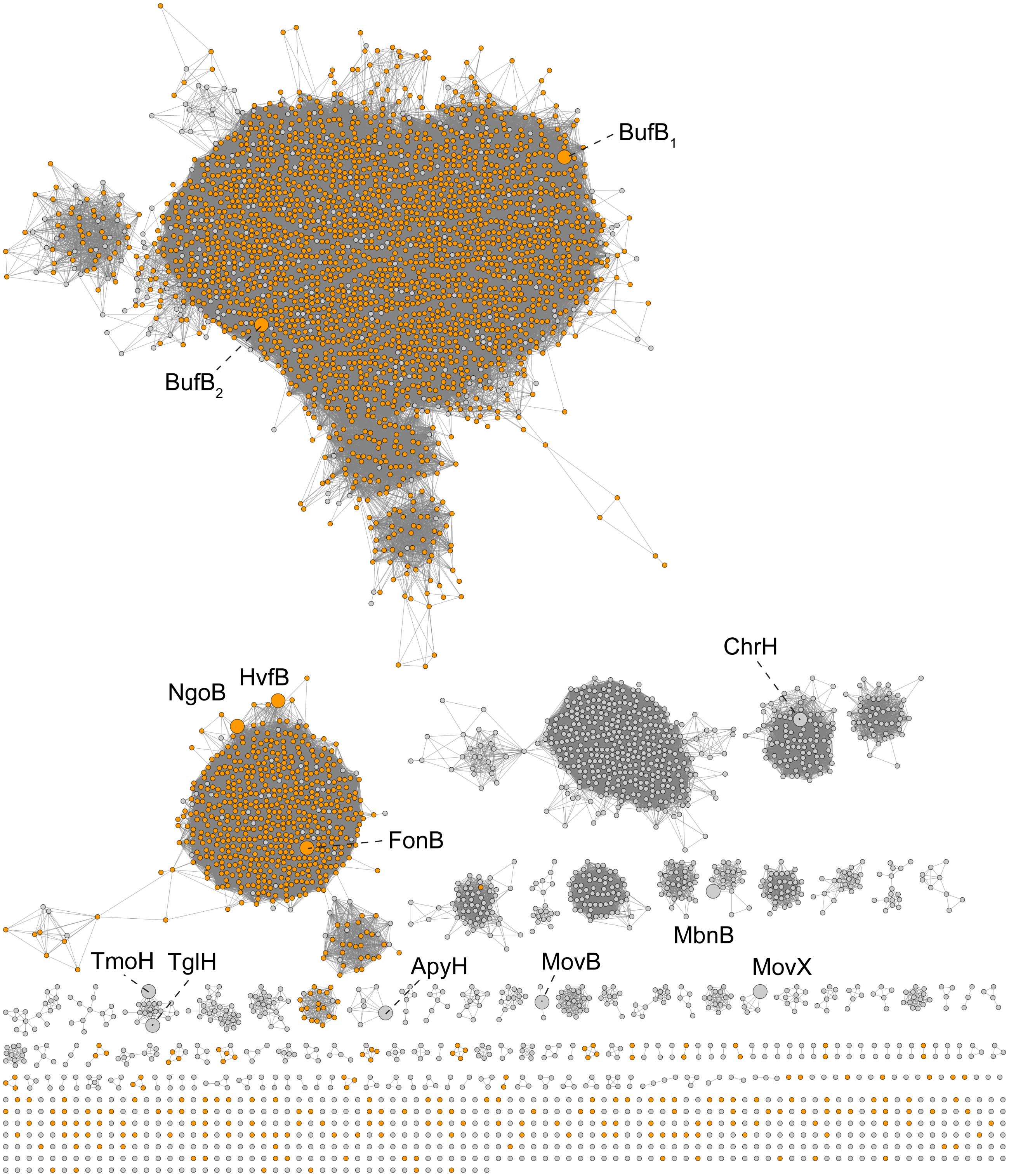


**Figure S2. Sequence similarity network of MNIO family.** Network was generated at alignment score of 75. Nodes were conflated at >60% identity. Color annotations: Orange, Local co-occurrence with DUF2063 protein (PME1).

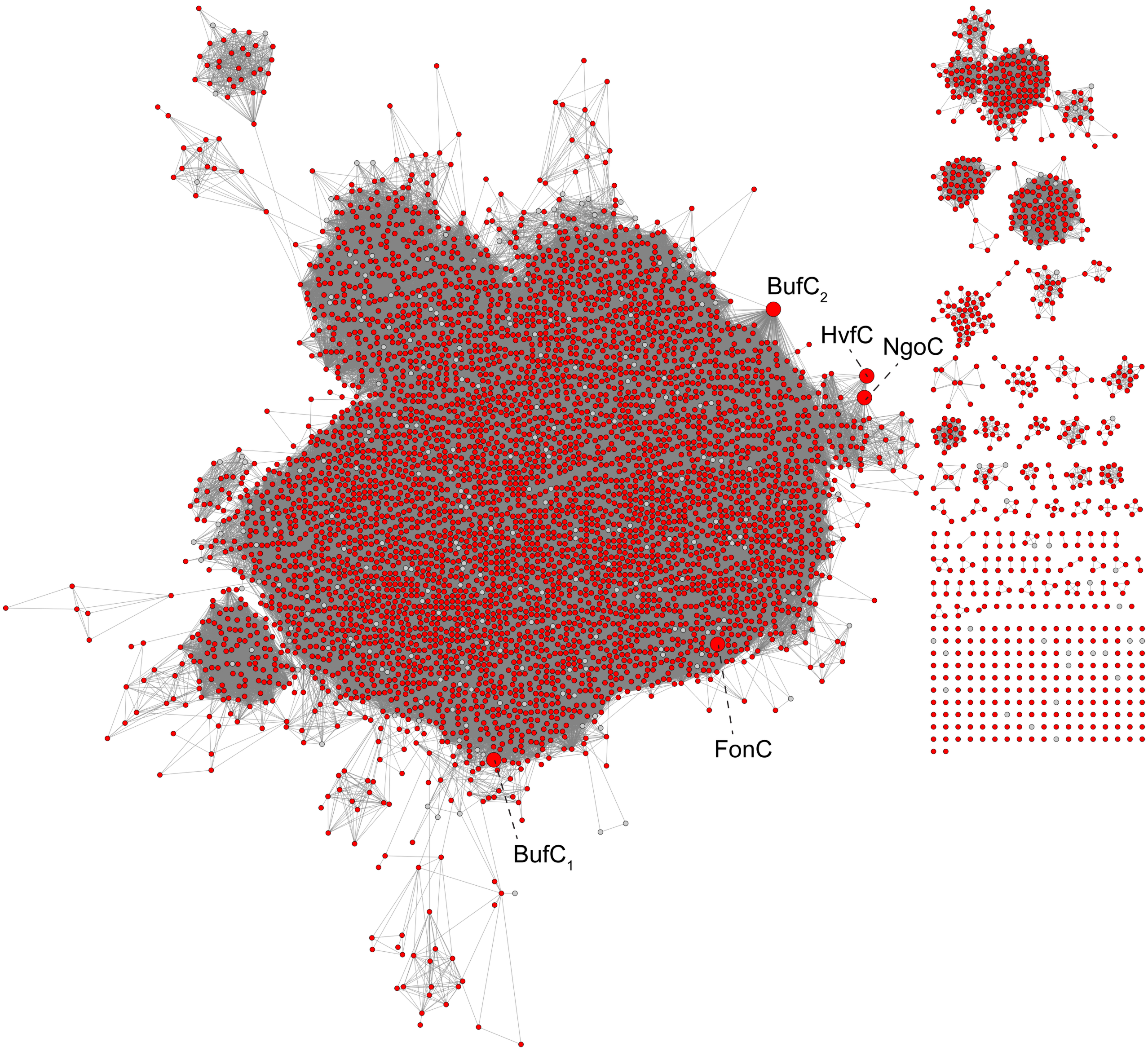


**Figure S3. Sequence similarity network for PME1 family.** Network was generated at alignment score of 34. Nodes were conflated at >70% identity. Color annotations: Red, Local co-occurrence with MNIO.

**a**

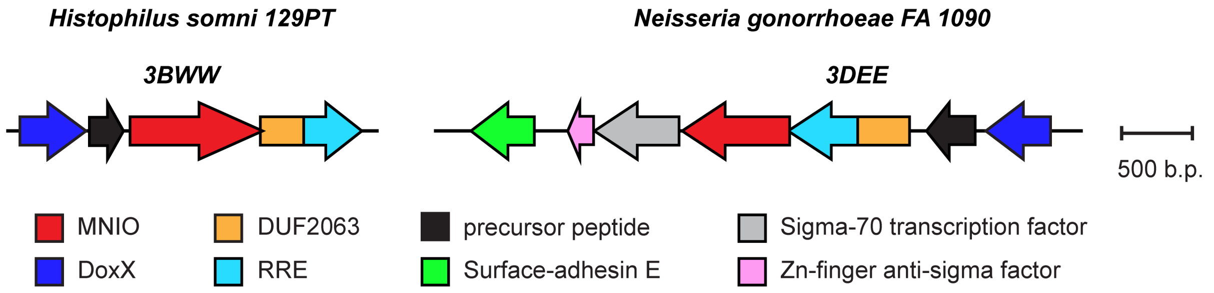


**b**

***HsoA sequence***

`MKKTTLVALAGALTMAVSAHADMKNTTNDCMEMNGKCVS`
`TKSA`**`EGKCGEGKCG`**`ATTPKKNMPKSA`**`EGKCGEGKCG`**`ASK`

***NgoA sequence***

`MNKNIAAALAGALSLSLAAGAVAAHKPASNATGVQKSA`**`QGSCG`**`ASKSA`**`EGSCG`**`ASK`
`SA`**`EGSCG`**`AAASKAG`**`EGKCGEGKCG`**`ATVKKAHKHTKASKAKAKSA`**`EGKCGEGKCG`**`SK`

**Figure S4. MNIO BGCs from *H. somni* and *N. gonorrhoeae*. a**, BGC diagrams for *hso* from *Histophilus somni* and *ngo* from *Neisseria gonorrhoeae*. PDB codes 3BWW[8] and 3DEE[9,10] are structurally characterized proteins encoded within these BGCs. **b**, HsoA and NgoA precursor peptide sequences.

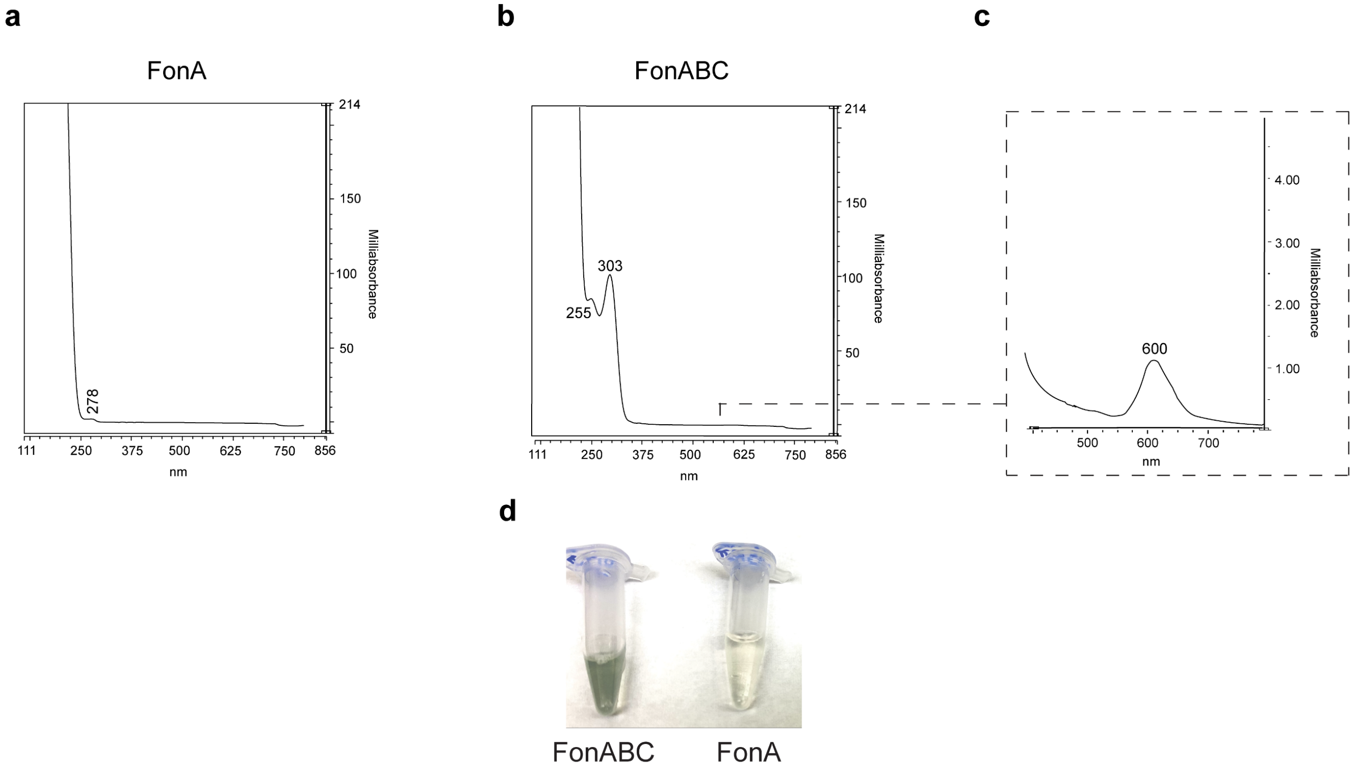


**Figure S5. UV-Vis spectral comparison of unmodified FonA to fontiphorin.** **a**, UV-Vis spectrum of purified FonA (unmodified). **b**, UV-Vis spectrum of purified FonABC (modified). **c**, Zoom-in of panel **b** showing a minor local absorbance maximum at 600 nm. **d**, Visual comparison of the color difference between FonA and FonABC upon concentration of the purified peptides to equal concentrations.

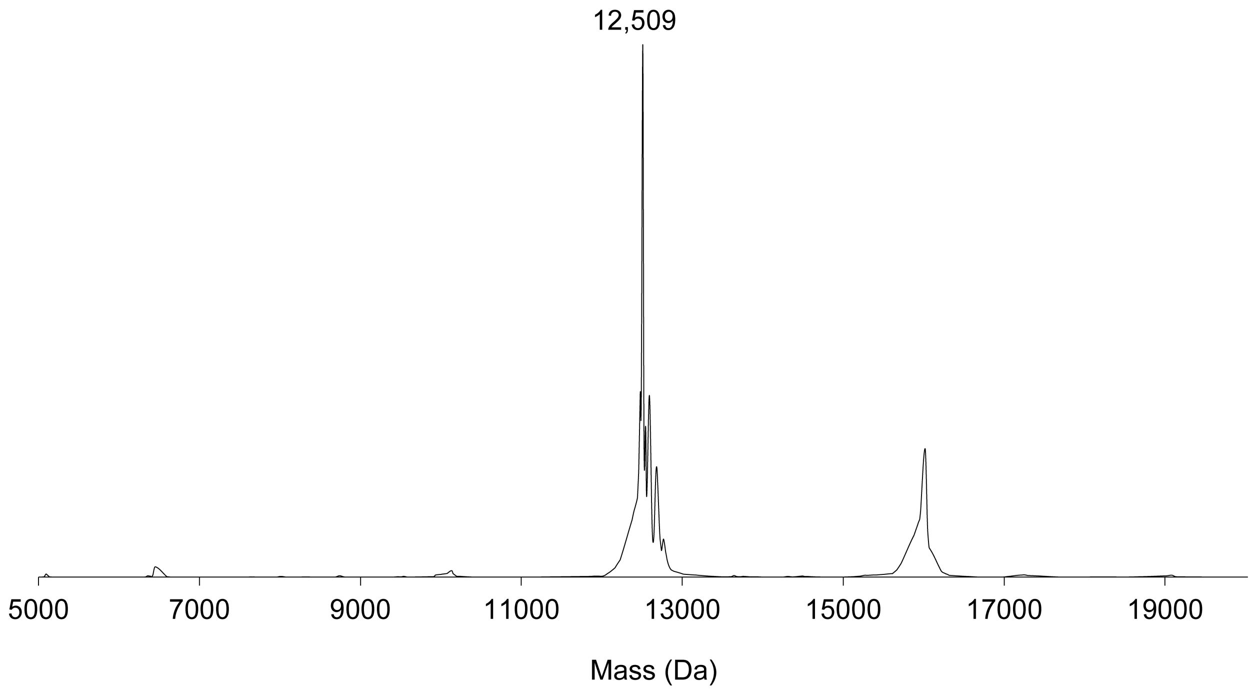


**Figure S6. Deconvoluted mass spectrum of full-length fontiphorin.** Expected average mass for unmodified FonA, 12,539 Da; Observed mass, 12,509 Da.

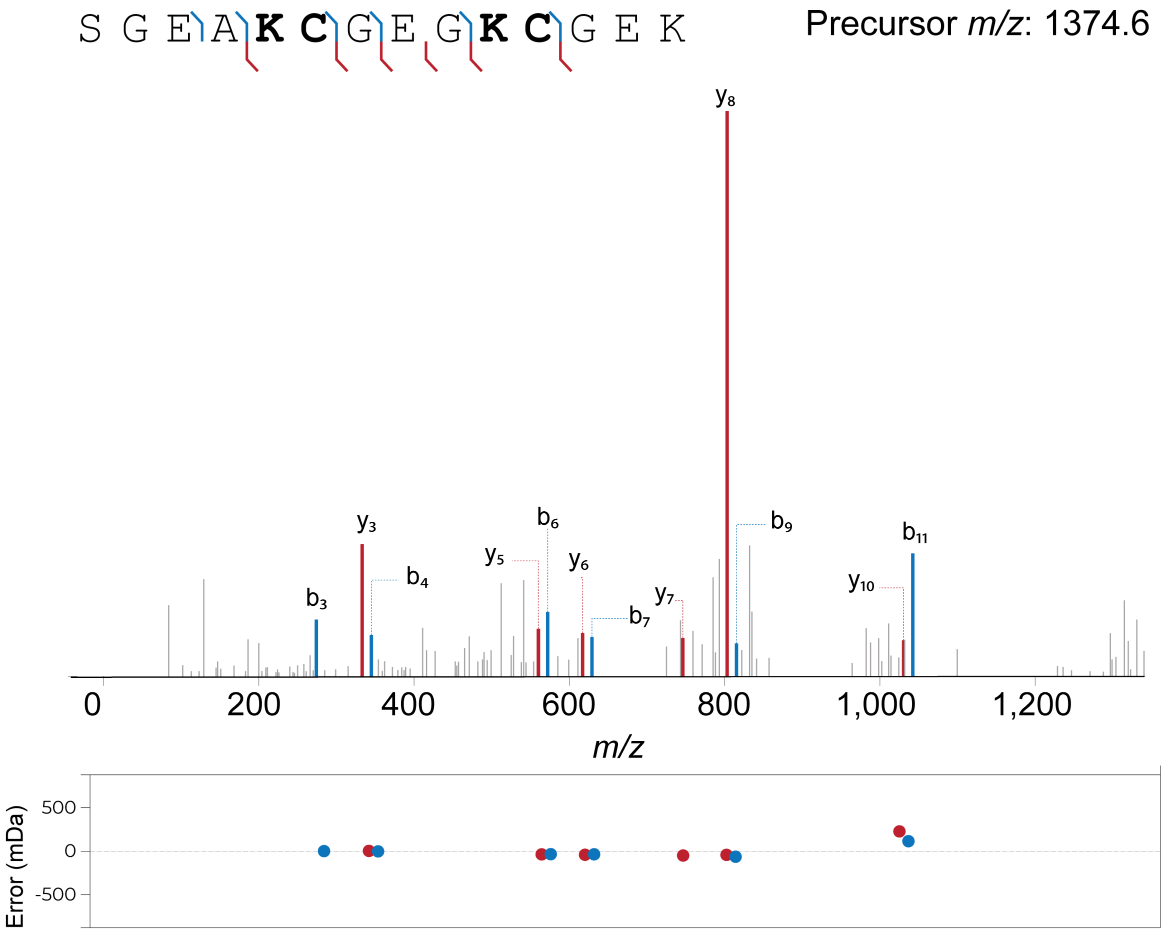


**Figure S7. MALDI-LIFT-MS for FonA$_{67\text{-}80}$.** Annotated b-ion and y-ion fragments are shown on the peptide sequence and tandem mass spectrum of modified FonA$_{67\text{-}80}$.

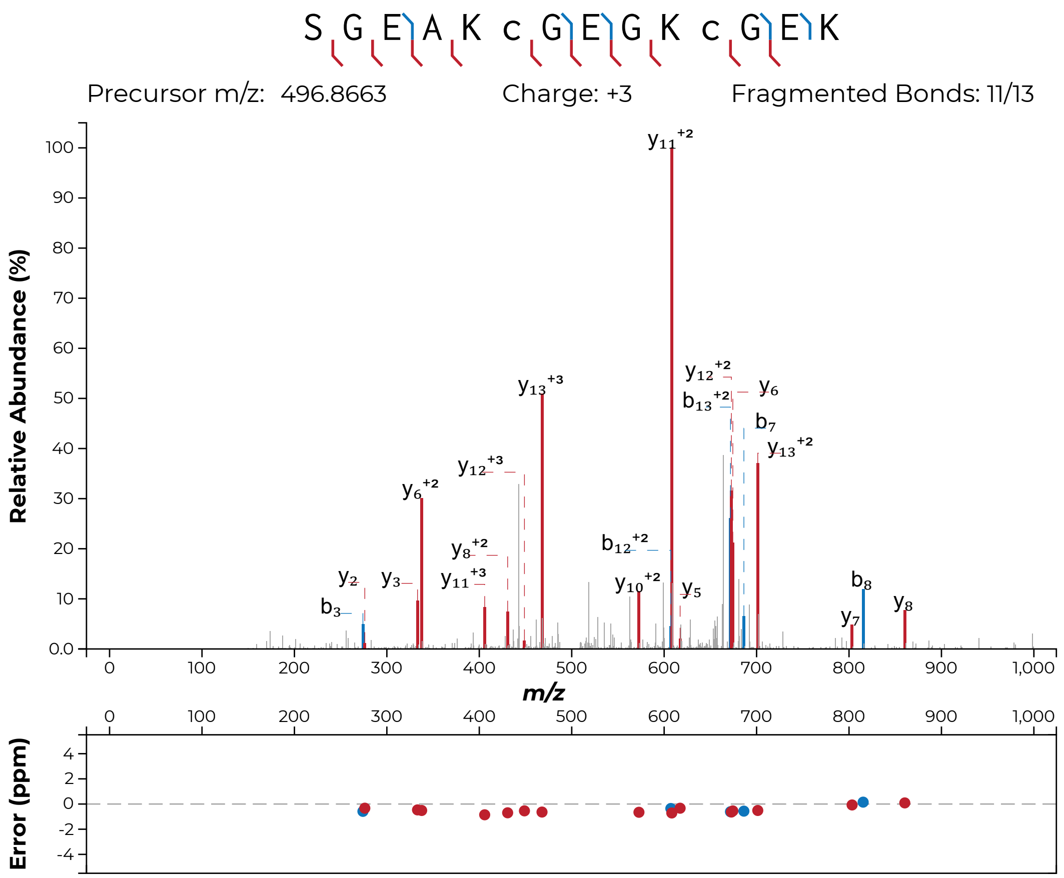


**Figure S8. HRMS/MS analysis of alkylated FonA$_{67\text{-}80}$.** Annotated b-ion and y-ion fragments are shown on the peptide sequence and tandem mass spectrum of modified and alkylated FonA$_{67\text{-}80}$.

**a**

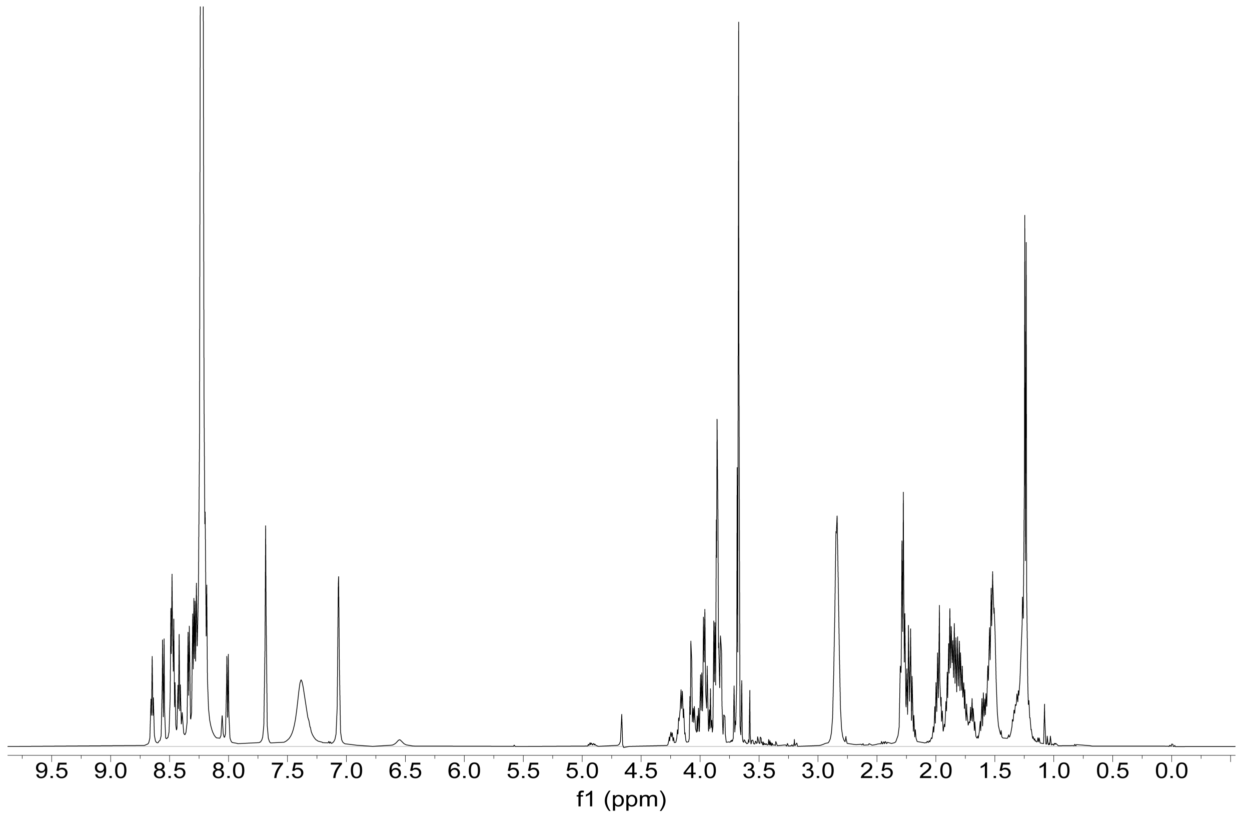


**Figure S9. NMR spectra for FonA$_{67\text{-}80}$.** **a**, $^{1}H$ NMR spectrum acquired in 90%:10% (*v/v*) $H_2O$:$D_2O$, both containing 0.1% formic acid-$d_2$. **b**, $^{1}H$ NMR spectrum acquired in 99.9% $D_2O$ + 0.1% formic acid-$d_2$. **c**, $^{13}C$ NMR spectrum acquired in 90%:10% (*v/v*) $H_2O$:$D_2O$, both containing 0.1% formic acid-$d_2$. **d**, $^{1}H$-$^{1}H$ TOCSY spectrum acquired in 90%:10% (*v/v*) $H_2O$:$D_2O$, both containing 0.1% formic acid-$d_2$. **e**, $^{1}H$-$^{13}C$ HSQC spectrum acquired in 99.9% $D_2O$ + 0.1% formic acid-$d_2$. **f**, $^{1}H$-$^{13}C$ HMBC spectrum acquired in 90%:10% (*v/v*) $H_2O$:$D_2O$, both containing 0.1% formic acid-$d_2$. **g**, $^{1}H$-$^{13}C$ HMBC spectrum acquired in 99.9% $D_2O$ + 0.1% formic acid-$d_2$.

b

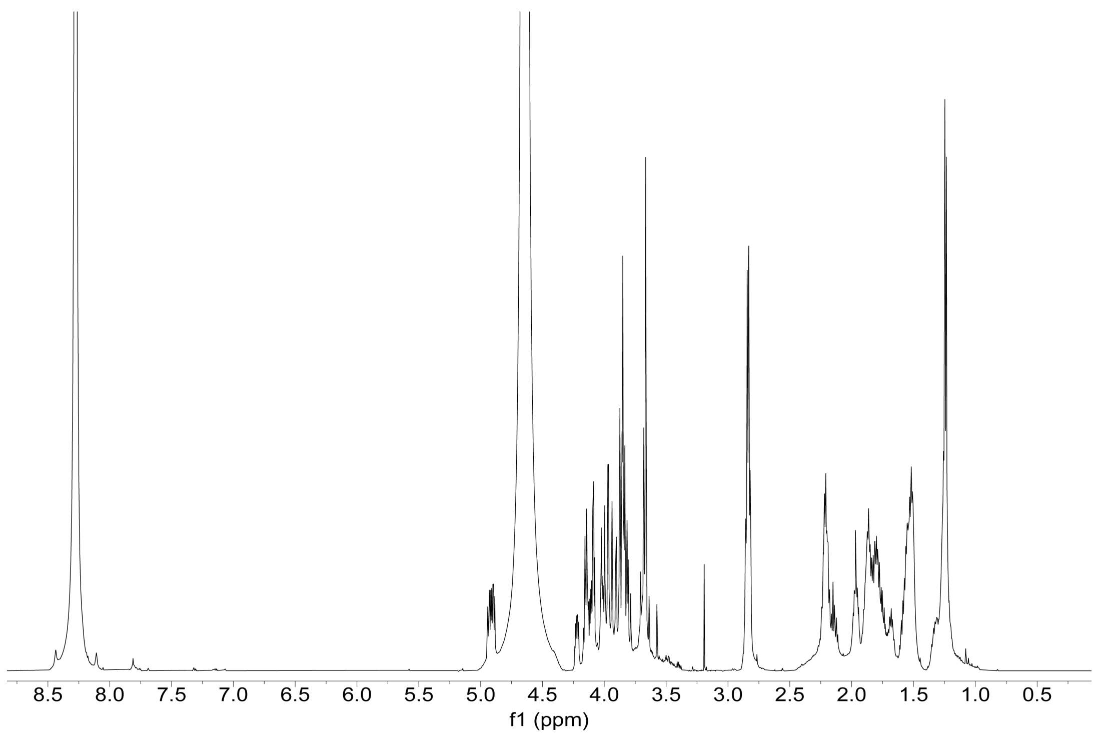


c

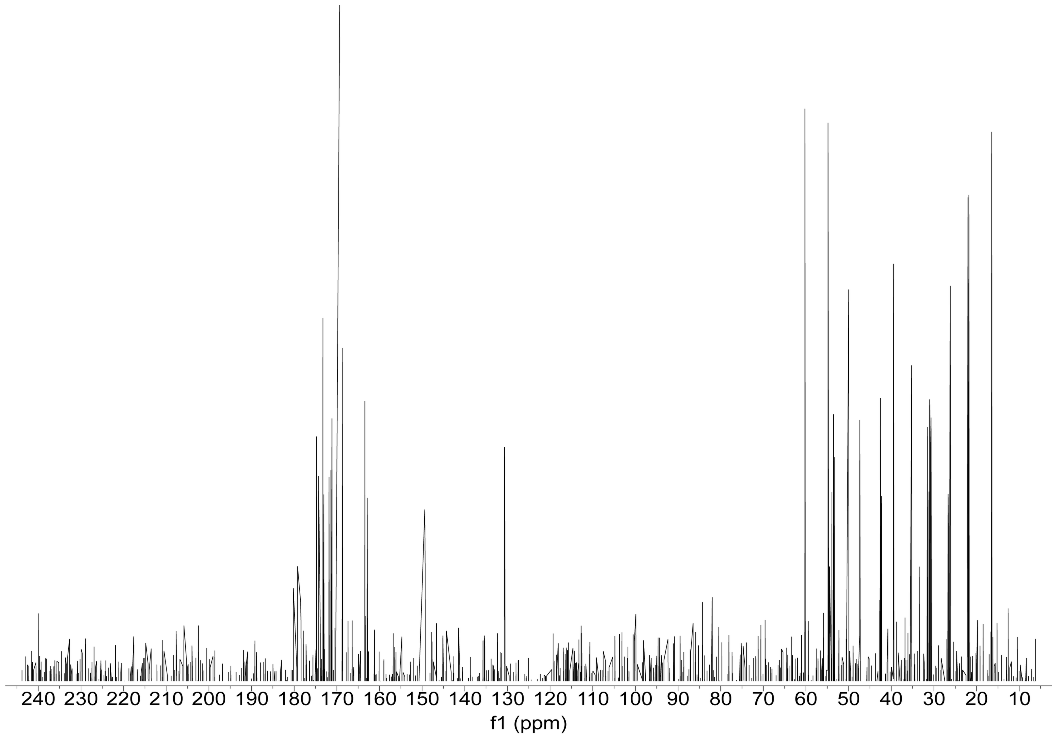


**Figure S9 (cont.). NMR spectra for FonA$_{67\text{-}80}$.**

**d**

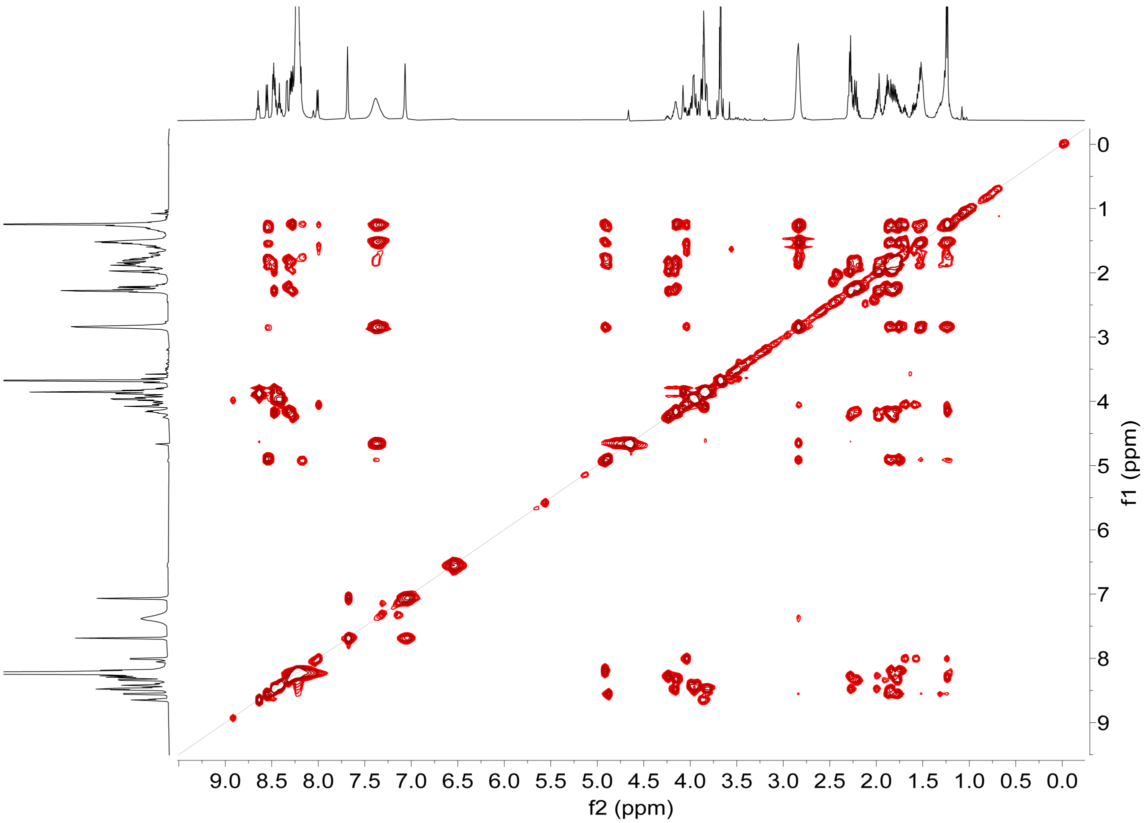


**e**

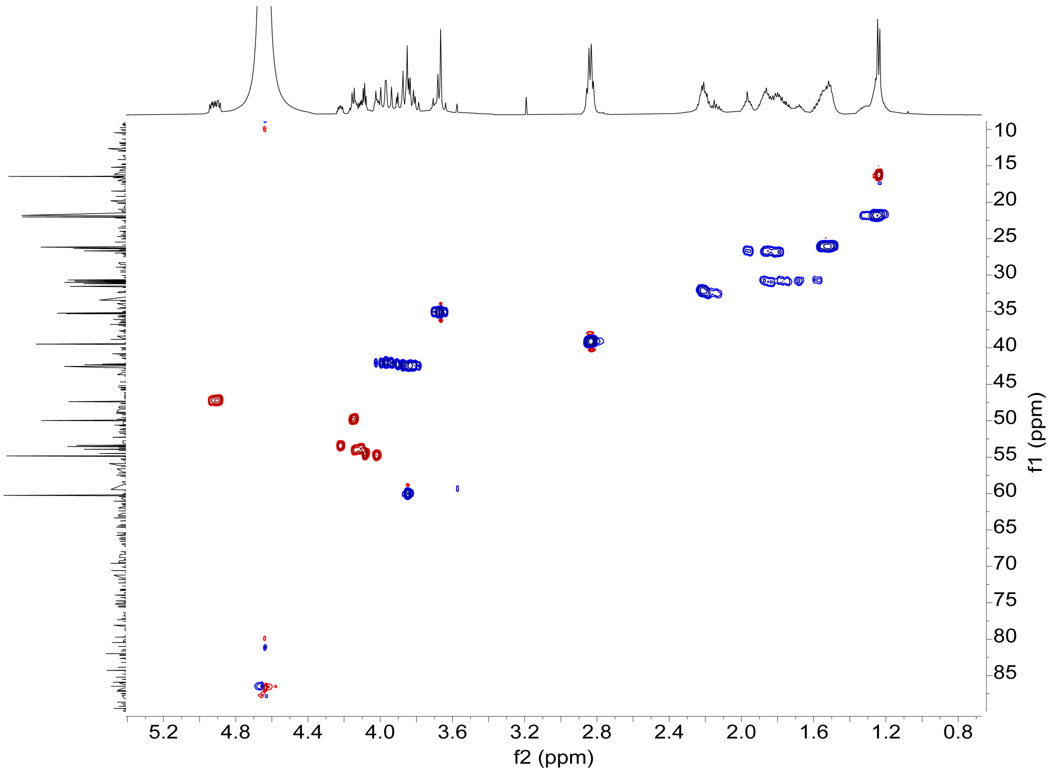


**Figure S9 (cont.). NMR spectra for FonA$_{67\text{-}80}$.**

f

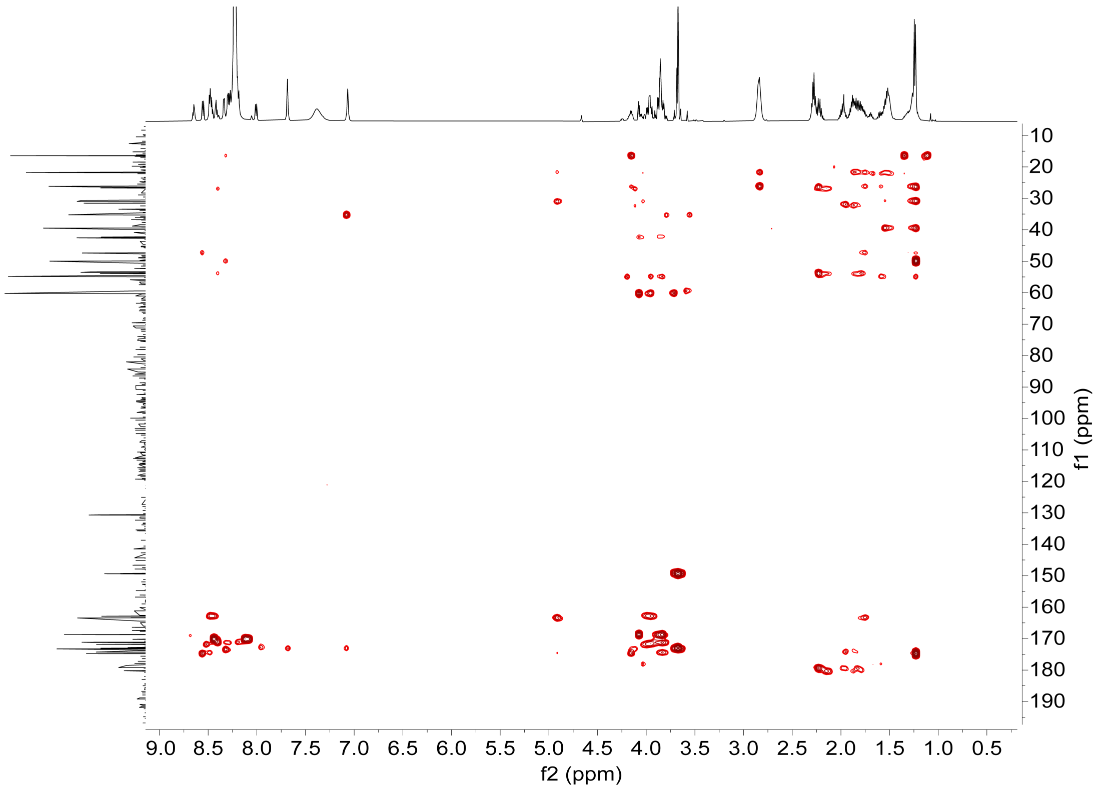


g

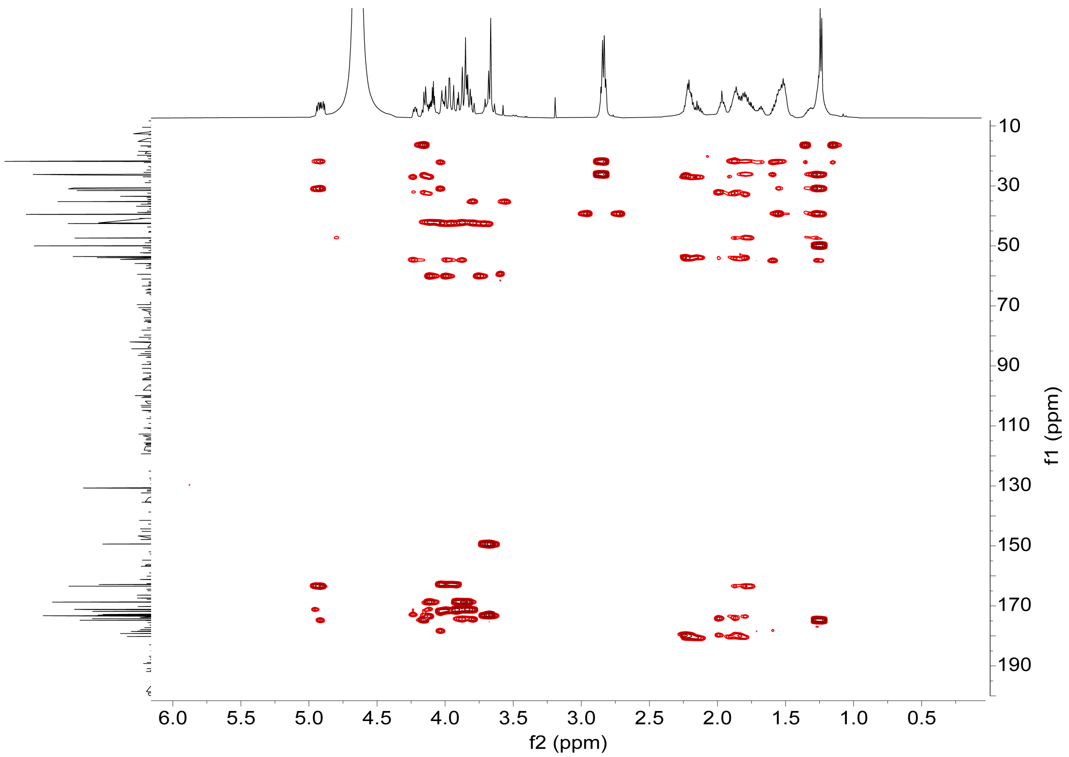


**Figure S9 (cont.). NMR spectra for FonA$_{67\text{-}80}$.**

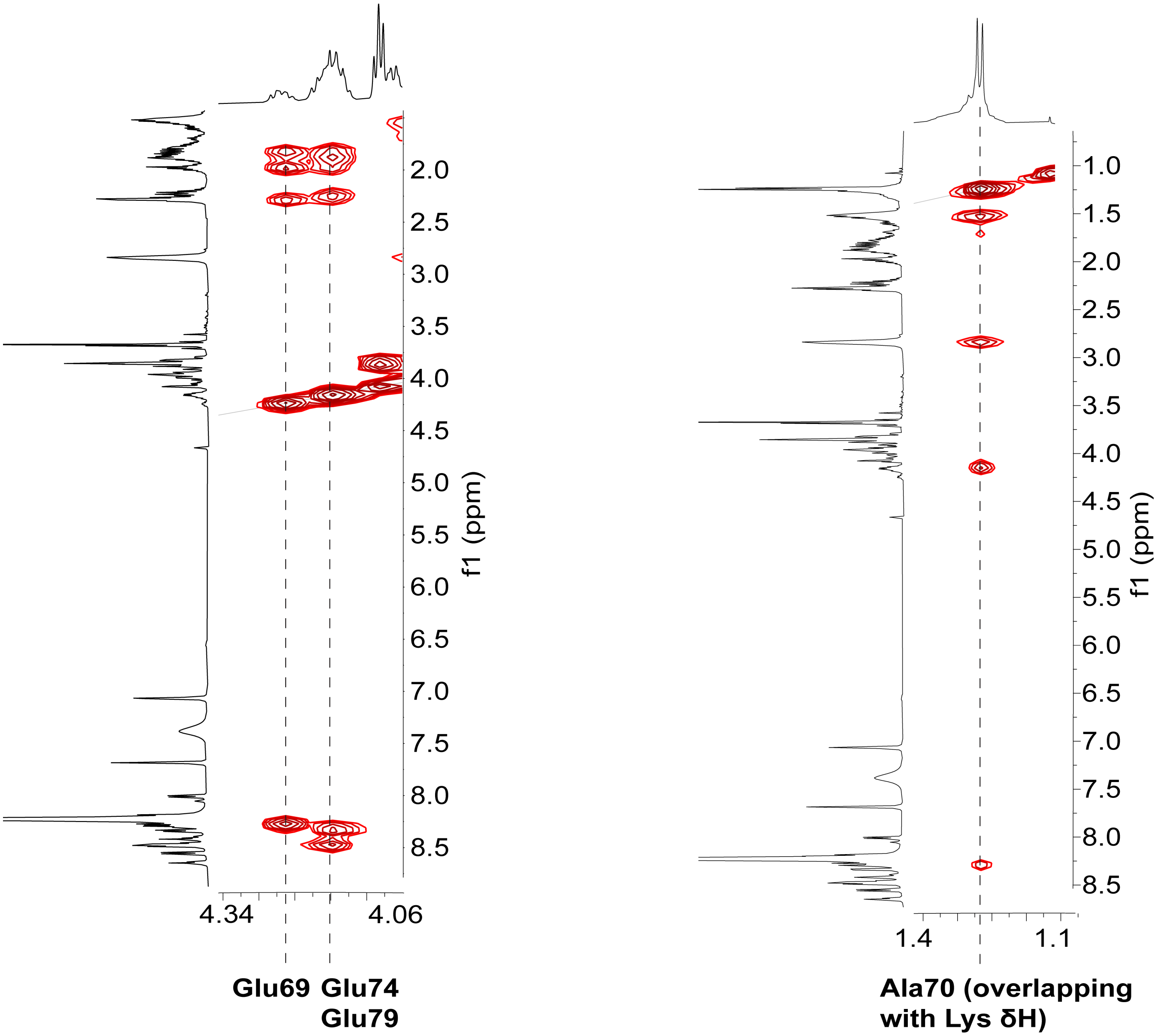


**Figure S10. Spin system assignments for unmodified residues in FonA$_{67\text{-}80}$.** All three Glu residues are observed as unmodified, with Glu74 and Glu79 seen to be slightly upfield-shifted due to being in closer proximity to the modified Cys-Gly motif. Ala70 is also observed. The remaining unmodified Ser and Gly residues undergo extensive signal overlap and cannot be differentiated.

a

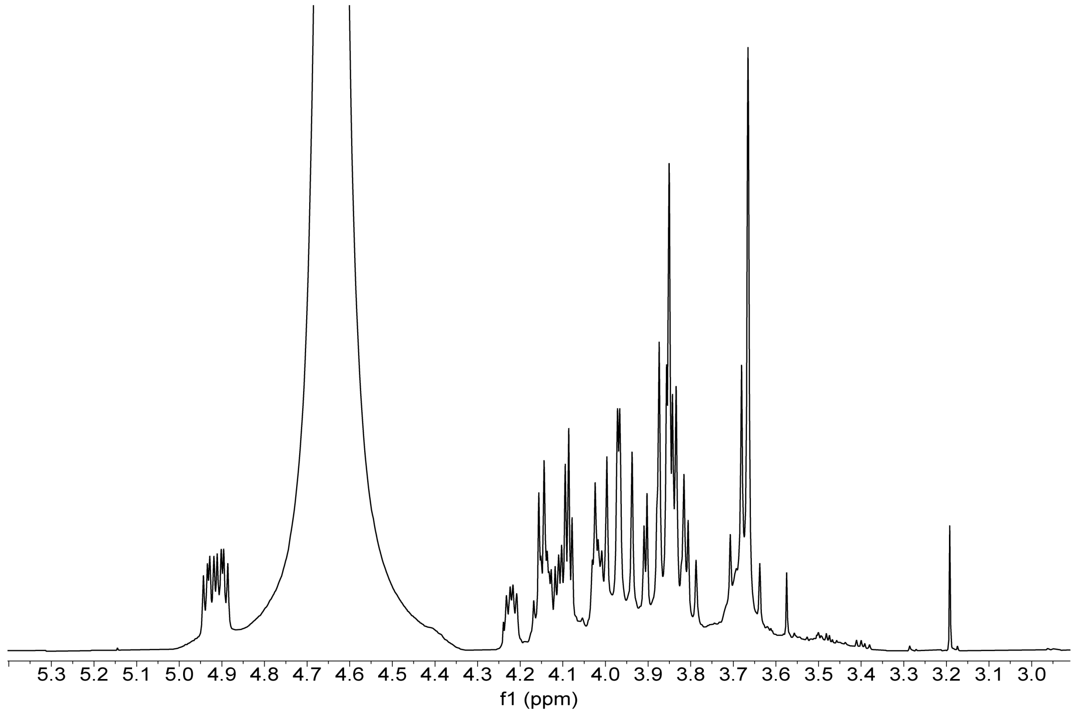


b

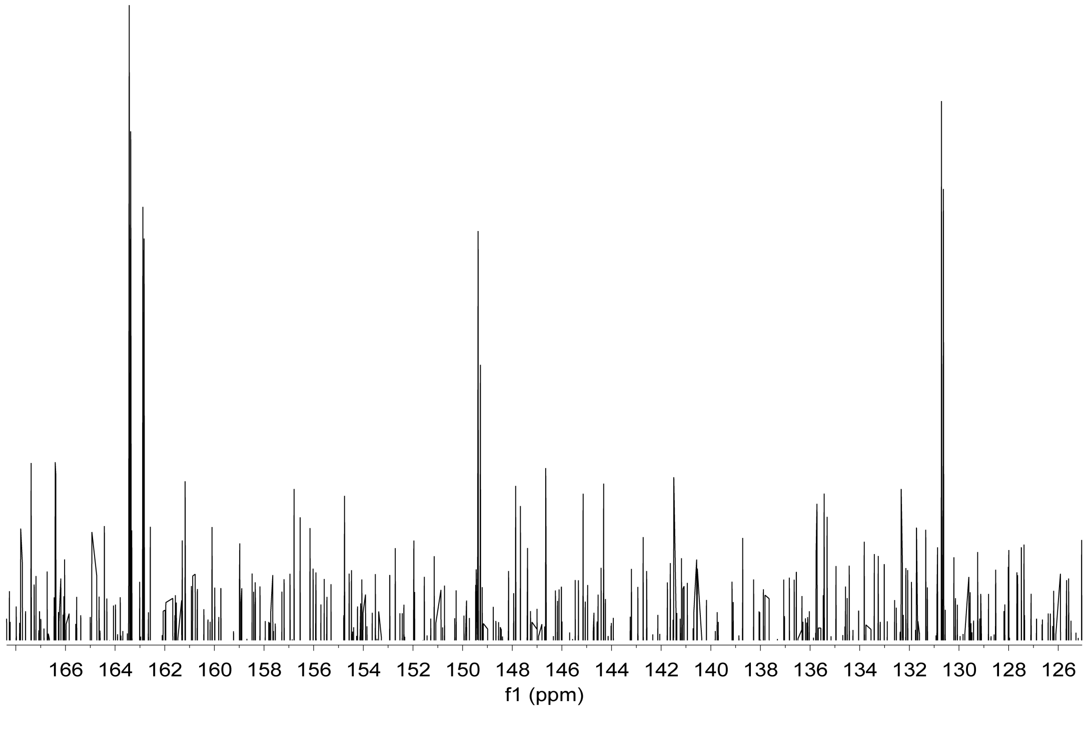


**Figure S11. Zoom-in showing anomalous FonA$_{67-80}$ $^{1}$H and $^{13}$C signals. a**, $^{1}$H NMR showing a set of downfield-shifted protons in the α proton region at 4.9 ppm. **b**, $^{13}$C NMR showing four $^{13}$C signals between 130-165 ppm, which do not belong to any canonical amino acids of the FonA$_{67-80}$ sequence.

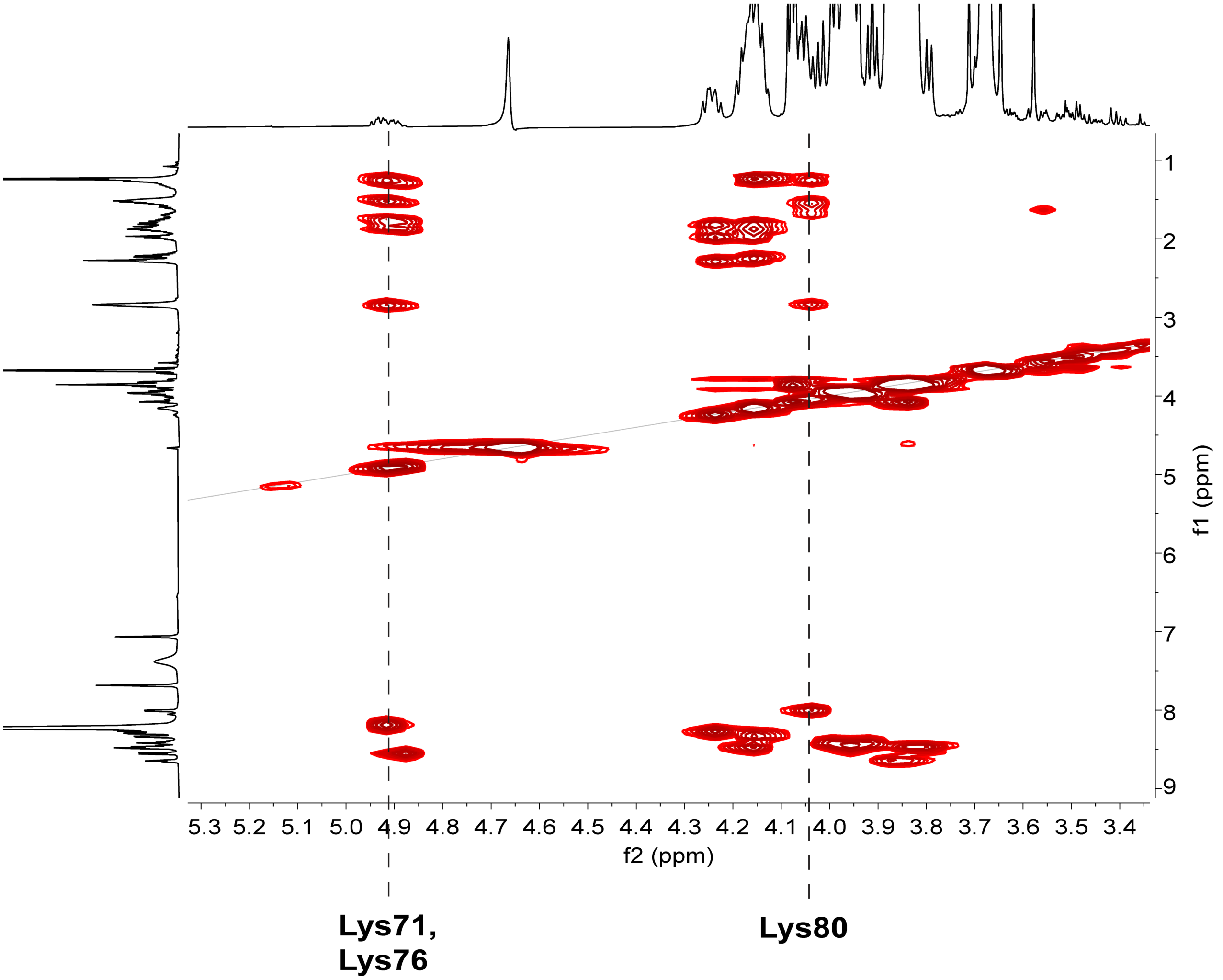


**Figure S12. Assignment/differentiation of FonA$_{67\text{-}80}$ Lys residues.** Correlations for three Lys side chains are shown, with two that are adjacent to the Cys-Gly motifs (Lys71 and Lys76) exhibiting downfield α protons.

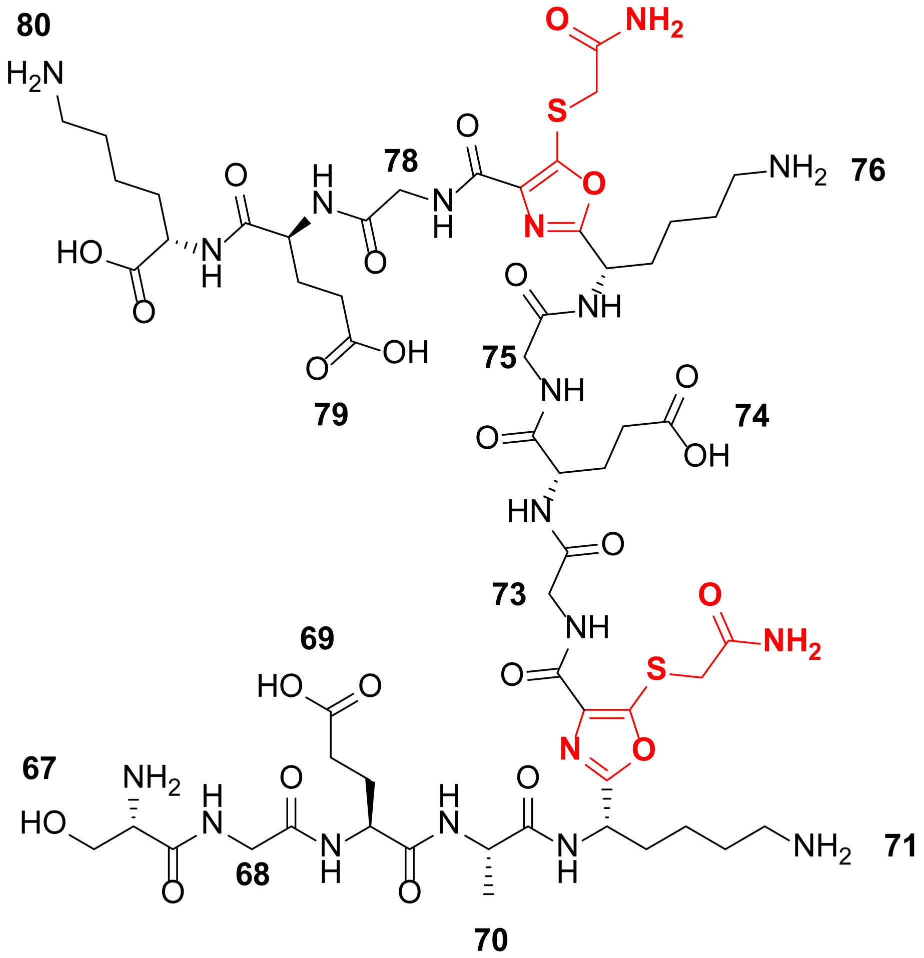


**Figure S13. FonA$_{67-80}$ structure.** Full structure of the modified, doubly alkylated FonA$_{67-80}$ peptide sequence. 5TO moieties (drawn in red) formed at Cys72/Gly73 and Cys77/Gly78 sites are drawn with corresponding acetamide alkylation of their free thiols.

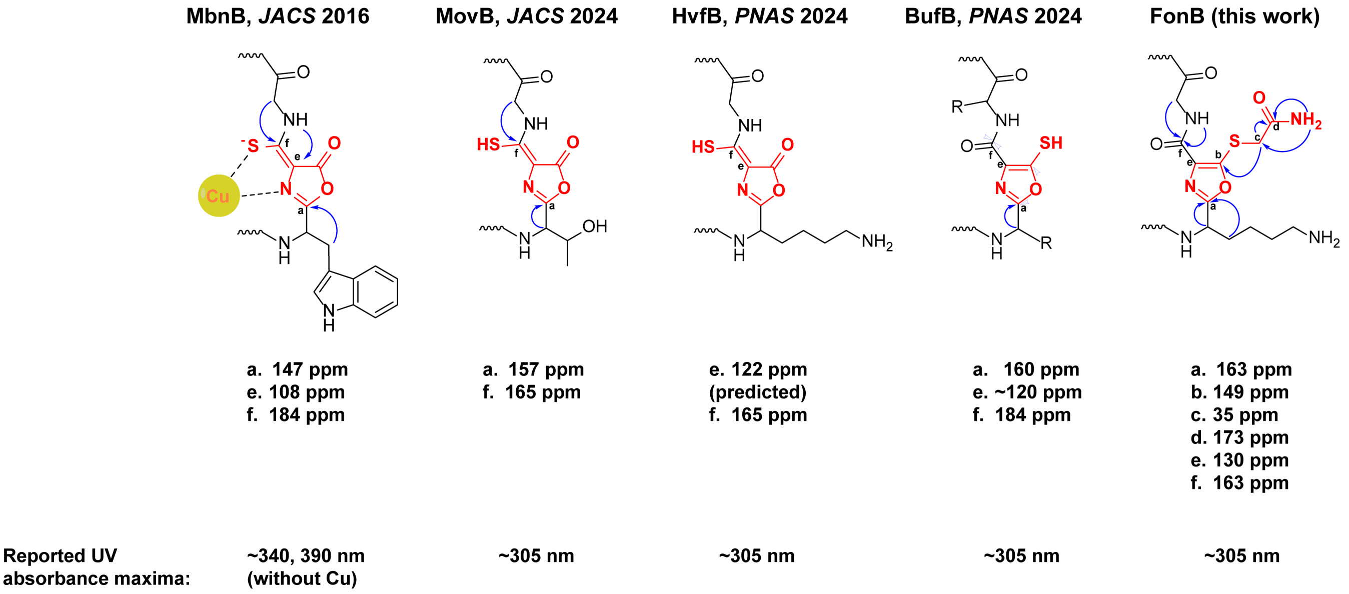


**Figure S14. Comparison of NMR and UV absorbance data for MNIO products.** $^{13}$C chemical shifts and UV absorbances of previously published heterocyclic MNIO products[11–14] are reported for comparison with the product of FonB (this work). Post-translational modifications are indicated in red.

**a**

| | FonB | HvfB | NgoB | BufB$_2$ | BufB$_1$ | MbnB | MovB |
|---|---|---|---|---|---|---|---|
| FonB | 100 | | | | | | |
| HvfB | 48 | 100 | | | | | |
| NgoB | 47 | 70 | 100 | | | | |
| BufB$_2$ | 39 | 38 | 37 | 100 | | | |
| BufB$_1$ | 36 | 33 | 34 | 43 | 100 | | |
| MbnB | 20 | 21 | 20 | 23 | 23 | 100 | |
| MovB | 17 | 21 | 20 | 22 | 20 | 34 | 100 |

| | FonB | HvfB | NgoB | BufB$_2$ | BufB$_1$ | MbnB | MovB |
|---|---|---|---|---|---|---|---|
| FonB | 100 | | | | | | |
| HvfB | 58 | 100 | | | | | |
| NgoB | 57 | 75 | 100 | | | | |
| BufB$_2$ | 49 | 48 | 47 | 100 | | | |
| BufB$_1$ | 48 | 45 | 46 | 51 | 100 | | |
| MbnB | 34 | 33 | 34 | 33 | 33 | 100 | |
| MovB | 31 | 33 | 32 | 35 | 34 | 34 | 100 |

**b**

| | FonC | HvfC | NgoC | BufC$_2$ | BufC$_1$ | MbnC | MovC |
|---|---|---|---|---|---|---|---|
| FonC | 100 | | | | | | |
| HvfC | 14 | 100 | | | | | |
| NgoC | 17 | 32 | 100 | | | | |
| BufC$_2$ | 15 | 16 | 15 | 100 | | | |
| BufC$_1$ | 19 | 14 | 12 | 23 | 100 | | |
| MbnC | 10 | 12 | 10 | 13 | 8 | 100 | |
| MovC | 13 | 12 | 13 | 11 | 9 | 14 | 100 |

| | FonC | HvfC | NgoC | BufC$_2$ | BufC$_1$ | MbnC | MovC |
|---|---|---|---|---|---|---|---|
| FonC | 100 | | | | | | |
| HvfC | 25 | 100 | | | | | |
| NgoC | 27 | 41 | 100 | | | | |
| BufC2 | 25 | 26 | 24 | 100 | | | |
| BufC1 | 28 | 24 | 21 | 29 | 100 | | |
| MbnC | 19 | 21 | 18 | 18 | 13 | 100 | |
| MovC | 23 | 22 | 22 | 17 | 17 | 24 | 100 |

**Figure S15. Comparison of MNIO and PME1 proteins.** Pairwise similarity and identity values are shown for relevant **a**, MNIO proteins and **b**, PME1 proteins: Top matrix, % identity; bottom matrix, % similarity. MbnC and MovC are not retrieved by the DUF2063 HMM.

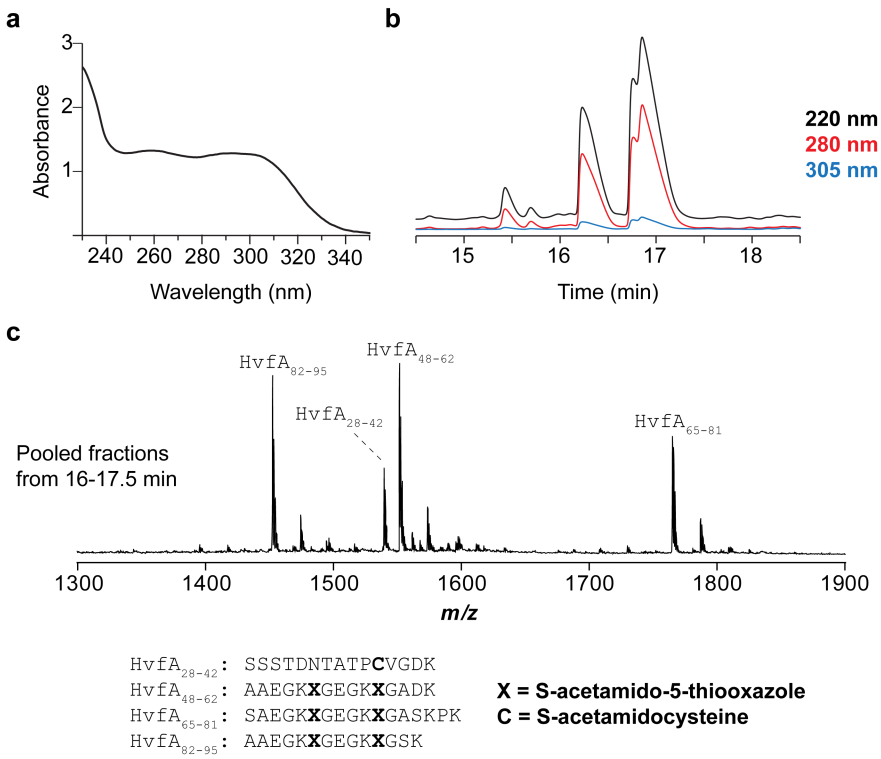


**Figure S16. Purification and alkylation of oxazolin peptides. a**, UV-Vis absorbance spectrum for full length oxazolin product purified by IMAC. Baseline subtraction was performed using the UV-Vis absorbance spectrum of buffer alone. **b**, HPLC chromatogram for oxazolin peptides following alkylation and digestion. **c**, MALDI-TOF-MS for fractions containing oxazolin peptides of interest.

**a**

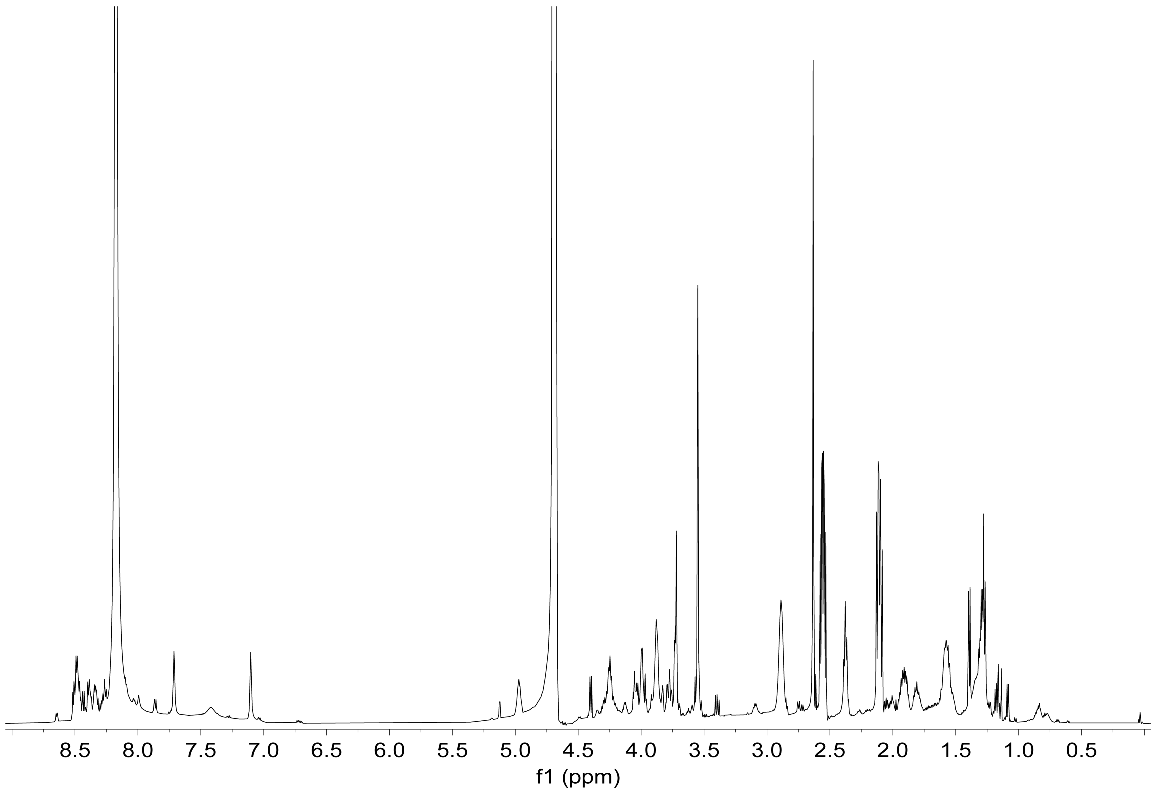


**Figure S17. NMR spectra of the proteolytically digested oxazolin sample.** **a**, $^{1}H$ NMR spectrum acquired in 90%:10% (*v/v*) $H_2O$:$D_2O$, both containing 0.1% formic acid-$d_2$. **b**, $^{1}H$-$^{13}C$ HMBC spectrum acquired in 90%:10% (*v/v*) $H_2O$:$D_2O$, both containing 0.1% formic acid-$d_2$. **c**, Key HMBC correlations which correspond to the same correlations observed in the $FonA_{67\text{-}80}$ NMR spectra.

b

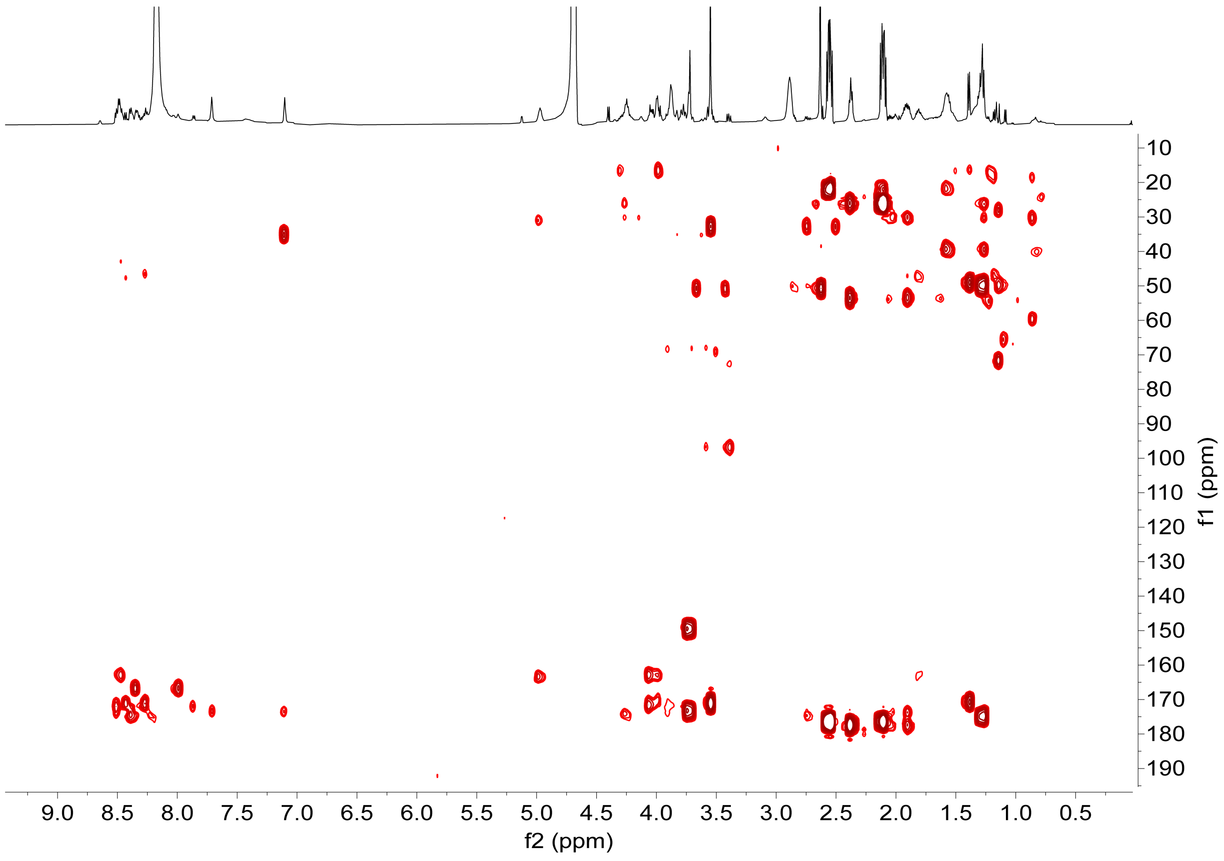


c

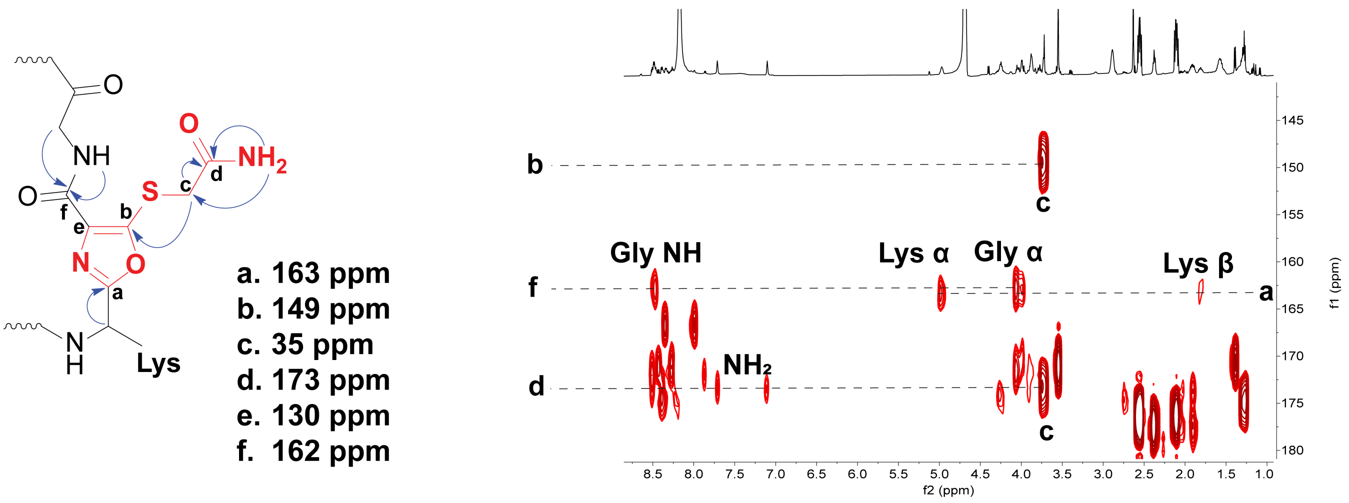


**Figure S17 (cont.). NMR spectra of the proteolytically digested oxazolin sample.**

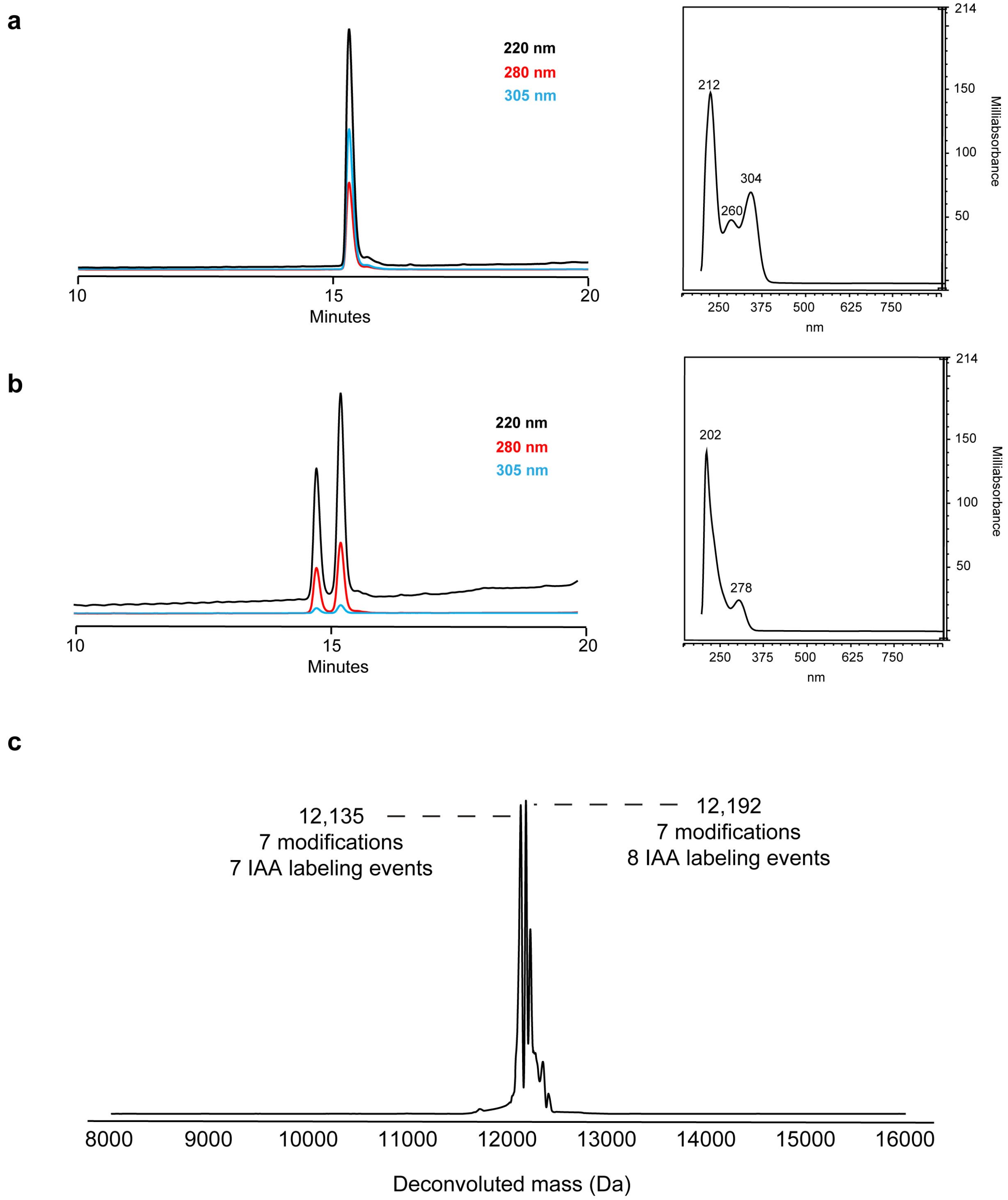


**Figure S18. Gonophorin UV and MS data. a**, HPLC chromatogram of His$_6$-tagged NgoA co-expressed with NgoB and NgoC, which displays strong absorbance at 305 nm. **b**, HPLC chromatogram of His$_6$-NgoABC after treatment with iodoacetamide. Acetamide labeling removes the 305 nm absorbance. **c**, Deconvoluted mass spectrum of IAA-treated His$_6$-NgoABC. All 7 Cys residues were found to be modified and alkylated, with a major species containing an 8$^{th}$ alkylation also observed (likely on the N-terminus).

a

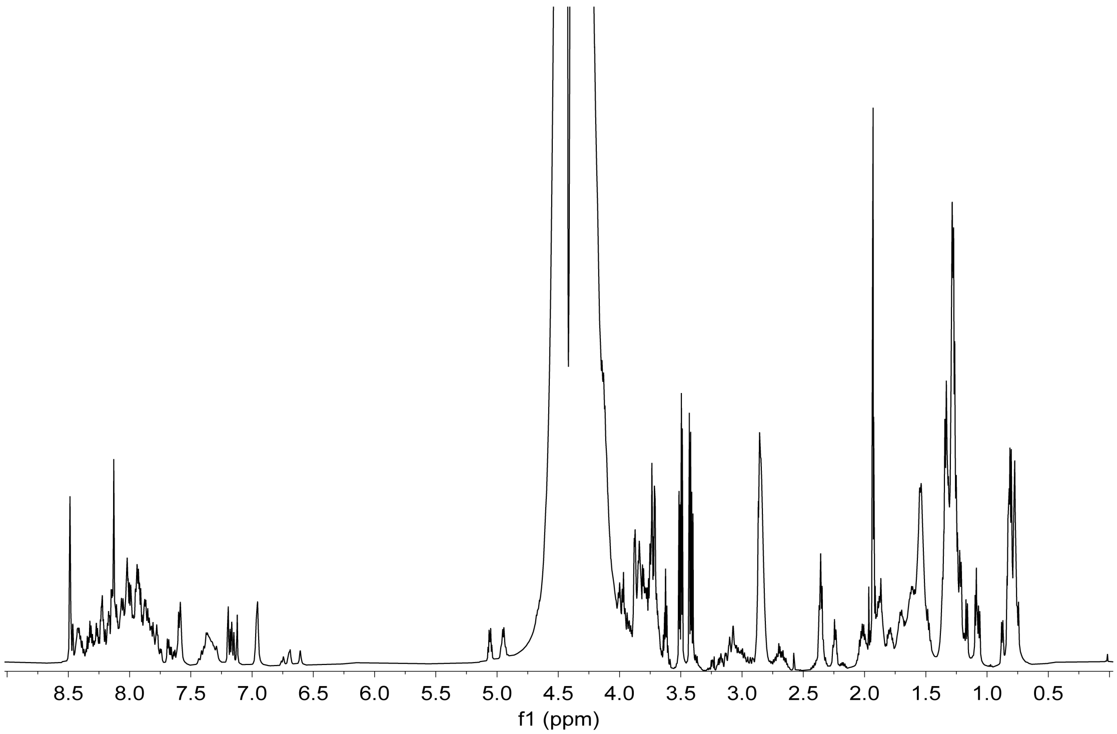


**Figure S19. Gonophorin NMR data.** **a**, $^{1}$H NMR spectrum of full-length modified and acetamide-alkylated NgoABC acquired in 90%:10% (*v/v*) $H_2O$:$D_2O$, both containing 0.1% formic acid-$d_2$. **b**, $^{1}$H-$^{13}$C HMBC spectrum acquired in 90%:10% (*v/v*) $H_2O$:$D_2O$, both containing 0.1% formic acid-$d_2$. **c**, Key HMBC correlations which correspond to the same correlations observed in the $FonA_{67-80}$ NMR spectrum.

b

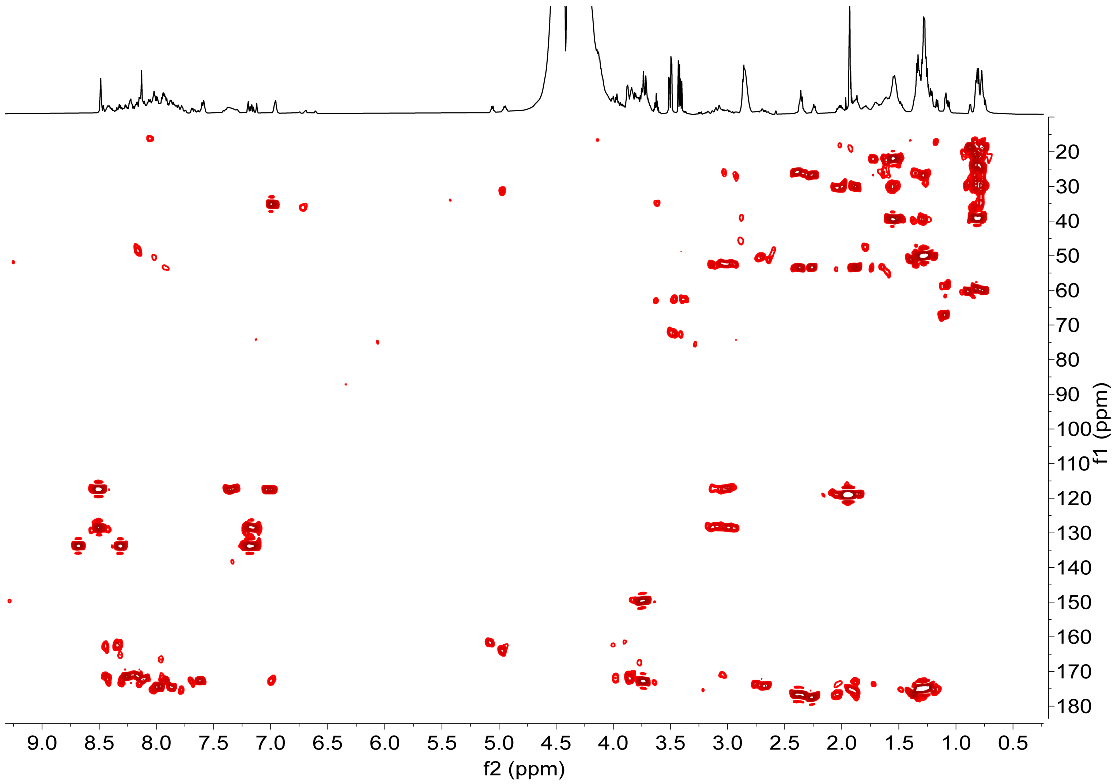


c

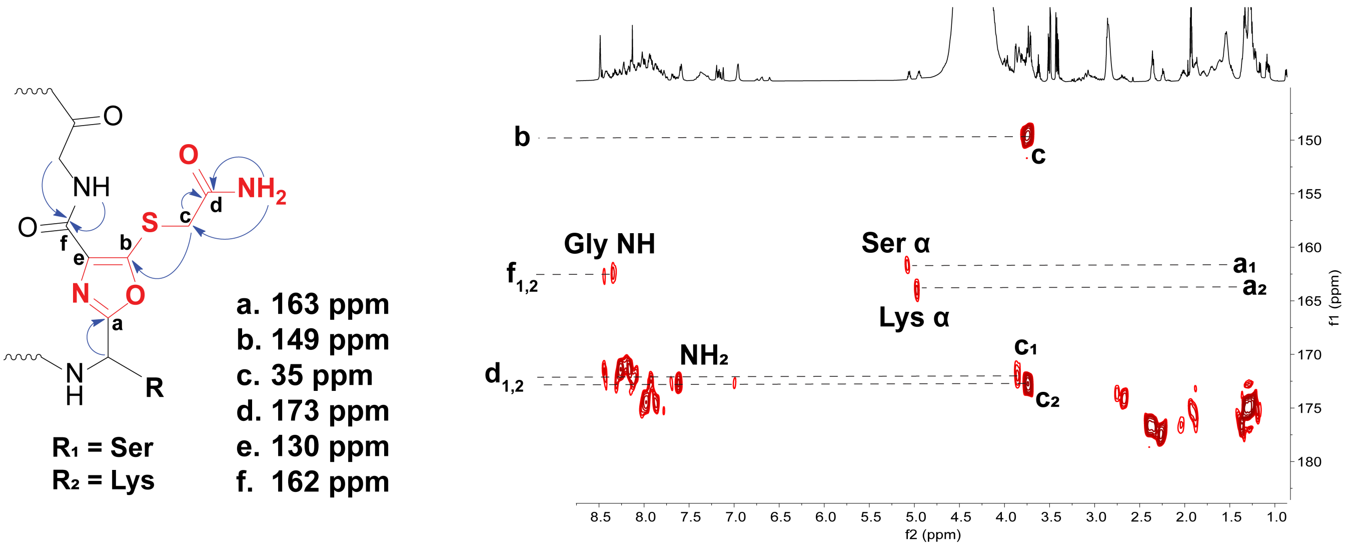


**Figure S19 (cont.). Gonophorin NMR data.**

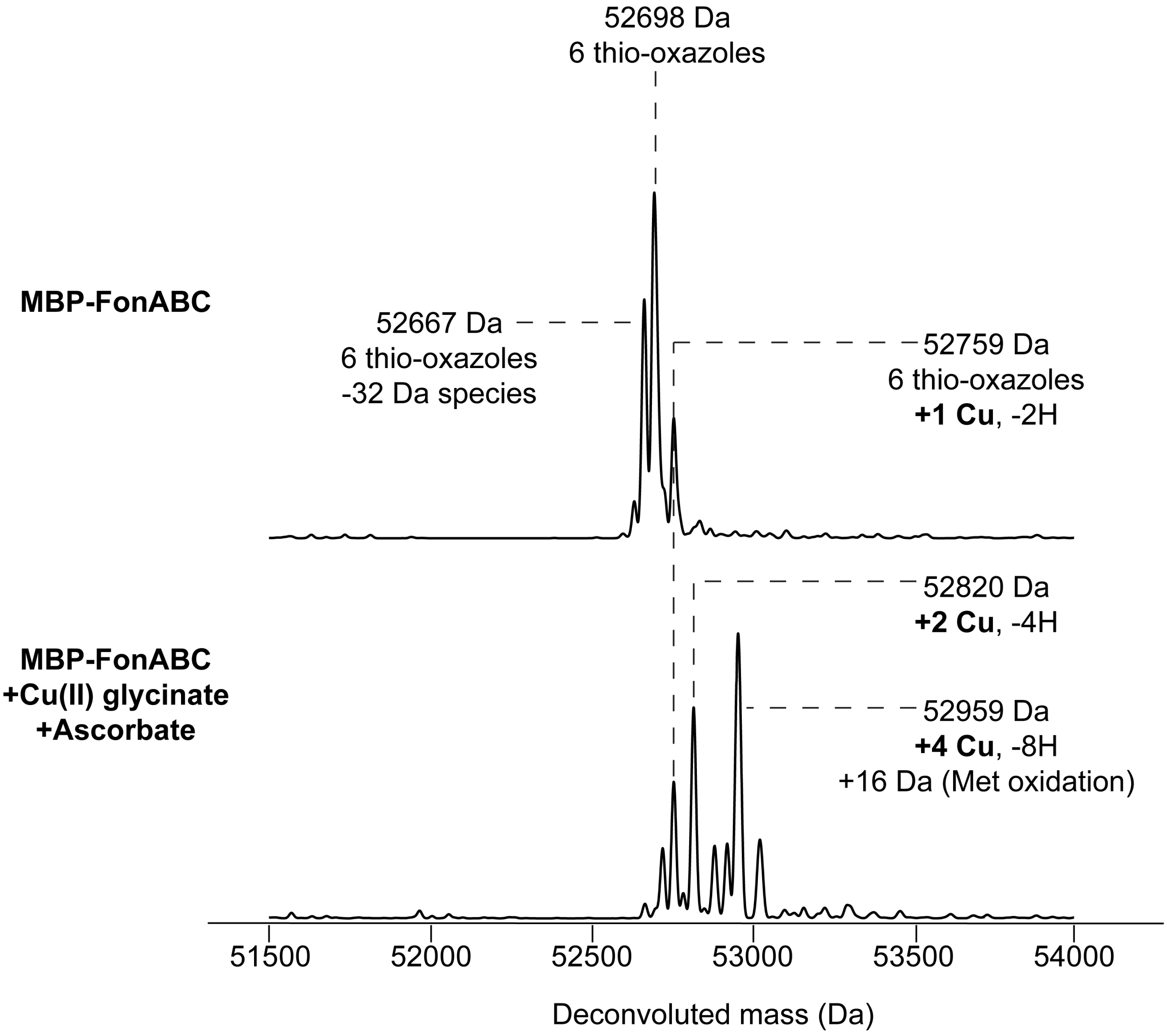


**Figure S20. Native MS of MBP-FonABC.** Deconvoluted high-resolution native mass spectra comparing MBP-FonABC and MBP-FonABC with added Cu(II) and sodium ascorbate.

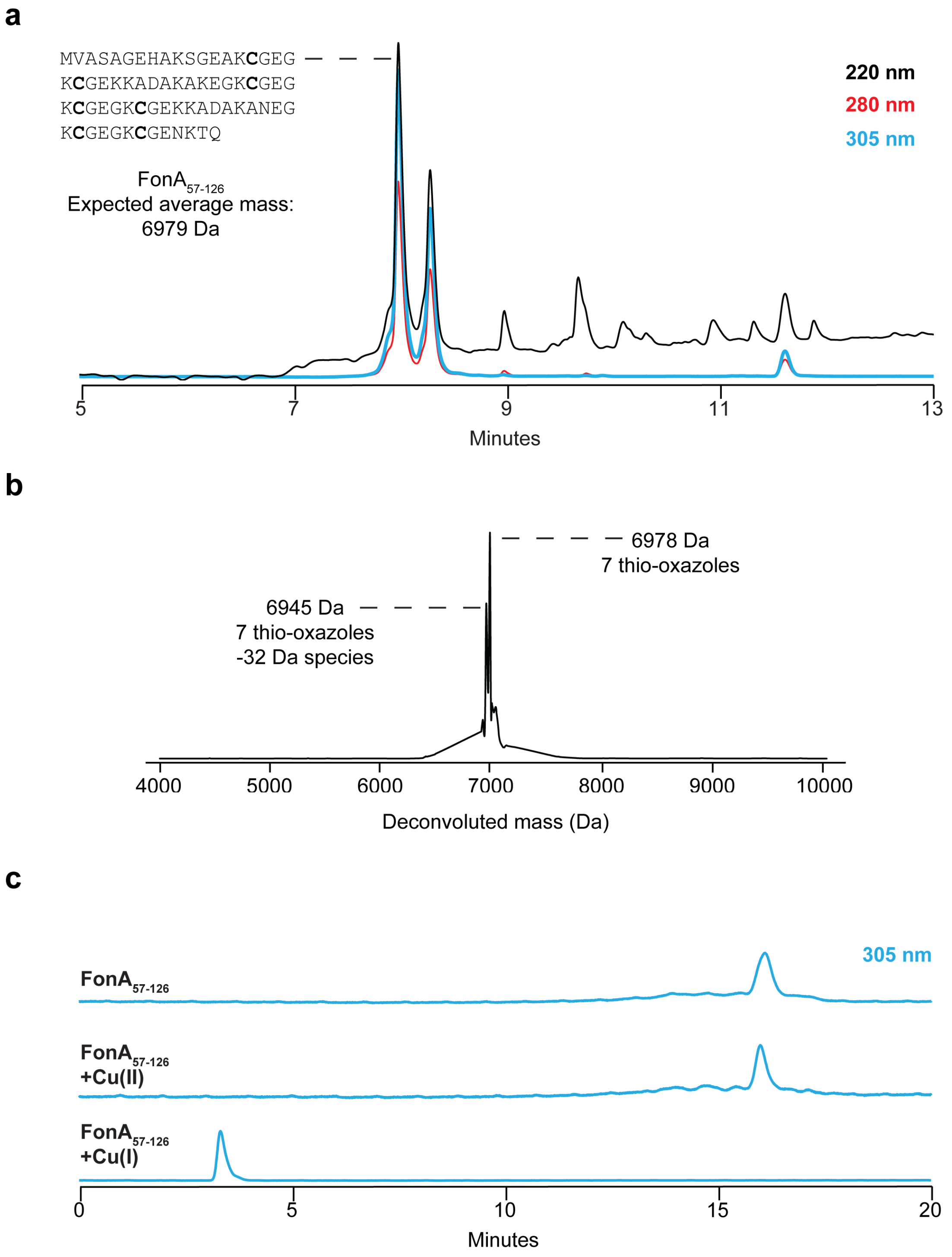


**Figure S21. LC retention time shift during fontiphorin Cu-binding assay.** **a**, HPLC chromatogram for purification of $\mathrm{FonA}_{57\text{-}126}$ following chymotrypsin digest. **b**, Deconvoluted mass spectrum of purified $\mathrm{FonA}_{57\text{-}126}$. **c**, Stacked HPLC chromatograms of $\mathrm{FonA}_{57\text{-}126}$ with Cu showing shift in retention time upon exposure to Cu(I).

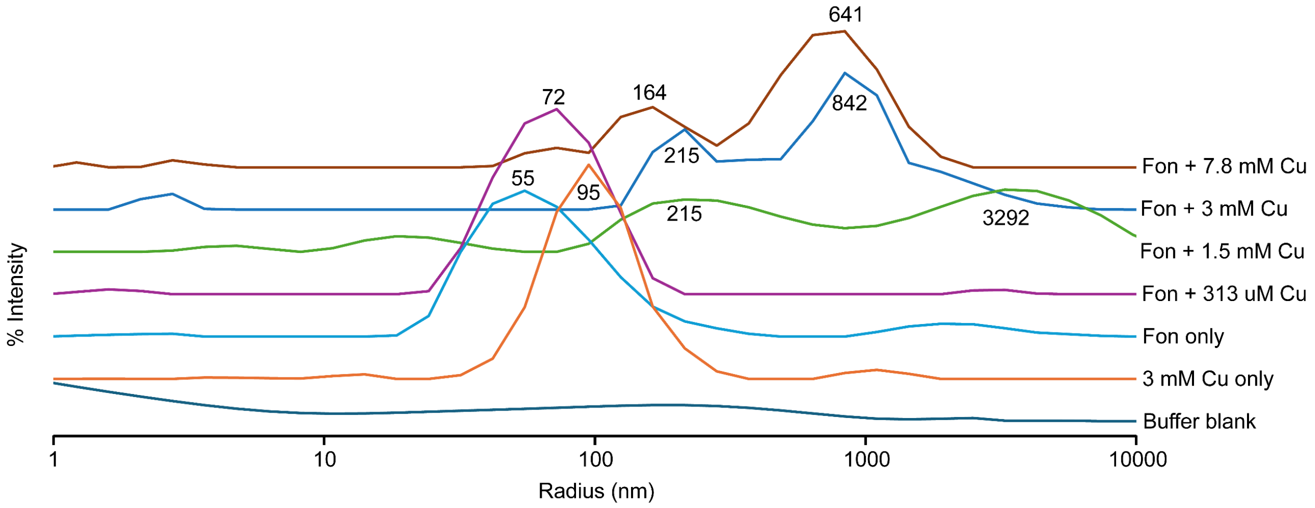


**Figure S22. Dynamic light scattering for fontiphorin.** Intensity-weighted size distribution curve is reported. Radii for local intensity maxima are listed for each curve. Measurements were obtained following laser auto-attenuation to avoid detector saturation, and each data point represents the average of three individual replicates.

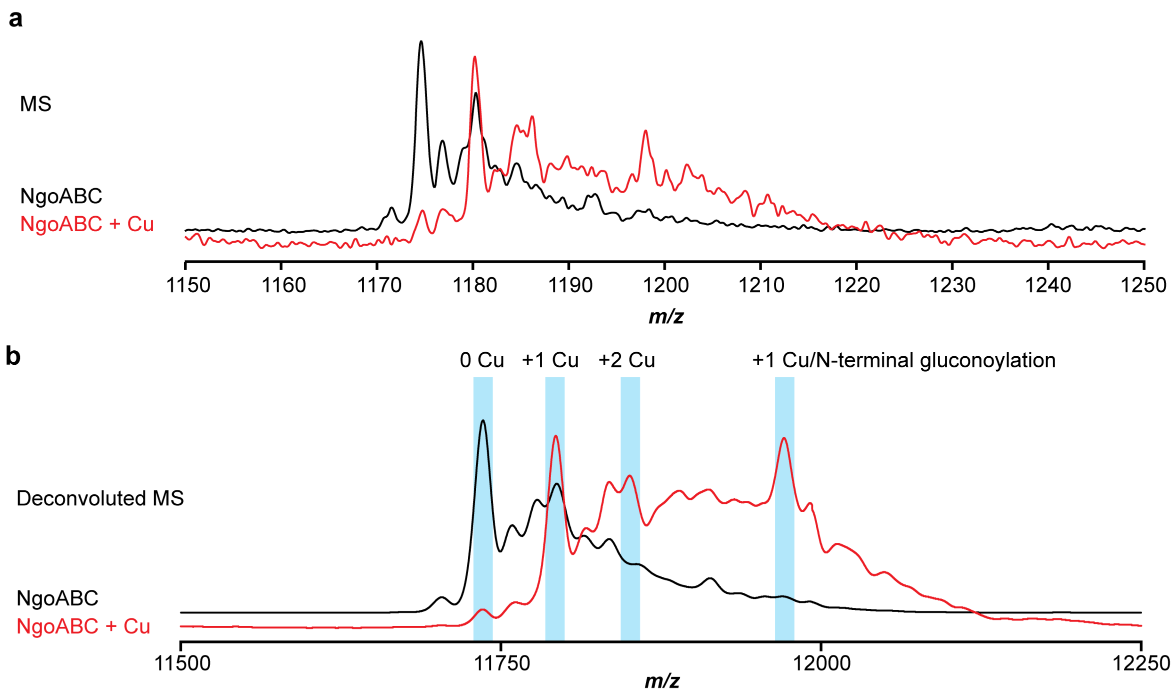


**Figure S23. LC-MS spectra of gonophorin with Cu loaded *in vivo*. a**, MS and **b**, deconvoluted MS for gonophorin with and without *in vivo* loaded Cu.

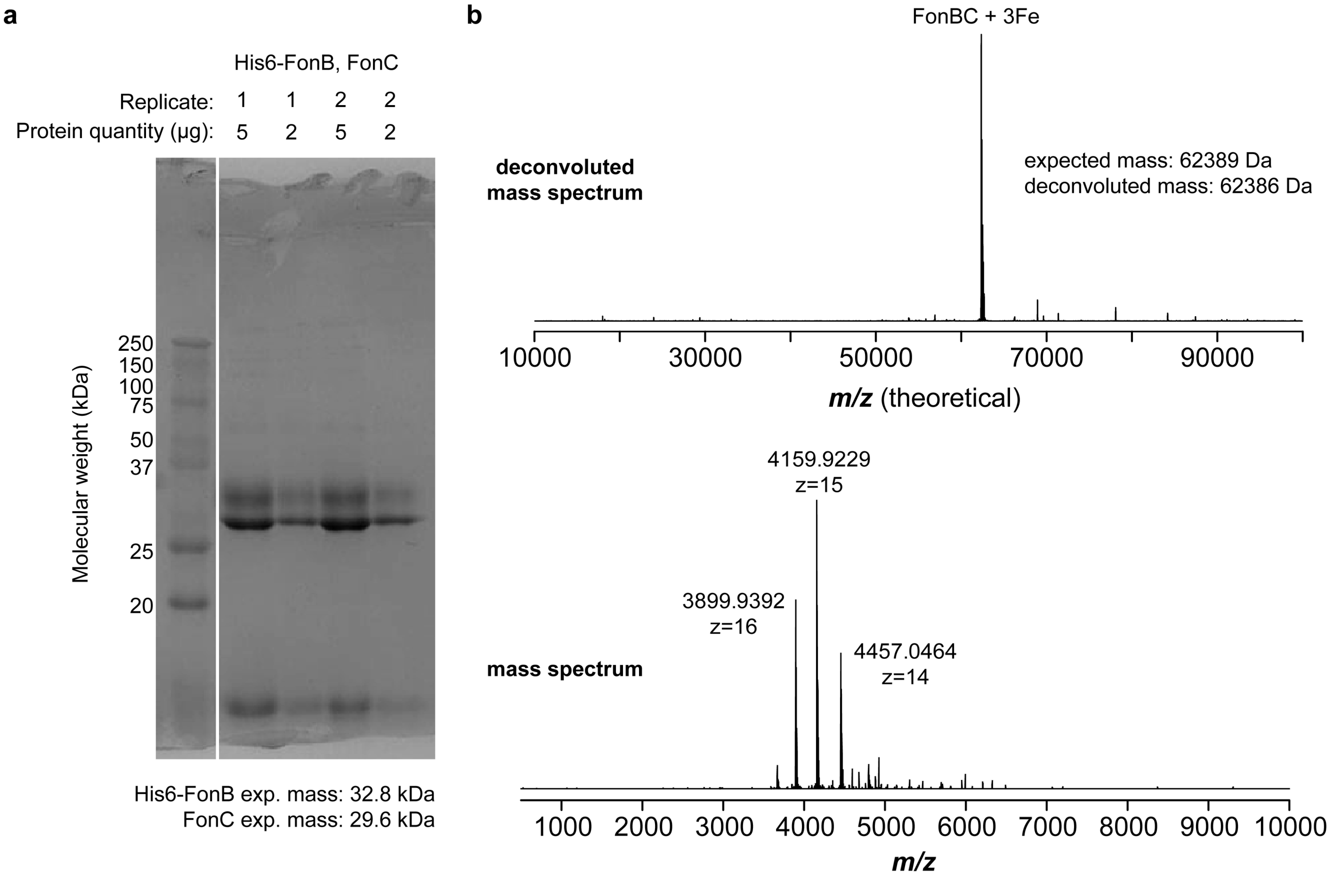


**Figure S24. Purification of FonBC dimer. a**, SDS-PAGE of purified His$_6$-FonBC. **b**, High-resolution native mass spectrum with corresponding deconvoluted mass spectrum shown.

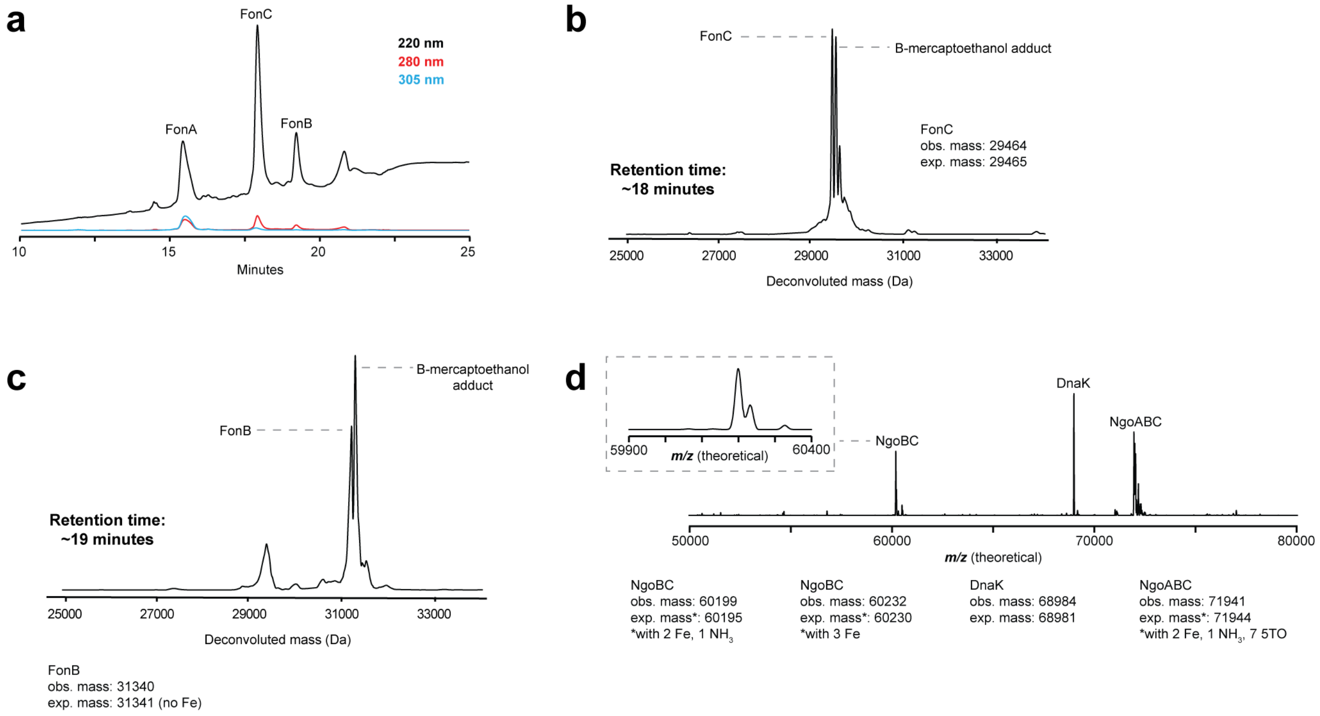


**Figure S25. FonA and NgoA co-purify with their respective MNIO and PME1. a**, HPLC trace of FonABC. **b**, Deconvoluted mass spectrum of FonC. **c**, Deconvoluted mass spectrum of FonC. **d**, Deconvoluted high-resolution mass spectrum of NgoABC.

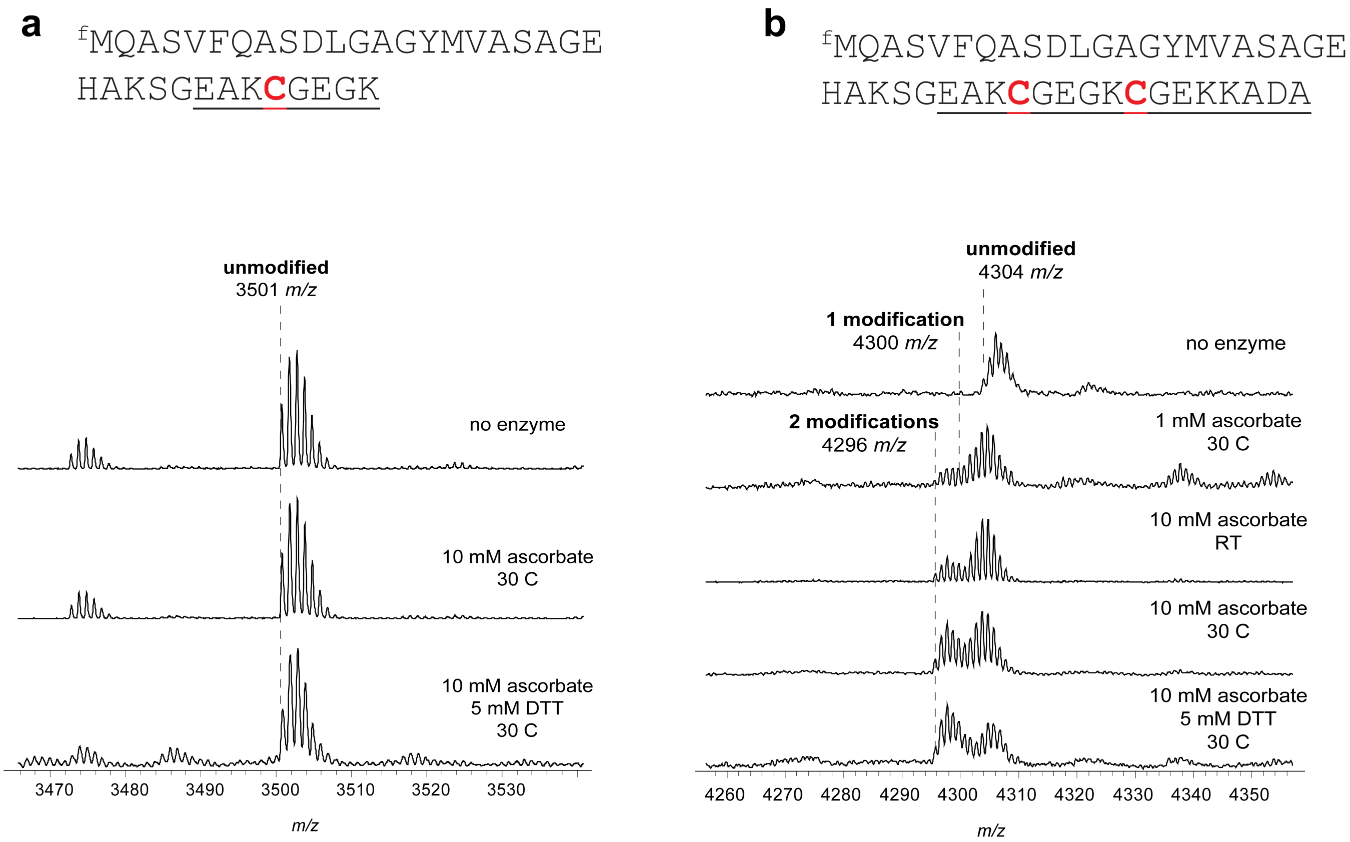


**Figure S26. Initial reconstitution of FonBC.** Example conditions tested during initial *in vitro* reaction condition optimization. Substrates tested; **a**, Peptide containing leader region with first repeat only. **b**, Peptide containing leader region with first two repeats. (Note: C-terminal Ala of the two Cys-containing substrate was removed for subsequent experiments). RT, room temperature.

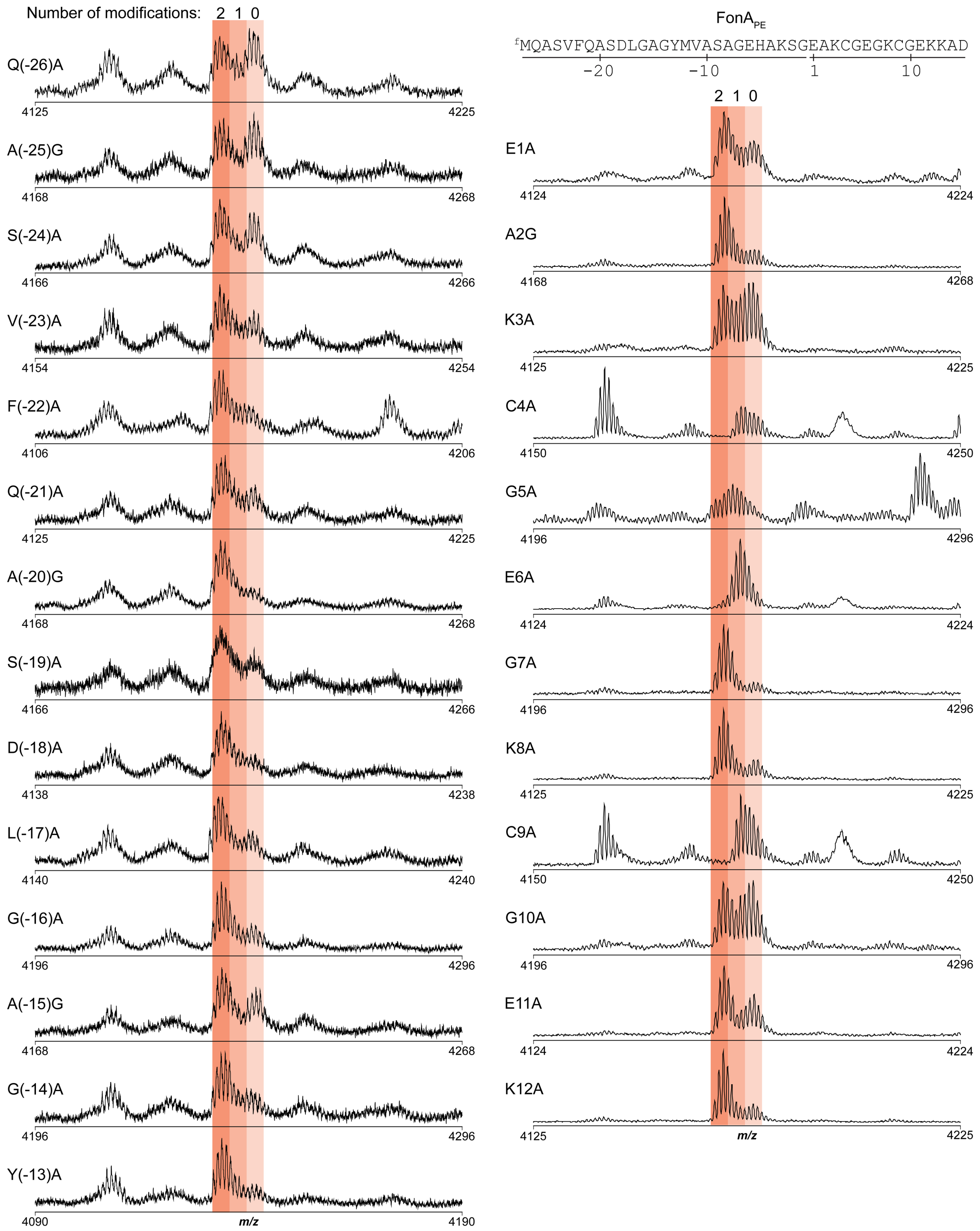


**Figure S27. FonB substrate tolerance for single-site variants.** Single-site Ala variants tested for leader and core region containing two repeats. Positions natively containing Ala were replaced with Gly.

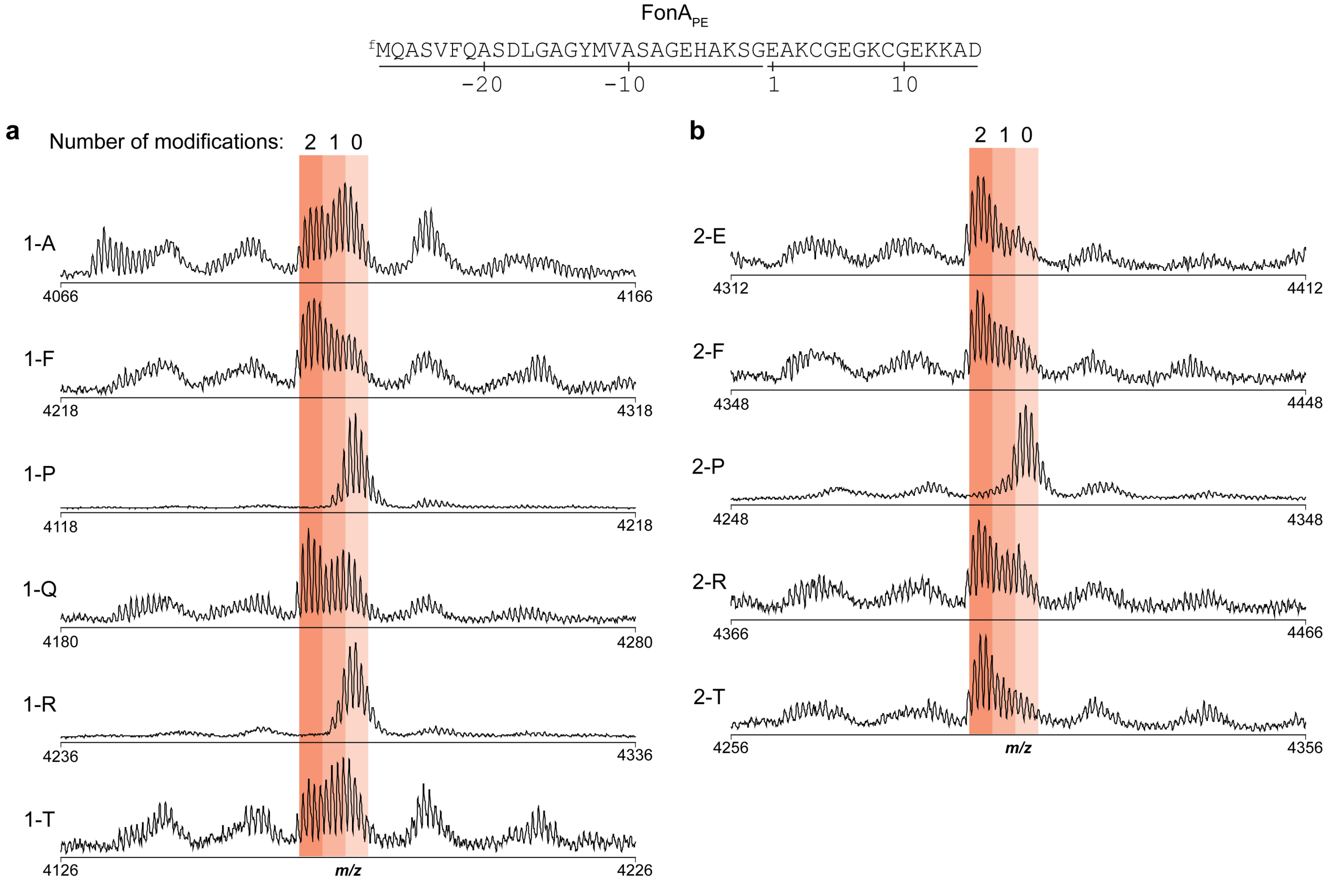


**Figure S28. FonB substrate tolerance for double-site variants.** MALDI-TOF-MS spectra of FonBC reactions on double-site FonA$_{PE}$ variants. Matching mutations were installed at repeat positions 1-5 (**a-e**, respectively). For example, FonA$_{PE}$1-A substitutes Ala for residues 1 and 6 of FonA$_{PE}$, yielding <u>A</u>AKCG<u>A</u>GKCG for core residues 1-10.

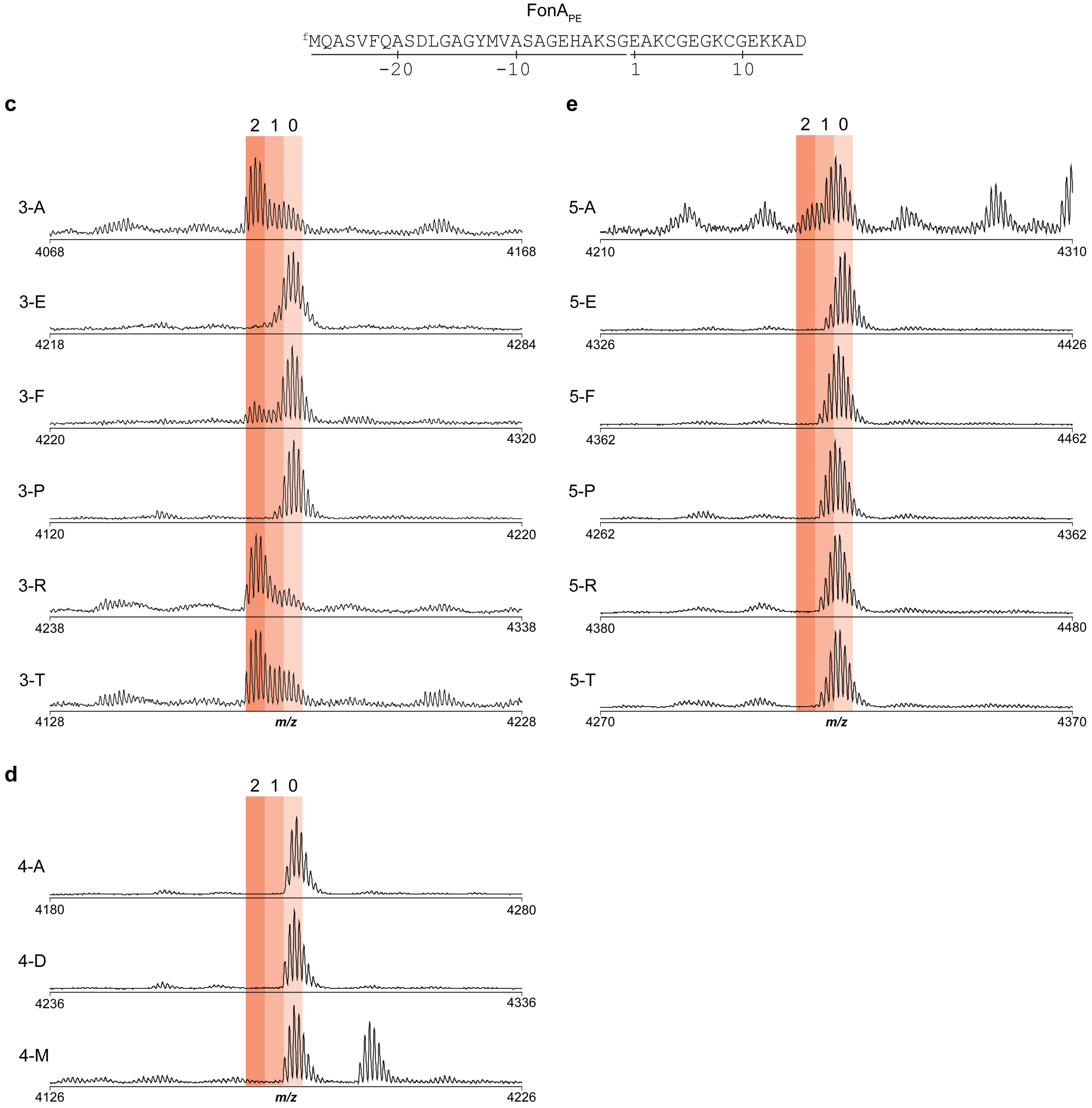


**Figure S28 (cont.). FonB substrate tolerance for double-site variants.**

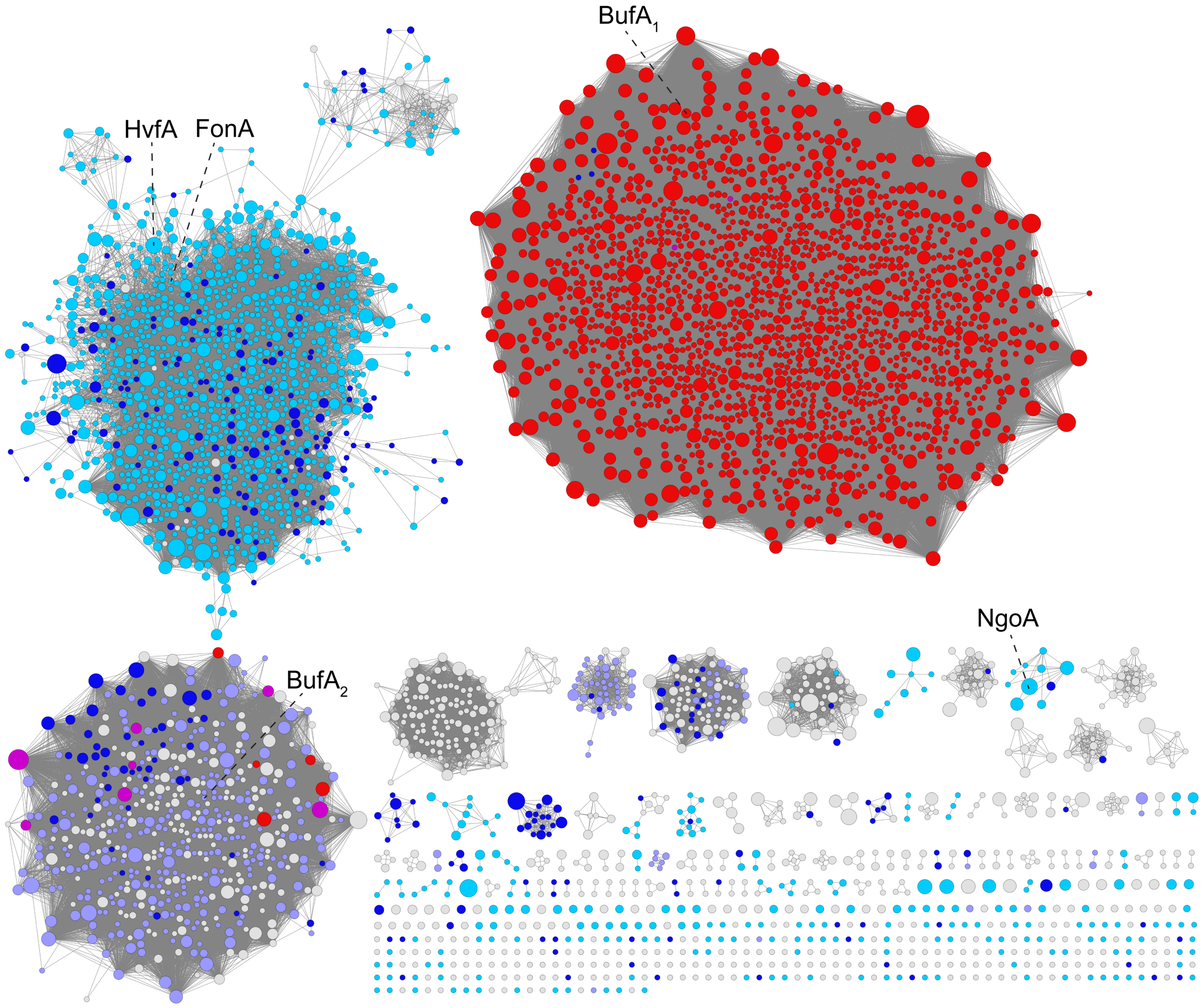


**Figure S29. Sequence similarity network for putative precursor peptides.** Network was generated at alignment score of 13. Nodes were conflated at >70% identity. Larger nodes represent larger numbers of conflated sequences. Color annotations: Red, match to DUF2282 HMM; Cyan, repeat motif containing Lys-Cys; Periwinkle, Repeat motif containing Cys-Lys; Dark blue, Any Cys-containing repeat motif; Purple, match to DUF2282 HMM and sequence has Cys-containing repeat motif.

**a**

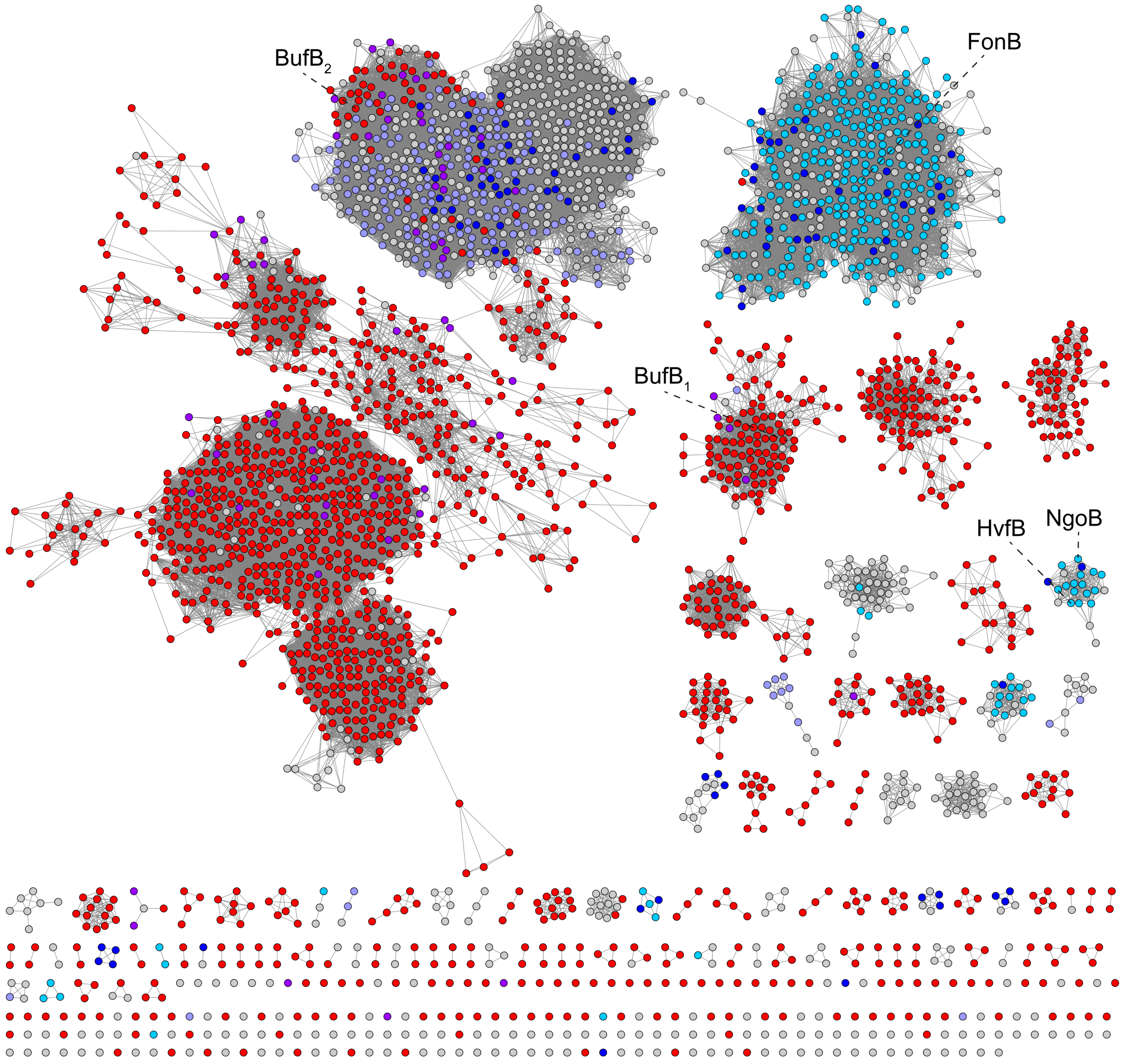


**Figure S30. Biosynthetic protein sequence similarity networks colored by precursor peptide family. a**, MNIO SSN (alignment score, 90; RepNode 0.60) colored by precursor peptide type. **b**, PME1 SSN (alignment score, 30; RepNode 0.70) colored by precursor peptide type. Color annotations: Red, match to DUF2282 HMM; Cyan, repeat motif containing “Lys-Cys”; Periwinkle, Repeat motif containing “Cys-Lys”; Dark blue, Any Cys-containing repeat motif; Purple, match to DUF2282 HMM and sequence has Cys-containing repeat motif.

**b**

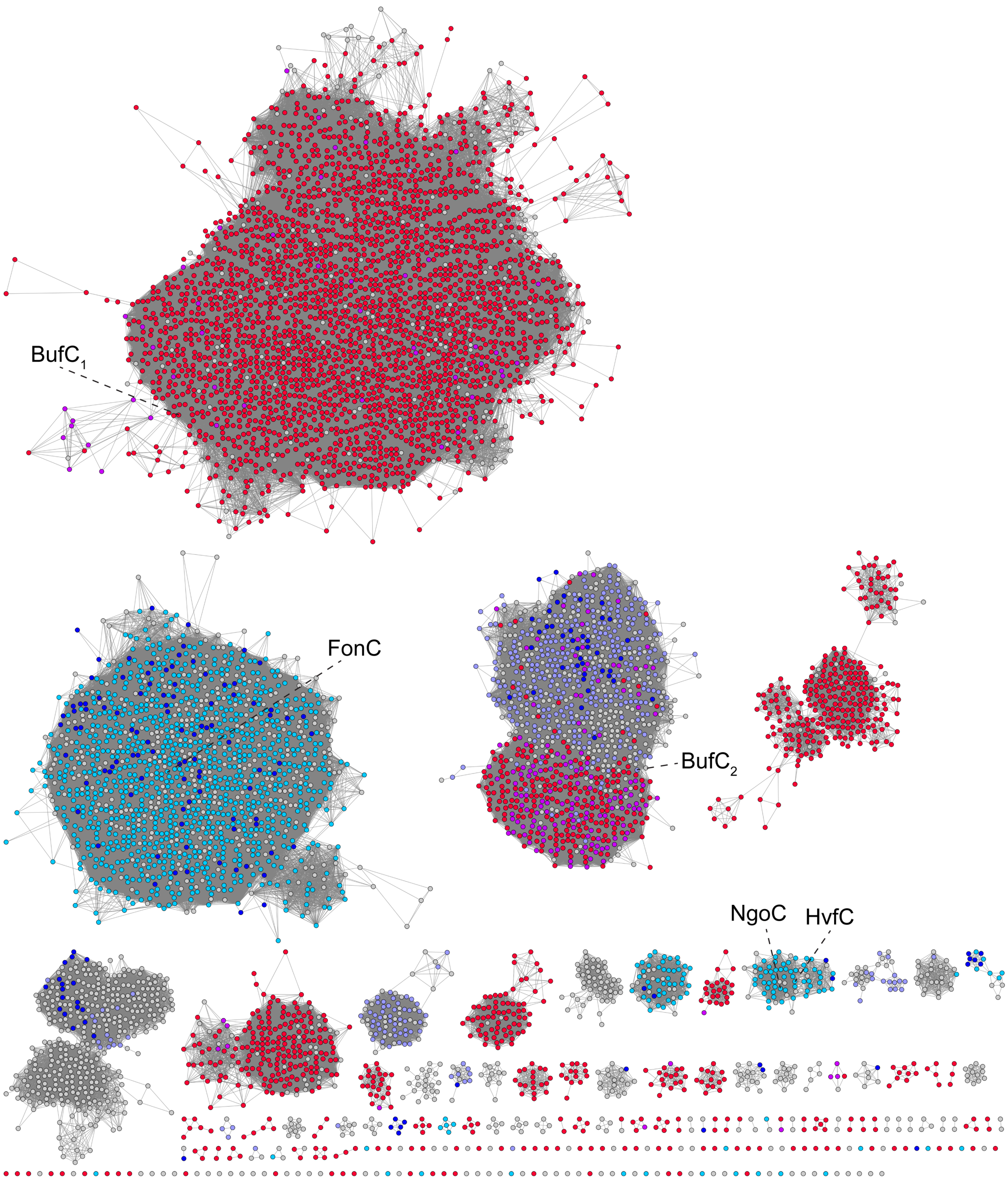


**Figure S30 (cont.). Biosynthetic protein sequence similarity networks colored by precursor peptide family.**

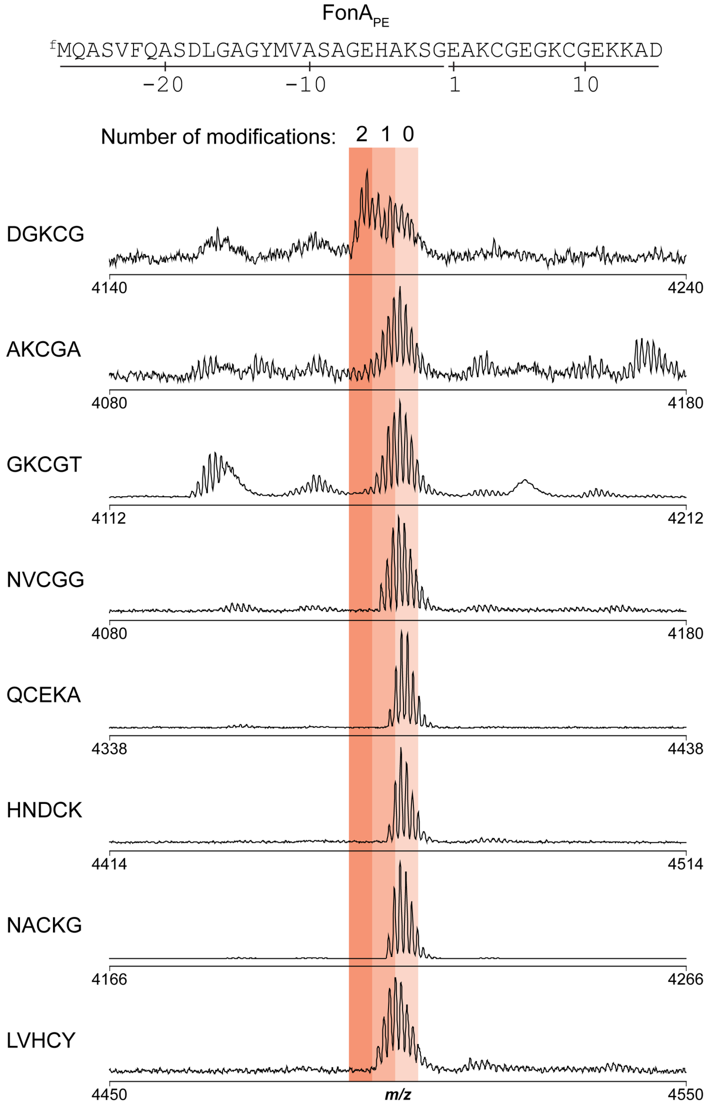


**Figure S31. FonB substrate tolerance for naturally occurring 5-mer repeats.** MALDI-TOF-MS spectra of FonBC reactions on FonA$_{PE}$ variants with the repeat regions altered to match other naturally occurring precursor peptides. For example, mutant FonA$_{PE}$DGKCG substitutes DGKCG both repeats of FonA$_{PE}$, yielding DGKCGDGKCG for core residues 1-10.

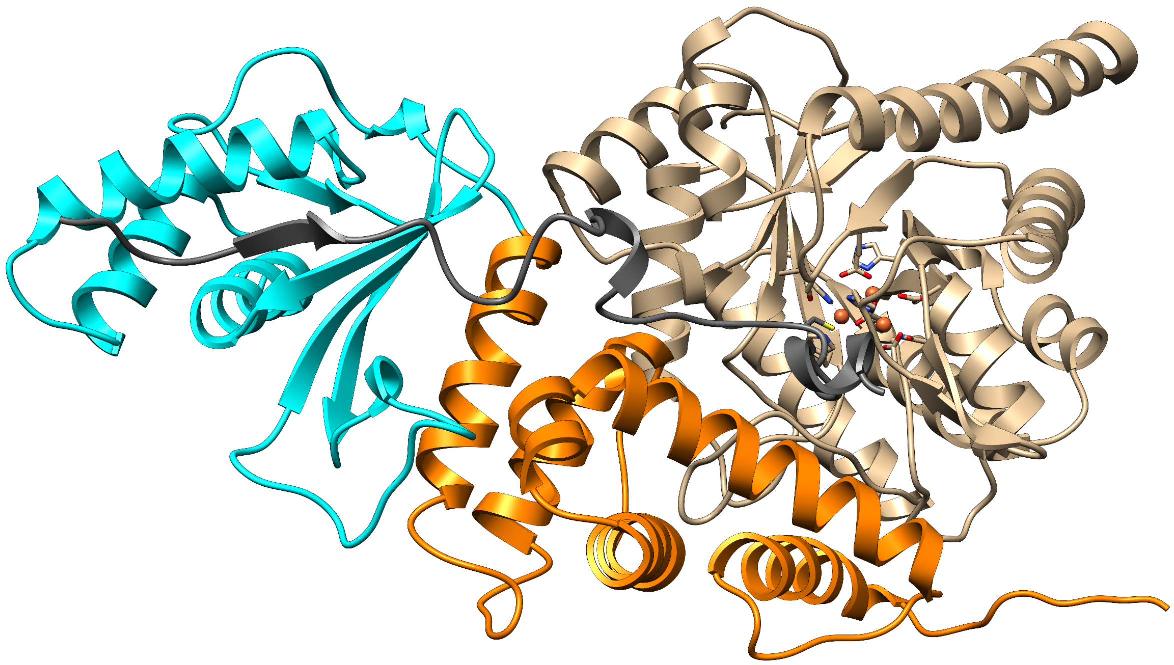

**Figure S32. AlphaFold model for FonA$_{PE}$BC.** Model was generated using AlphaFold 3.[15] Three Fe atoms were also added during prediction. Color annotations: Gray, FonA$_{PE}$; Tan, FonB; Orange, FonC DUF2063 domain; Cyan, FonC RRE domain.

**a**

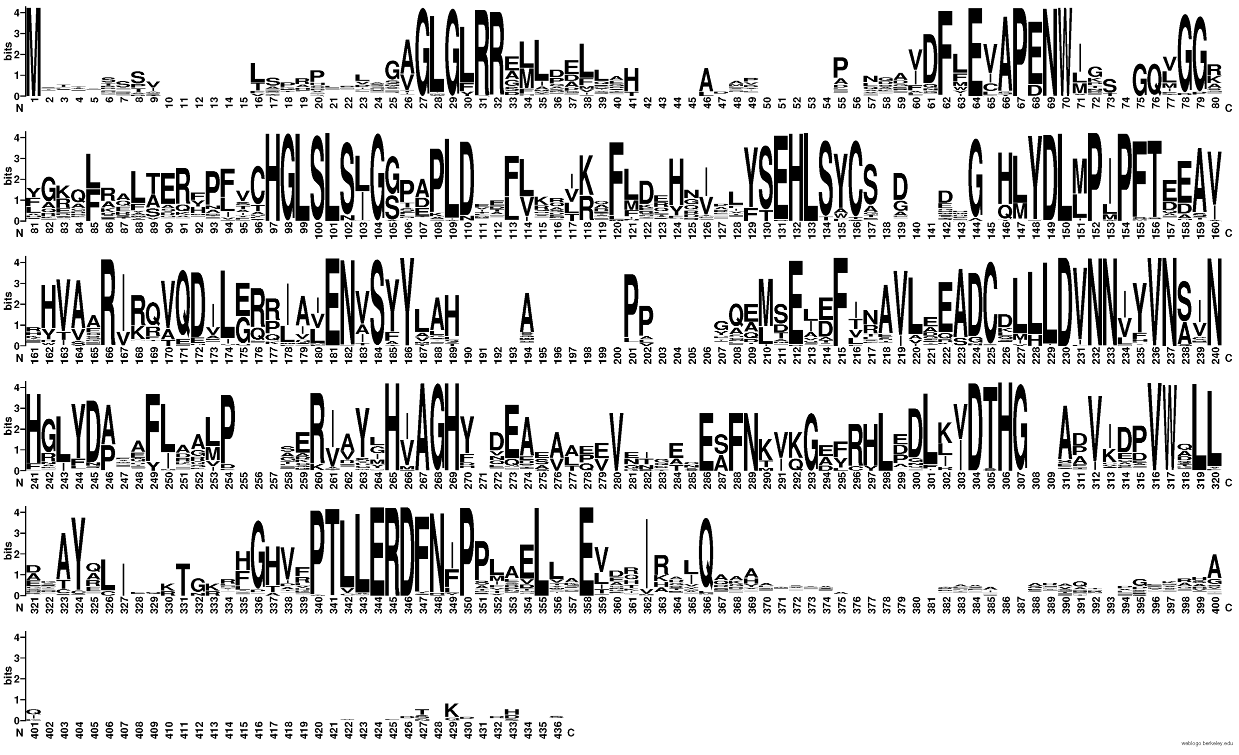

**Figure S33. MNIO sequence logos by substrate family. a**, Sequence logo (*n* = 100) for MNIO enzymes predicted to modify Lys-Cys family repeat sequences. **b**, Sequence logo (*n* = 100) for MNIO enzymes predicted to modify Cys-Lys family repeat sequences. **c**, Sequence logo (*n* = 100) for MNIO enzymes predicted to modify DUF2282 substrates.

b

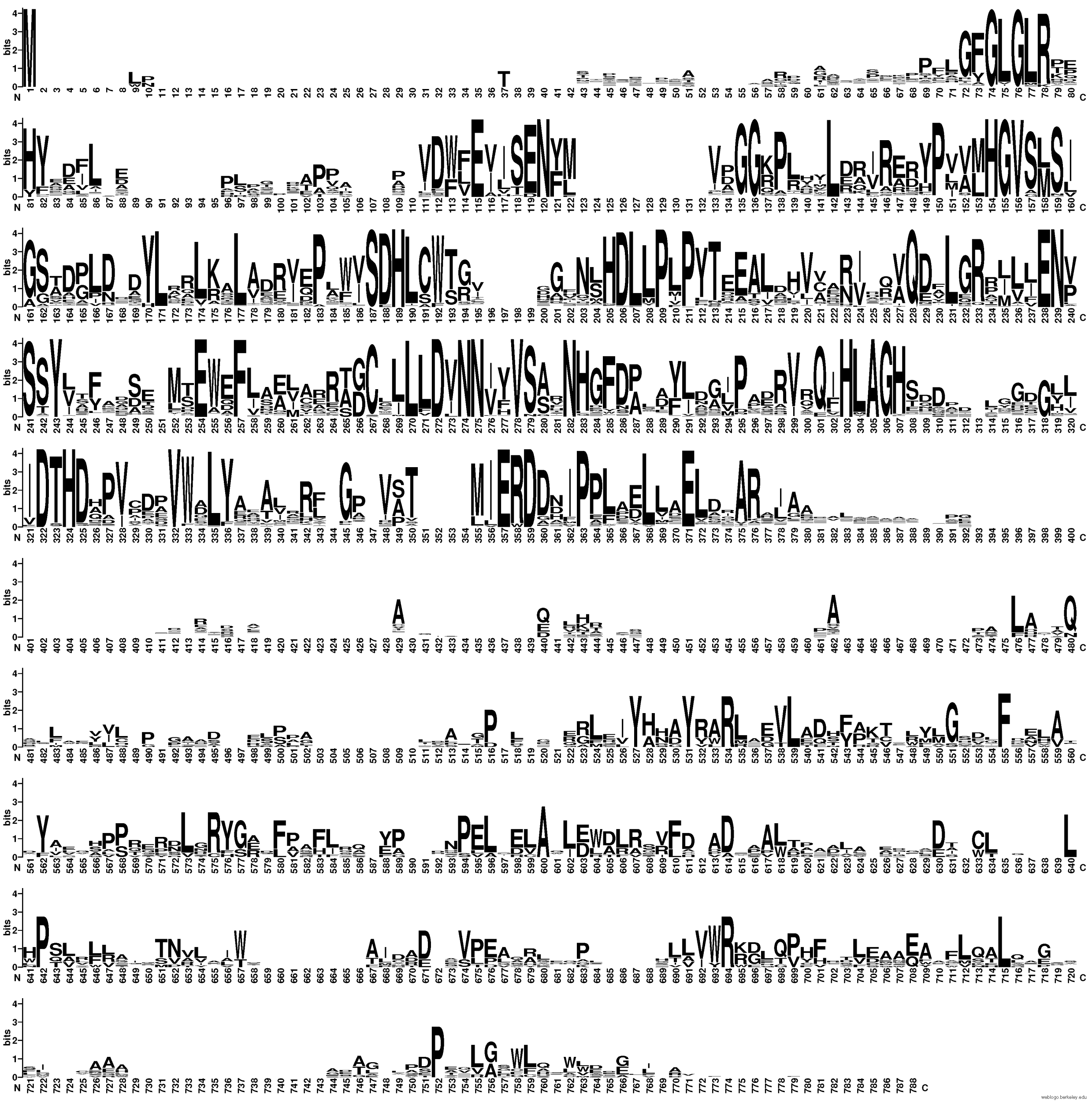

**Figure S33 (cont.). MNIO sequence logos by substrate family.**

c

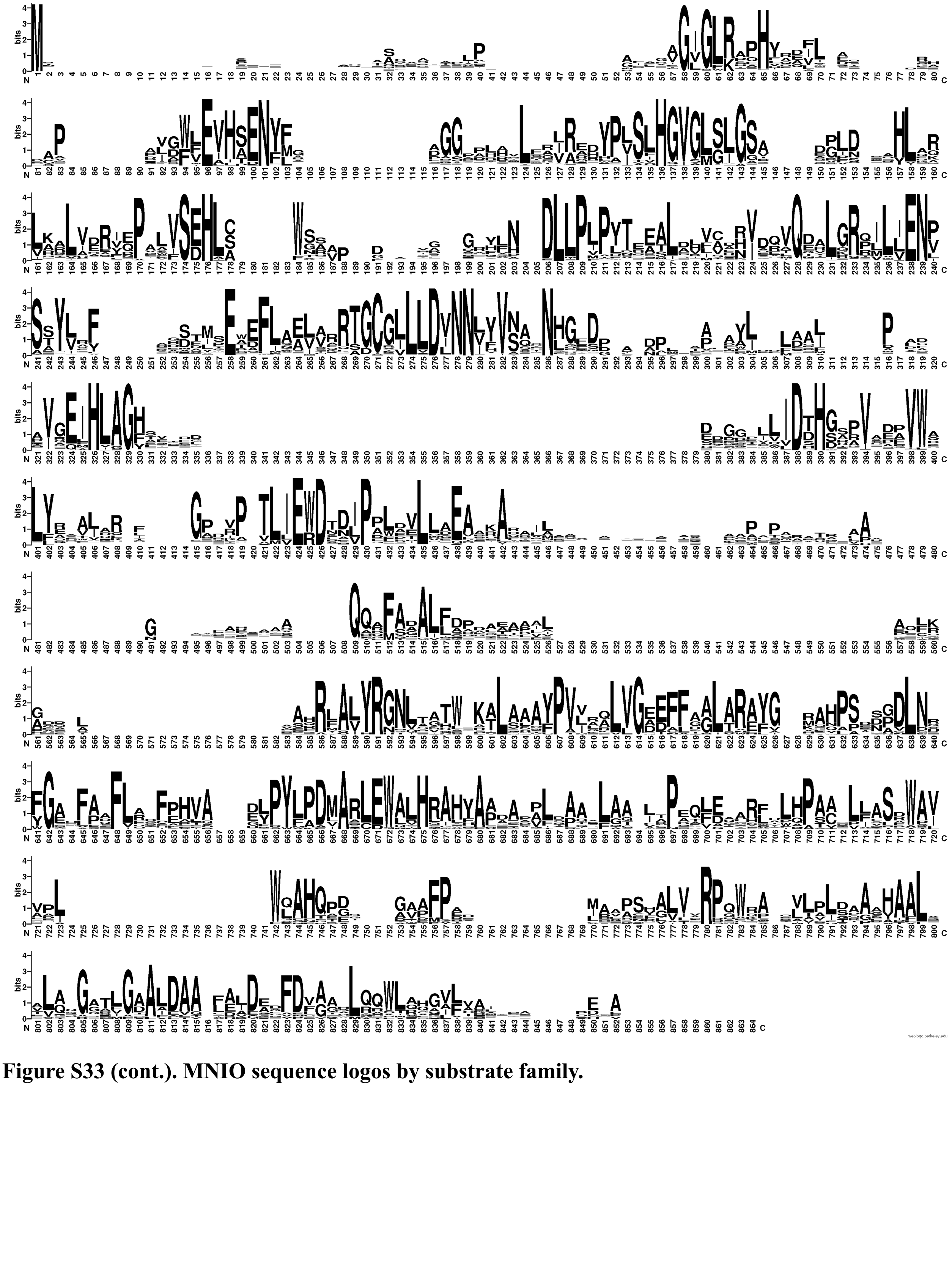

**Figure S33 (cont.). MNIO sequence logos by substrate family.**

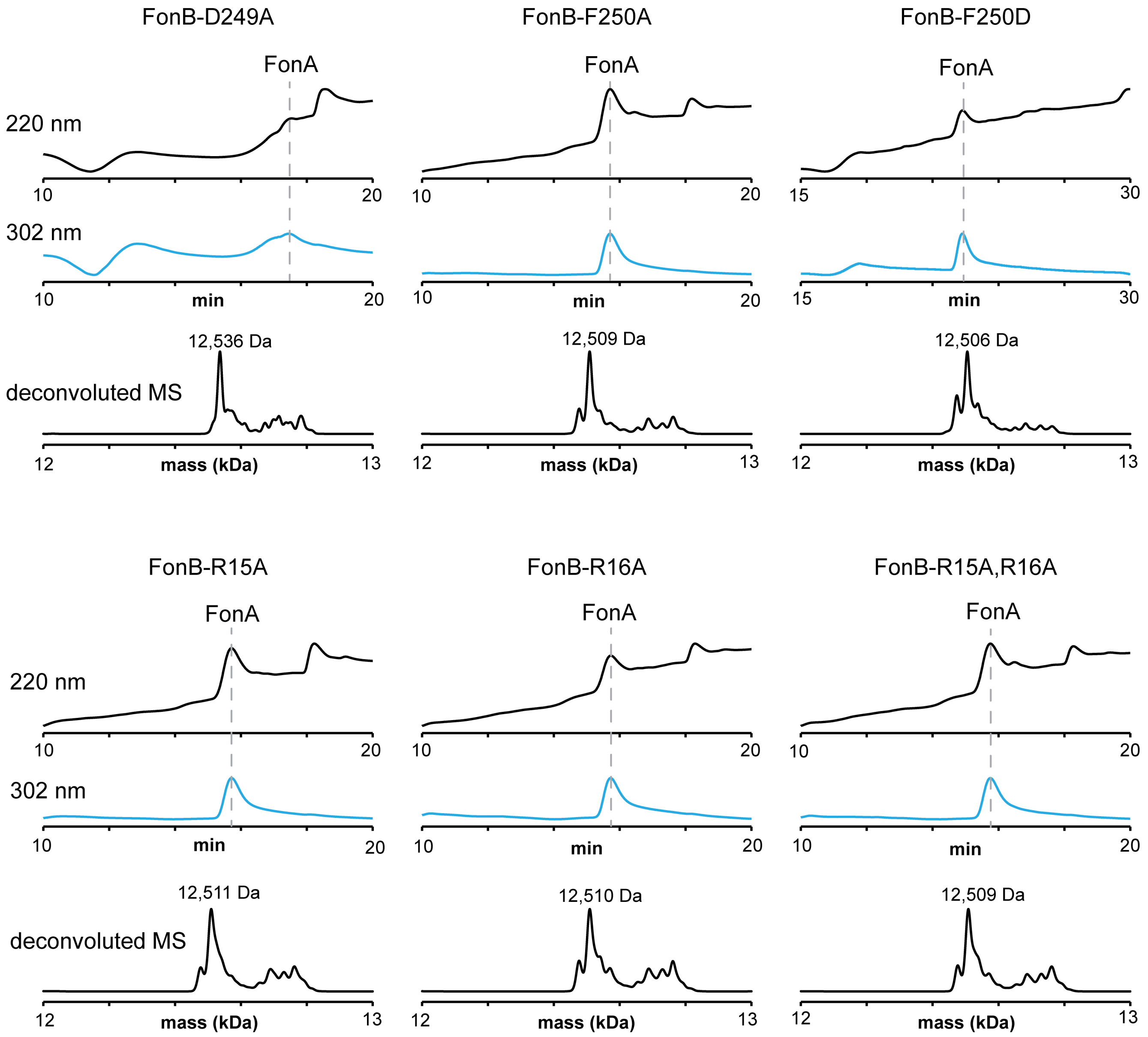


**Figure S34. FonB variant heterologous co-expressions.** UV chromatograms and deconvoluted mass spectra of FonA peptides upon co-expression with FonB variants and FonC.

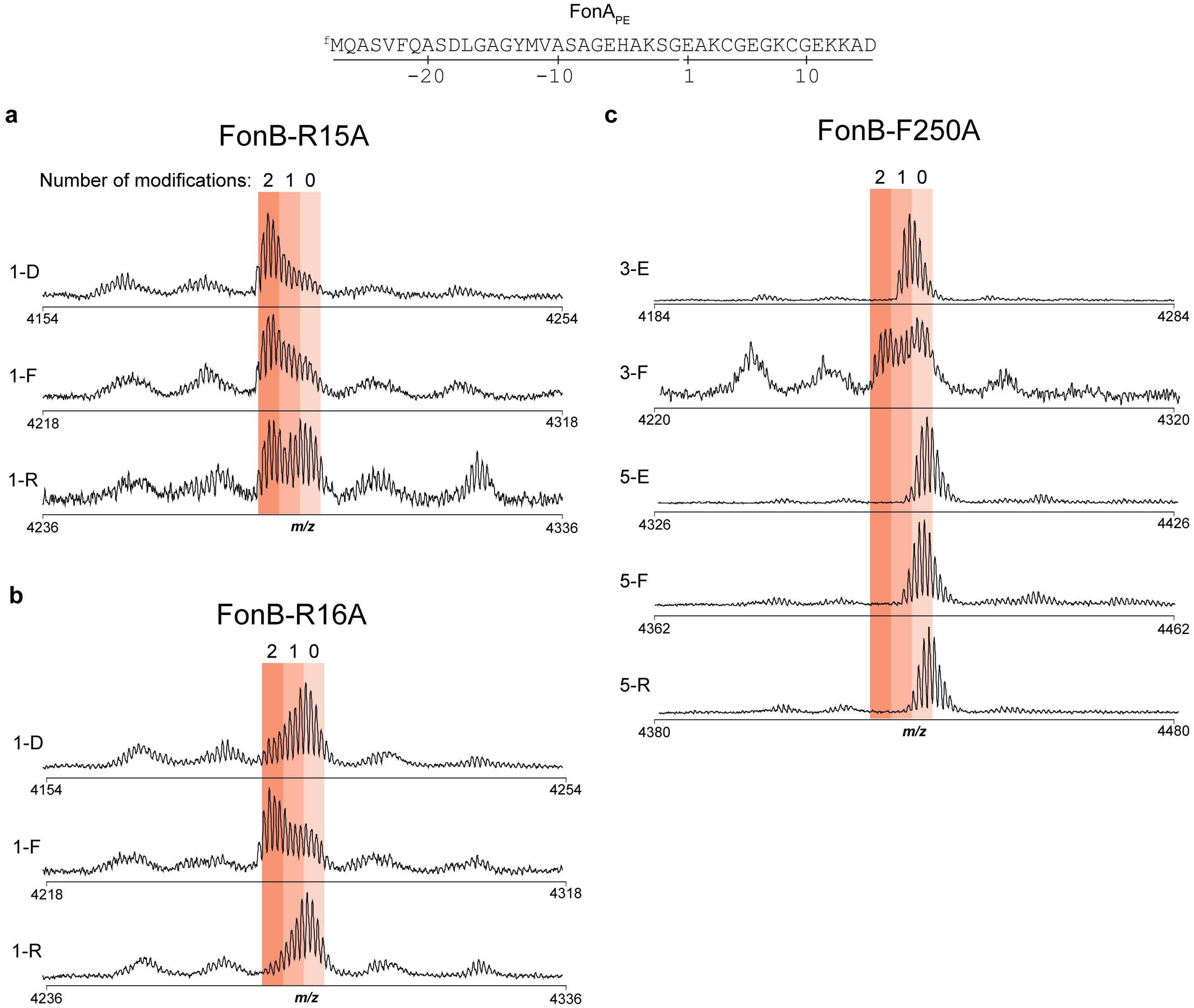


**Figure S35. *In vitro* activity of FonB Ala variants.** MALDI-TOF-MS spectra of reactions containing combinations of FonA$_{PE}$ variants, FonC, and FonB Ala variants. **a**, FonB-R15A. **b**, FonB-R16A. **c**, FonB-F250A. For example, FonA$_{PE}$1-A substitutes Ala for residues 1 and 6 of FonA$_{PE}$.

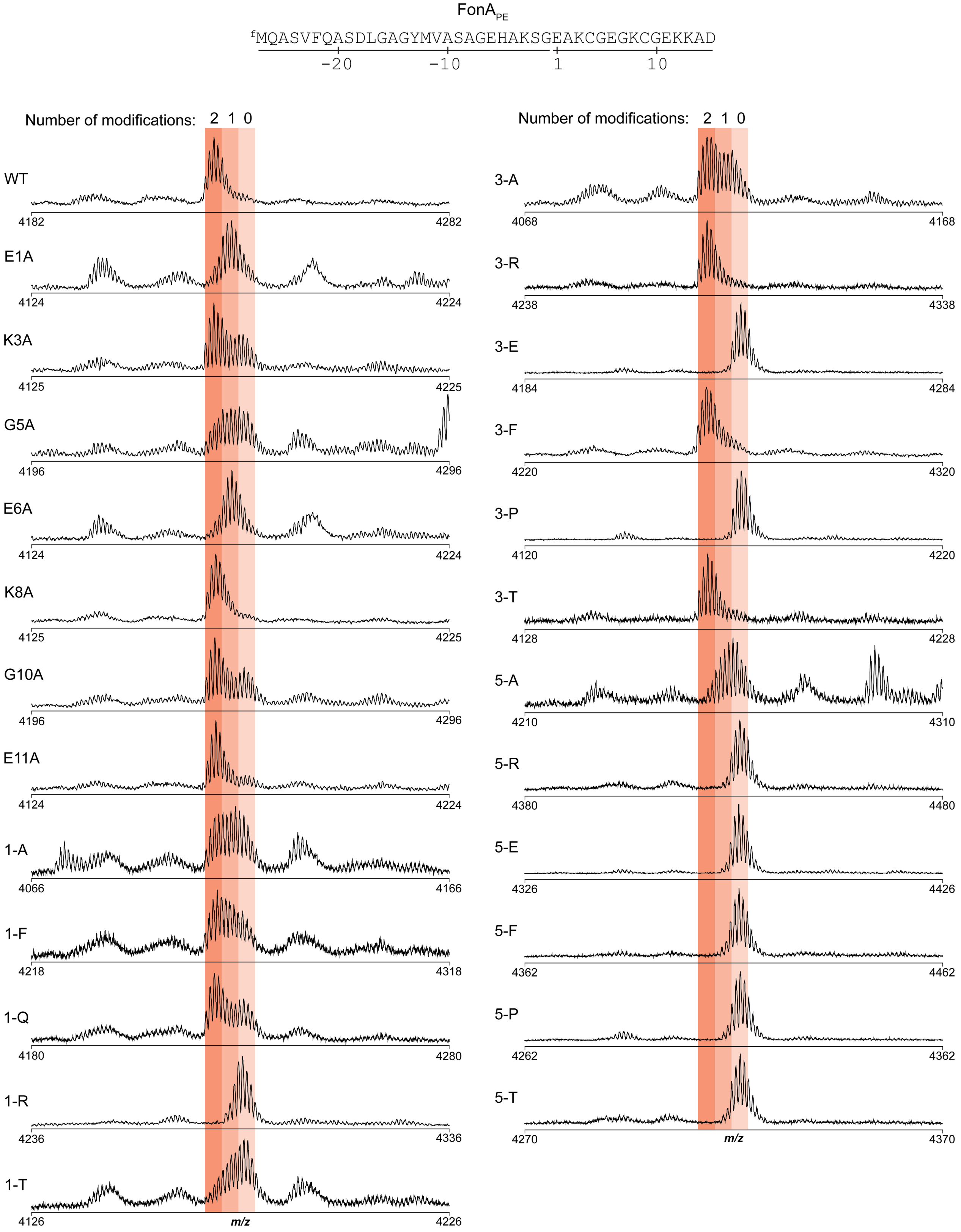


**Figure S36. FonB-F250D substrate tolerance.** MALDI-TOF-MS spectra of FonB-(F250D)C reactions on FonA$_{PE}$ variants. For example, FonA$_{PE}$1-A substitutes Ala for residues 1 and 6 of FonA$_{PE}$.

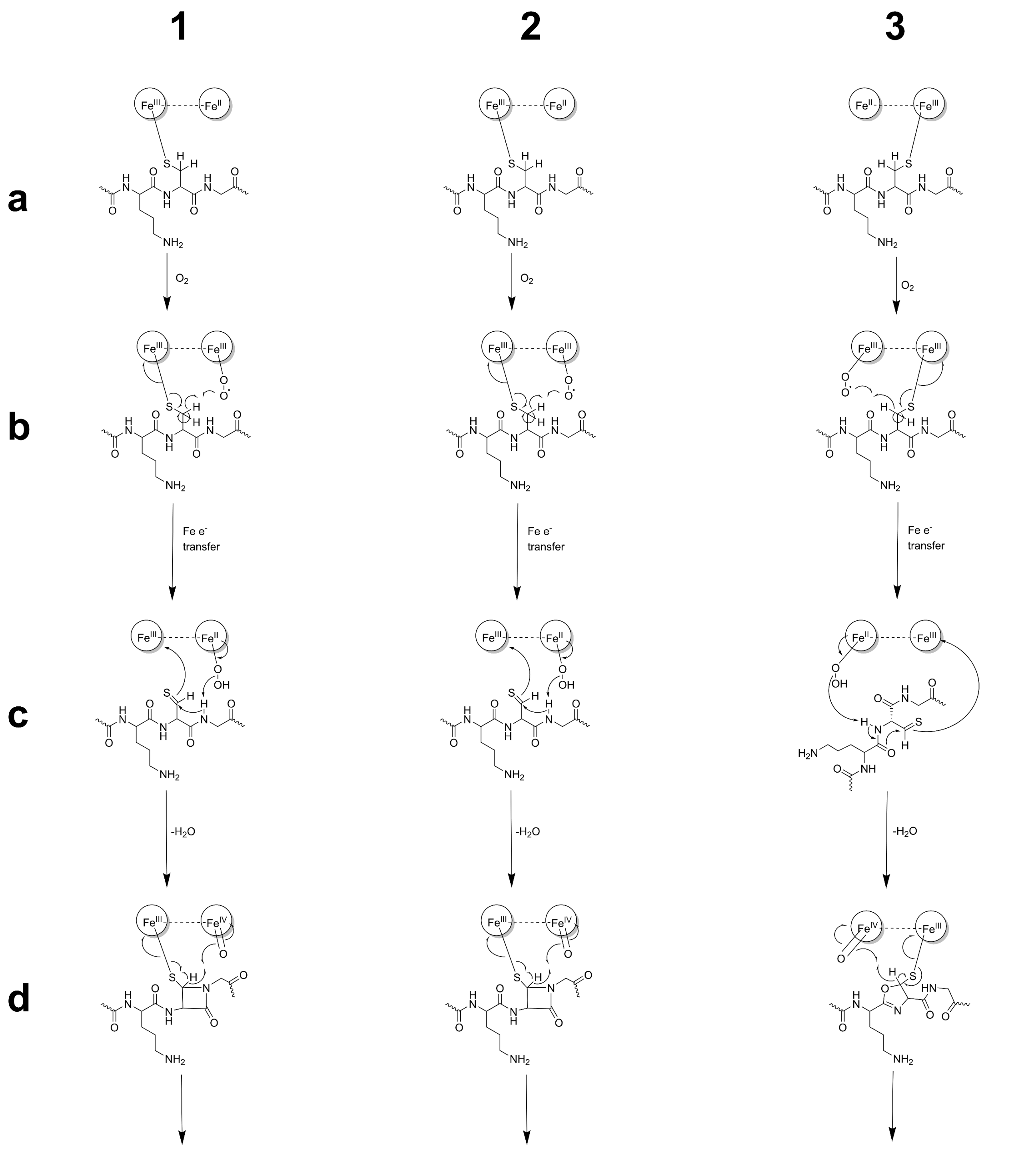

**Figure S37. Mechanistic proposals for 5-thiooxazole installation.** Proposed mechanisms for both oxazolone/thioamide (OxT, mechanism **1**) and 5-thiooxazole (5TO, mechanisms **2** and **3**) formation shown with various intermediates (**a-g**). In mechanisms **1** and **2**, differences in opening of the azetidinone intermediate **e** yield the respective OxT and 5TO structures. In mechanism **3,** the iron-superoxo species abstracts the N-terminal amide N-H rather than the C-terminal amide N-H, resulting in a five-membered ring intermediate, subsequently yielding 5TO.

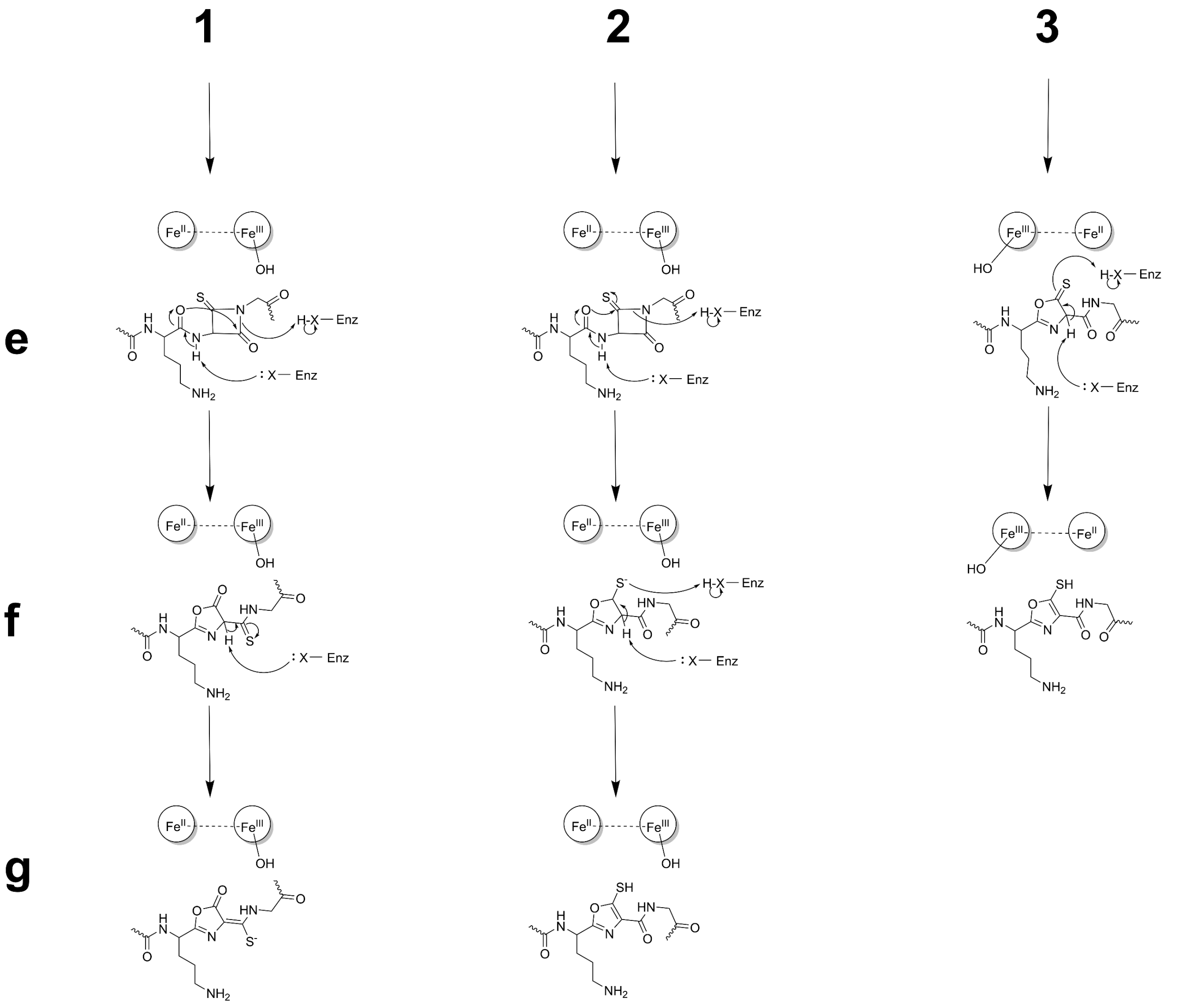


**Figure S37 (cont.). Mechanistic proposals for 5-thiooxazole installation.**

**Supplementary References.**

(10) Das, D.; Grishin, N. V.; Kumar, A.; Carlton, D.; Bakolitsa, C.; Miller, M. D.; Abdubek, P.; Astakhova, T.; Axelrod, H. L.; Burra, P.; Chen, C.; Chiu, H.-J.; Chiu, M.; Clayton, T.; Deller, M. C.; Duan, L.; Ellrott, K.; Ernst, D.; Farr, C. L.; Feuerhelm, J.; Grzechnik, A.; Grzechnik, S. K.; Grant, J. C.; Han, G. W.; Jaroszewski, L.; Jin, K. K.; Johnson, H. A.; Klock, H. E.; Knuth, M. W.; Kozbial, P.; Krishna, S. S.; Marciano, D.; McMullan, D.; Morse, A. T.; Nigoghossian, E.; Nopakun, A.; Okach, L.; Oommachen, S.; Paulsen, J.; Puckett, C.; Reyes, R.; Rife, C. L.; Sefcovic, N.; Tien, H. J.; Trame, C. B.; van den Bedem, H.; Weekes, D.; Wooten, T.; Xu, Q.; Hodgson, K. O.; Wooley, J.; Elsliger, M.-A.; Deacon, A. M.; Godzik, A.; Lesley, S. A.; Wilson, I. A. The Structure of the First Representative of Pfam Family PF09836 Reveals a Two-Domain Organization and Suggests Involvement in Transcriptional Regulation. *Acta Crystallograph. Sect. F Struct. Biol. Cryst. Commun.* **2010**, *66* (10), 1174–1181. https://doi.org/10.1107/S1744309109022672.

(11) Kenney, G. E.; Goering, A. W.; Ross, M. O.; DeHart, C. J.; Thomas, P. M.; Hoffman, B. M.; Kelleher, N. L.; Rosenzweig, A. C. Characterization of Methanobactin from Methylosinus Sp. LW4. *J. Am. Chem. Soc.* **2016**, *138* (35), 11124–11127. https://doi.org/10.1021/jacs.6b06821.

(12) Chioti, V. T.; Clark, K. A.; Ganley, J. G.; Han, E. J.; Seyedsayamdost, M. R. N–Cα Bond Cleavage Catalyzed by a Multinuclear Iron Oxygenase from a Divergent Methanobactin-like RiPP Gene Cluster. *J. Am. Chem. Soc.* **2024**, *146* (11), 7313–7323. https://doi.org/10.1021/jacs.3c11740.

(13) Manley, O. M.; Shriver, T. J.; Xu, T.; Melendrez, I. A.; Palacios, P.; Robson, S. A.; Guo, Y.; Kelleher, N. L.; Ziarek, J. J.; Rosenzweig, A. C. A Multi-Iron Enzyme Installs Copper-Binding Oxazolone/Thioamide Pairs on a Nontypeable Haemophilus Influenzae Virulence Factor. *Proc. Natl. Acad. Sci.* **2024**, *121* (28), e2408092121. https://doi.org/10.1073/pnas.2408092121.

(14) Leprevost, L.; Jünger, S.; Lippens, G.; Guillaume, C.; Sicoli, G.; Oliveira, L.; Falcone, E.; de Santis, E.; Rivera-Millot, A.; Billon, G.; Stellato, F.; Henry, C.; Antoine, R.; Zirah, S.; Dubiley, S.; Li, Y.; Jacob-Dubuisson, F. A Widespread Family of Ribosomal Peptide Metallophores Involved in Bacterial Adaptation to Metal Stress. *Proc. Natl. Acad. Sci.* **2024**, *121* (49), e2408304121. https://doi.org/10.1073/pnas.2408304121.

(15) Abramson, J.; Adler, J.; Dunger, J.; Evans, R.; Green, T.; Pritzel, A.; Ronneberger, O.; Willmore, L.; Ballard, A. J.; Bambrick, J.; Bodenstein, S. W.; Evans, D. A.; Hung, C.-C.; O'Neill, M.; Reiman, D.; Tunyasuvunakool, K.; Wu, Z.; Žemgulytė, A.; Arvaniti, E.; Beattie, C.; Bertolli, O.; Bridgland, A.; Cherepanov, A.; Congreve, M.; Cowen-Rivers, A. I.; Cowie, A.; Figurnov, M.; Fuchs, F. B.; Gladman, H.; Jain, R.; Khan, Y. A.; Low, C. M. R.; Perlin, K.; Potapenko, A.; Savy, P.; Singh, S.; Stecula, A.; Thillaisundaram, A.; Tong, C.; Yakneen, S.; Zhong, E. D.; Zielinski, M.; Žídek, A.; Bapst, V.; Kohli, P.; Jaderberg, M.; Hassabis, D.; Jumper, J. M. Accurate Structure Prediction of Biomolecular Interactions with AlphaFold 3. *Nature* **2024**, *630* (8016), 493–500. https://doi.org/10.1038/s41586-024-07487-w.
